# Differential effects of group III metabotropic glutamate receptors on spontaneous inhibitory synaptic currents in spine-innervating double bouquet and parvalbumin-expressing dendrite-targeting GABAergic interneurons in human neocortex

**DOI:** 10.1101/2022.03.05.483105

**Authors:** Istvan P. Lukacs, Ruggiero Francavilla, Martin Field, Emily Hunter, Michael Howarth, Sawa Horie, Puneet Plaha, Richard Stacey, Laurent Livermore, Olaf Ansorge, Gabor Tamas, Peter Somogyi

**Author notes:** Correspondence: Dr Istvan P. Lukacs or Prof Peter Somogyi, Department of Pharmacology, University of Oxford, Mansfield Road, Oxford, OX1 3QT. I.P.L., Institute of Experimental Medicine, 43 Szigony Street, Budapest, Hungary, 1083; of R.F., Centre de Recherche, CHU Sainte-Justine (CHUSJ), Montreal, QC, Canada, H3T 1C5; of M.H., Worldwide Clinical Trials, Beeston Business Park, Nottingham, UK, NG9 1LA; of S.H., Kawasaki Medical School, 577 Matsushima, Kurashiki, Okayama, Japan, 701-0192; of L.L., Department of Neurosurgery, Leeds General Infirmary, Great George Street, Leeds, UK, LS1 3EX.

## Abstract

Diverse neocortical GABAergic neurons specialise in synaptic targeting and their effects are modulated by presynaptic metabotropic glutamate receptors (mGluRs) suppressing neurotransmitter release in rodents, but their effects in human neocortex are unknown. We tested whether activation of group III mGluRs by L-AP4 changes GABA_A_ receptor-mediated spontaneous inhibitory postsynaptic currents (sIPSCs) in two distinct dendritic spine-innervating GABAergic interneurons recorded *in vitro* in human neocortex. Calbindin-positive *double bouquet cells* (DBC) had columnar “horsetail” axons descending through layers II-V innervating dendritic spines (48%) and shafts, but not somata of pyramidal and non-pyramidal neurons. *Parvalbumin-expressing dendrite-targeting cell* (PV-DTC) axons extended in all directions innervating dendritic spines (22%), shafts (65%) and somata (13%). As measured, 20% of GABAergic neuropil synapses innervate spines, hence DBCs, but not PV-DTCs, preferentially select spine targets. Group III mGluR activation paradoxically increased the frequency of sIPSCs in DBCs (to median 137% of baseline), but suppressed it in PV-DTCs (median 92%), leaving the amplitude unchanged. The facilitation of sIPSCs in DBCs may result from their unique GABAergic input being disinhibited via network effect. We conclude that dendritic spines receive specialised, diverse GABAergic inputs, and group III mGluRs differentially regulate GABAergic synaptic transmission to distinct GABAergic cell types in human cortex.

## Introduction

Dendritic spines receive the vast majority of synapses in the cerebral cortex and most of the presynaptic terminals release glutamate to them (Ramon y Cajal, 1888; Kasthuri et al., 2015; Cano-Astorga et al., 2021). Dendritic spines are highly plastic structures and constantly change their shape and size in response to synaptic activity (Maletic-Savatic et al., 1999; Holtmaat et al., 2006; Araya et al., 2014). Those connected to their parent dendritic shafts via narrow spine necks functionally isolate synaptic junctions from each other and allow independent changes in membrane potential (Kwon et al., 2017; Cornejo et al., 2021), receptor activity (Matsuzaki et al., 2001), signalling molecular mechanisms such as changes in calcium concentration (Yuste et al., 2000; Chen et al., 2011) and use-dependent changes in individual synaptic strengths (Matsuzaki et al., 2004; Lee et al., 2009). Most dendritic spines originate from pyramidal cells, in addition to other glutamatergic cortical neurons such as spiny stellate cells in layer IV, and receive a single synaptic input. With few exceptions, GABAergic neurons have fewer or no spines. A relatively small, but significant fraction of dendritic spines also receive an additional GABAergic synapse next to the glutamatergic one, as shown in rodents (Kubota et al., 2007), the cat (Beaulieu & Somogyi, 1990) and monkey (Beaulieu et al., 1992). Because of the vast volume density of dendritic spines, GABAergic synapses on spines are more than twice as frequent as on cell bodies in both the monkey and cat cortex (Beaulieu et al., 1992).

The relative isolation of dendritic spine inputs from the rest on the neuron suggests that glutamatergic and GABAergic synapses on a single spine have more influence on each other than on other inputs to the same neuron. It has been shown that spine-innervating GABAergic synapses can selectively attenuate somatically and synaptically evoked calcium influx into single dendritic spines (Chiu et al., 2013). However, how this influences synaptic plasticity is unknown. It is also unknown how such doubly innervated spines are distributed on the neuron, or if the spines receiving a GABAergic input are just a random subset or a functionally distinct population. That the latter may be the case is indicated by the finding that in the rat frontal cortex, spines receiving a thalamo-cortical glutamatergic input are more likely to be innervated also by a GABAergic synapse than other spines (Kubota et al., 2007). Thus, cortical GABAergic neurons, which might specialise in spine innervation could selectively regulate particular glutamatergic inputs to a given postsynaptic neuron.

Several distinct types of GABAergic cortical interneuron have been shown to innervate dendritic spines to differing degrees in addition to dendritic shafts and somata in rodents (Kawaguchi & Kubota, 1998; Kubota et al., 2007), the cat (Somogyi et al., 1983; Kisvárday et al., 1985; Tamás et al., 1997a) and the monkey (Somogyi & Cowey, 1981). There are clear molecular and some synaptic circuit homologies amongst the GABAergic cell types of the rodent and human cerebral cortex (Hodge et al., 2019; Bakken et al., 2021). However, with the exception of the GABAergic axo-axonic cell (Kisvárday et al., 1986), the basket cell (Szabadics et al., 2006) and the rosehip cell (Boldog et al., 2018) at present it is unknown to what degree distinct human GABAergic neuronal types dedicate their outputs selectively to functionally different subcellular domains of postsynaptic neurons. A potential compartmentalisation of the postsynaptic neuron surface by local GABAergic neurons has fundamental consequences for the regulation of multiple synaptic inputs to a given neuron and for the temporal dynamics of input to output transformation (Lovett-Barron et al., 2012; Somogyi et al., 2014; Doron et al., 2017).

In the context of synaptic plasticity and the GABAergic innervation of dendritic spines, a particular interest is the iconic double bouquet cell (DBC), which in the cat and monkey appears to innervate dendritic spines preferentially (Somogyi & Cowey, 1981; Tamás et al., 1997a) and forms a system or radial GABAergic projection through cortical layers (Somogyi et al., 1981). In the current study, the name “double bouquet cell” is used in a narrow sense for neurons with cell bodies in layers 2 and upper 3 and having descending ‘horse tail’-like narrow axonal bundles through all layers. The “horsetail” axon of DBCs in primates strongly resembles the regularly spaced radial axonal bundles revealed by immunoreactions to the calcium binding protein calbindin (CB), which has been used to study these cells (DeFelipe et al., 1989; del Río & DeFelipe, 1995; Peters & Sethares, 1997). The small diameter radial axon bundle has been the subject of speculation regarding its potential roles in ‘minicolumns’ (del Río & Defelipe, 1997b; DeFelipe et al., 2006), but the activity and cortical role of DBCs remains unknown.

Another iconic GABAergic cell type is the ‘basket cell’, so named because it is assumed that its terminals form perisomatic baskets that are thought to have a large inhibitory effect on the output of the postsynaptic neurons. Basket cells, including parvalbumin-expressing (PV+) cells, show great diversity in molecular expression (Tasic et al., 2018; Hodge et al., 2019; Gouwens et al., 2020; Bakken et al., 2021), lateral spread and interlaminar specificity of their axons (Somogyi et al., 1983; DeFelipe et al., 1986; Kisvárday et al., 1987), and their synaptic output to different postsynaptic targets on the same cell, including dendritic spines (Somogyi et al., 1983; Somogyi & Soltész, 1986; Kawaguchi & Kubota, 1998; Szabadics et al., 2006). In the human cerebral cortex the synaptic output selectivity, if any, of PV+ basket cells is unknown.

In the present study on the human cerebral cortex, we explored and compared the evoked firing behaviour and synaptic connectivity of DBCs and putative basket cells in surgically resected samples, which were removed from patients for the treatment of tumours or temporal lobe epilepsy. We have also tested if synaptic inputs to the two distinct cell types differed in their regulation by glutamate receptors. We chose to study group III metabotropic glutamate receptors (mGluRs), because of previous indications from studies on rodents that group III mGluRs are differentially expressed in cortical GABAergic neurons (Dalezios et al., 2002; Somogyi et al., 2003) and selectively regulate GABAergic synaptic transmission (Semyanov & Kullmann, 2000; Kogo et al., 2004; Klar et al., 2015), and because of the large investment into developing drugs acting on group III mGluRs for the treatment of neurological and psychiatric conditions (Maksymetz et al., 2017; Charvin, 2018; Ferraguti, 2018). The results presented here, some of which have been published in abstract form (Lukacs et al., 2019, 2020), reveal a high degree of multidimensional specialisation of human GABAergic interneurons.

## Methods

### Ethical approval and patient consent

Human tissue samples were collected from patients undergoing neurosurgery at the John Radcliffe Hospital (Oxford) for the treatment of brain tumours or temporal lobe epilepsy (table 2) in accordance with the Human tissue Act 2004 (UK), under the ethical licence (15/SC/0639) of the Oxford Brain Bank (OBB), Department of Neuropathology, John Radcliffe Hospital, Oxford, UK. Patients consented to providing the samples after they were fully informed by a medical professional. All collected samples represented “access tissue”, that is tissue necessarily removed in order to access the diseased part of the brain, but not or only partially affected by the pathological process, which otherwise would have been discarded. An additional three tissue samples (see table 2) were imported through the OBB from the laboratory of professor Gábor Tamás, at the University of Szeged, Hungary, which were collected after informed consent of patients in the Department of Neurosurgery, University of Szeged, Hungary, under the ethical licence (75/2004), as specified by Hungarian law and the University of Szeged.

**Table 1.** Primary antibody information

| molecule | host species | antigen | clonality | internal ref. | dilution | stock protein concentration (mg/ml) |
| --- | --- | --- | --- | --- | --- | --- |
| calbindin | rabbit | recombinant rat calbindin D-28k | polyclonal | 989 | 1:5000 | antiserum |
| calbindin | goat | mouse calbindin D-28k (NM009788) expressed in bacteria | polyclonal | 1427 | 1:500 | 200 |
| cannabinoid type 1 receptor (CB1) | guinea-pig | mouse CB1, C-terminal 31 aa (NM007726) | polyclonal | 1274 | 1:5000 | 200 |
| CB1 | rabbit | synthetic peptide aa residues 443-473, mouse and human | polyclonal | 1558 | 1:10000 | 10000 |
| cholecystokinin (CCK) | mouse | gastrin-17 | monoclonal | 1052 | 1:1000 | 5000 |
| calretinin (CR) | goat | human recombinant calretinin | polyclonal | 1116 | 1:1000 | antiserum |
| hexaribonucleotid binding protein-3 (NeuN) | mouse |  | monoclonal | 842 | 1:1000 | 1000 |
| parvalbumin (PV) | guinea-pig | rat recombinant protein aa 1-110 | polyclonal | 1310 | 1:1000-1:5000 | antiserum |
| PV | goat | rat muscle parvalbumin | polyclonal | 1258 | 1:1000 | antiserum |
| PV | mouse | purified carp II parvalbumin | monoclonal | 922 | 1:1000 | ascites |
| somatostatin (SM) | sheep | somatostatin conjugated to carrier protein | polyclonal | 1085 | 1:500-1:5000 | 5000 |
| SM | mouse | somatostatin-14 conjugated to carrier protein | monoclonal | 1276 | 1:400 | 140 |
| vesicular GABA transporter (VGAT) | guinea-pig | rat recombinant protein aa 2-115 | polyclonal | 1321 | 1:500-1:1000 | antiserum |
| vasoactive intestinal polypeptide (VIP) | rabbit | porcine VIP coupled to bovine thyroglobulin with carbodiimide linker | polyclonal | 1523 | 1:5000 | antiserum |
| VIP | mouse | VIP | monoclonal | 1053 | 1:25000 | antiserum |
| γ-aminobutyric acid (GABA) | rabbit | GABA-bovine serum albumin conjugate | polyclonal | 9 | 1:1000 | antiserum |

Table 1. Continued
| molecule | host species | source | product code | specificity reference |
| --- | --- | --- | --- | --- |
| calbindin | rabbit | SWANT | CB-38 | (Airaksinen et al., 1997) |
| calbindin | goat | Frontier Institute | Calbindin-Go-Af1040 | (Nakagawa et al., 1998) |
| cannabinoid type 1 receptor (CB1) | guinea-pig | Frontiers Institute | CB1-GP-Af530 | (Fukudome et al., 2004) |
| CB1 | rabbit | Immunogenes | Anti-CB1 (Rabbit) Polyclonal Antibody | (Dudok et al., 2015) |
| cholecystokinin (CCK) | mouse | CURE/Digestive Diseases research Center, Antibody/RIA Core, NIH Grant #DK41301 | 9303 | (Ohning et al., 1996; Geerling et al., 2006) |
| calretinin (CR) | goat | SWANT | CG1 | (Schiffmann et al., 1999) |
| hexaribonucleotid binding protein-3 (NeuN) | mouse | Chemicon International | MAB377 | (Tippett et al., 2007) |
| parvalbumin (PV) | guinea-pig | Synaptic Systems | 195 004 | synaptic systems |
| PV | goat | SWANT | PVG-213 | (Schwaller et al., 1999) |
| PV | mouse | SWANT | 235 | (Celio et al., 1988) |
| somatostatin (SM) | sheep | Cortex Biochem | CR2056SP |  |
| SM | mouse | Genetex | GTX71935 | (Kubota et al., 2011) |
| vesicular GABA transporter (VGAT) | guinea-pig | Synaptic Systems | 131 004 | (Ramer, 2008) |
| vasoactive intestinal polypeptide (VIP) | rabbit | Immunostar | 20077 | Immunostar |
| VIP | mouse | CURE/Digestive Diseases research Center, Antibody/RIA Core, NIH Grant #DK41301 | 55 | (Wong et al., 1996) |
| GABA | rabbit | produced by Prof. Somogyi | n.a. | (Hodgson et al., 1985) |

**Table 2.**
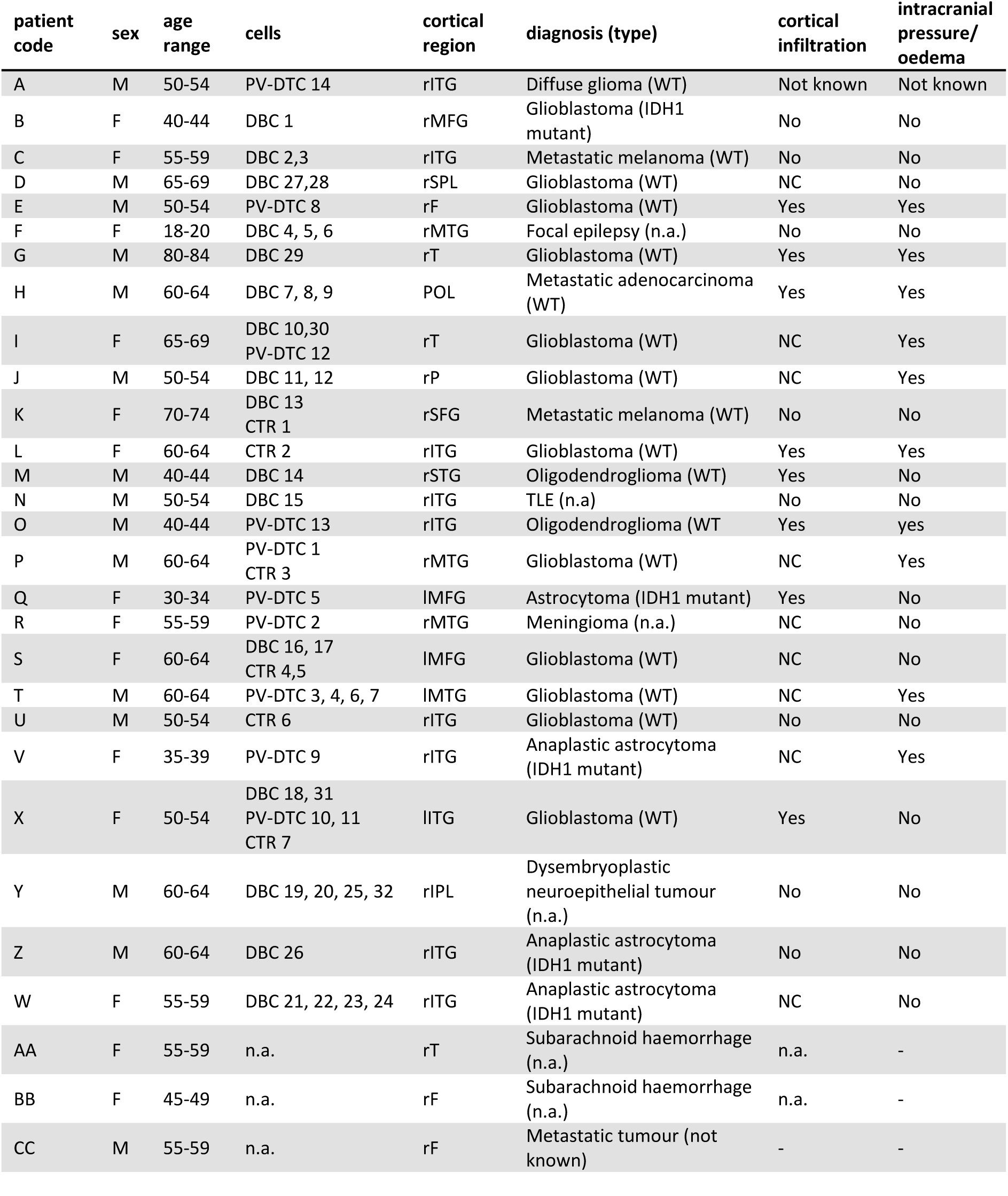
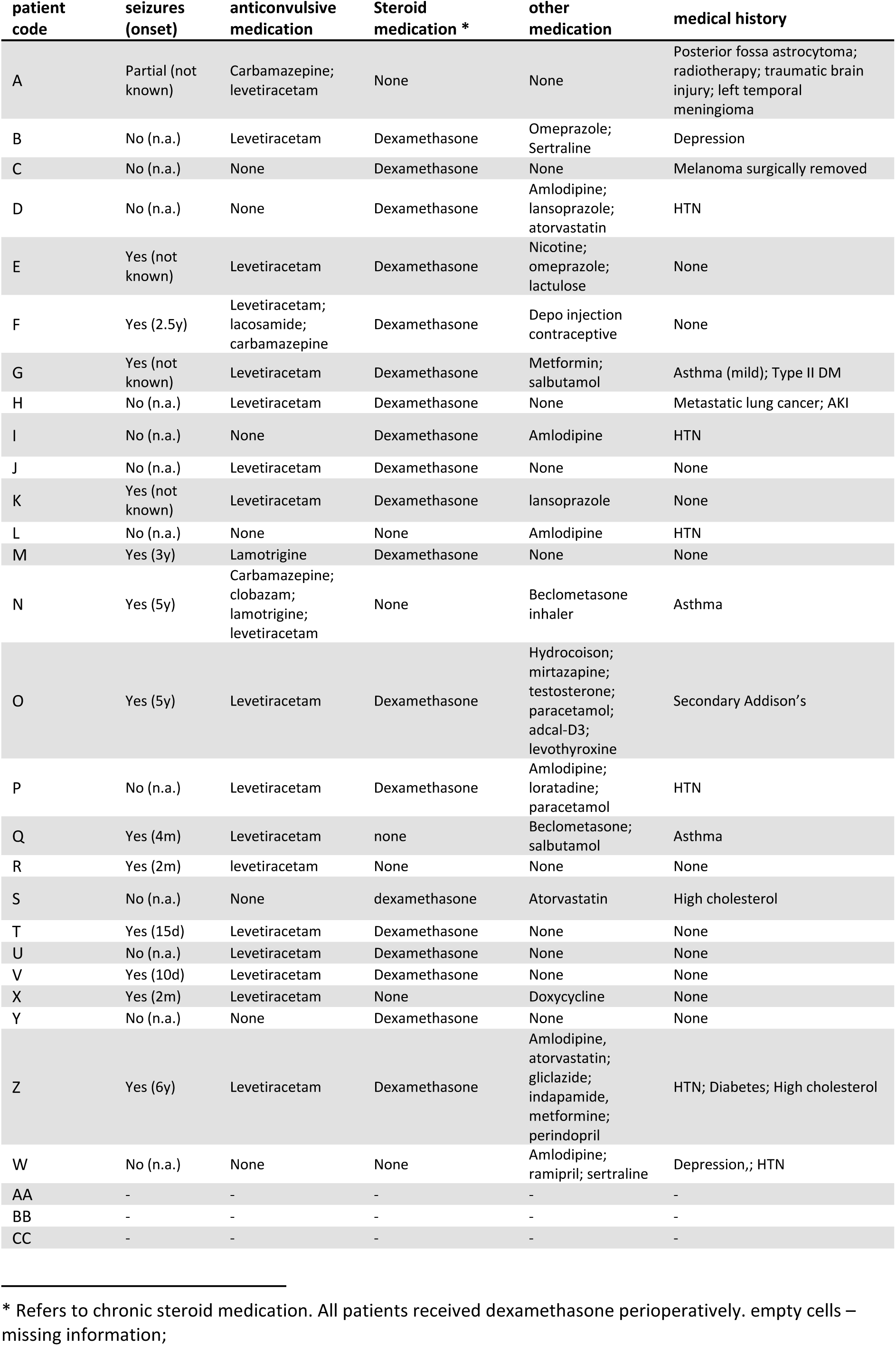
Patient information

| patient code | sex | age range | cells | cortical region | diagnosis (type) | cortical infiltration | intracranial pressure/oedema |
| --- | --- | --- | --- | --- | --- | --- | --- |
| A | M | 50-54 | PV-DTC 14 | rITG | Diffuse glioma (WT) | Not known | Not known |
| B | F | 40-44 | DBC 1 | rMFG | Glioblastoma (IDH1 mutant) | No | No |
| C | F | 55-59 | DBC 2,3 | rITG | Metastatic melanoma (WT) | No | No |
| D | M | 65-69 | DBC 27,28 | rSPL | Glioblastoma (WT) | NC | No |
| E | M | 50-54 | PV-DTC 8 | rF | Glioblastoma (WT) | Yes | Yes |
| F | F | 18-20 | DBC 4, 5, 6 | rMTG | Focal epilepsy (n.a.) | No | No |
| G | M | 80-84 | DBC 29 | rT | Glioblastoma (WT) | Yes | Yes |
| H | M | 60-64 | DBC 7, 8, 9 | POL | Metastatic adenocarcinoma (WT) | Yes | Yes |
| I | F | 65-69 | DBC 10,30<br>PV-DTC 12 | rT | Glioblastoma (WT) | NC | Yes |
| J | M | 50-54 | DBC 11, 12 | rP | Glioblastoma (WT) | NC | Yes |
| K | F | 70-74 | DBC 13<br>CTR 1 | rSFG | Metastatic melanoma (WT) | No | No |
| L | F | 60-64 | CTR 2 | rITG | Glioblastoma (WT) | Yes | Yes |
| M | M | 40-44 | DBC 14 | rSTG | Oligodendroglioma (WT) | Yes | No |
| N | M | 50-54 | DBC 15 | rITG | TLE (n.a.) | No | No |
| O | M | 40-44 | PV-DTC 13 | rITG | Oligodendroglioma (WT) | Yes | yes |
| P | M | 60-64 | PV-DTC 1<br>CTR 3 | rMTG | Glioblastoma (WT) | NC | Yes |
| Q | F | 30-34 | PV-DTC 5 | lMFG | Astrocytoma (IDH1 mutant) | Yes | No |
| R | F | 55-59 | PV-DTC 2 | rMTG | Meningioma (n.a.) | NC | No |
| S | F | 60-64 | DBC 16, 17<br>CTR 4,5 | lMFG | Glioblastoma (WT) | NC | No |
| T | M | 60-64 | PV-DTC 3, 4, 6, 7 | lMTG | Glioblastoma (WT) | NC | Yes |
| U | M | 50-54 | CTR 6 | rITG | Glioblastoma (WT) | No | No |
| V | F | 35-39 | PV-DTC 9 | rITG | Anaplastic astrocytoma (IDH1 mutant) | NC | Yes |
| X | F | 50-54 | DBC 18, 31<br>PV-DTC 10, 11<br>CTR 7 | lITG | Glioblastoma (WT) | Yes | No |
| Y | M | 60-64 | DBC 19, 20, 25, 32 | rIPL | Dysembryoplastic neuroepithelial tumour (n.a.) | No | No |
| Z | M | 60-64 | DBC 26 | rITG | Anaplastic astrocytoma (IDH1 mutant) | No | No |
| W | F | 55-59 | DBC 21, 22, 23, 24 | rITG | Anaplastic astrocytoma (IDH1 mutant) | NC | No |
| AA | F | 55-59 | n.a. | rT | Subarachnoid haemorrhage (n.a.) | n.a. | - |
| BB | F | 45-49 | n.a. | rF | Subarachnoid haemorrhage (n.a.) | n.a. | - |
| CC | M | 55-59 | n.a. | rF | Metastatic tumour (not known) | - | - |

Table 2. Continued
| patient code | seizures (onset) | anticonvulsive medication | Steroid medication * | other medication | medical history |
| --- | --- | --- | --- | --- | --- |
| A | Partial (not known) | Carbamazepine; levetiracetam | None | None | Posterior fossa astrocytoma; radiotherapy; traumatic brain injury; left temporal meningioma |
| B | No (n.a.) | Levetiracetam | Dexamethasone | Omeprazole; Sertraline | Depression |
| C | No (n.a.) | None | Dexamethasone | None | Melanoma surgically removed |
| D | No (n.a.) | None | Dexamethasone | Amlodipine; lansoprazole; atorvastatin | HTN |
| E | Yes (not known) | Levetiracetam | Dexamethasone | Nicotine; omeprazole; lactulose | None |
| F | Yes (2.5y) | Levetiracetam; lacosamide; carbamazepine | Dexamethasone | Depo injection contraceptive | None |
| G | Yes (not known) | Levetiracetam | Dexamethasone | Metformin; salbutamol | Asthma (mild); Type II DM |
| H | No (n.a.) | Levetiracetam | Dexamethasone | None | Metastatic lung cancer; AKI |
| I | No (n.a.) | None | Dexamethasone | Amlodipine | HTN |
| J | No (n.a.) | Levetiracetam | Dexamethasone | None | None |
| K | Yes (not known) | Levetiracetam | Dexamethasone | lansoprazole | None |
| L | No (n.a.) | None | None | Amlodipine | HTN |
| M | Yes (3y) | Lamotrigine | Dexamethasone | None | None |
| N | Yes (5y) | Carbamazepine; clobazam; lamotrigine; levetiracetam | None | Beclometasone inhaler | Asthma |
| O | Yes (5y) | Levetiracetam | Dexamethasone | Hydrocortisone; mirtazapine; testosterone; paracetamol; adcal-D3; levothyroxine | Secondary Addison's |
| P | No (n.a.) | Levetiracetam | Dexamethasone | Amlodipine; loratadine; paracetamol | HTN |
| Q | Yes (4m) | Levetiracetam | none | Beclometasone; salbutamol | Asthma |
| R | Yes (2m) | levetiracetam | None | None | None |
| S | No (n.a.) | None | dexamethasone | Atorvastatin | High cholesterol |
| T | Yes (15d) | Levetiracetam | Dexamethasone | None | None |
| U | No (n.a.) | Levetiracetam | Dexamethasone | None | None |
| V | Yes (10d) | Levetiracetam | Dexamethasone | None | None |
| X | Yes (2m) | Levetiracetam | None | Doxycycline | None |
| Y | No (n.a.) | None | Dexamethasone | None | None |
| Z | Yes (6y) | Levetiracetam | Dexamethasone | Amlodipine, atorvastatin; gliclazide; indapamide, metformine; perindopril | HTN; Diabetes; High cholesterol |
| W | No (n.a.) | None | None | Amlodipine; ramipril; sertraline | Depression;; HTN |
| AA | - | - | - | - | - |
| BB | - | - | - | - | - |
| CC | - | - | - | - | - |

\* Refers to chronic steroid medication. All patients received dexamethasone perioperatively. empty cells – missing information;

### Sample collection and slice preparation

Samples were collected between 10 am and 5 pm (UK time) as described previously (Bocchio et al., 2019; Field et al., 2021). A small (0.5–2 cm^3^) block of neocortex was dropped into cold cutting artificial cerebrospinal fluid (ACSF) saturated with carbogen (95 % O_2_, 5 % CO_2_), and divided into smaller blocks as necessary. The cutting ACSF contained the following (in mM): 65 sucrose, 85 NaCl, 25 NaHCO_3_, 2.5 KCl, 1.25 NaH_2_PO_4_, 0.5 CaCl_2_, 7 MgCl_2_, 10 glucose (pH ∼7.3; ∼300 mOsm/L) for samples A-C and AA-CC (see table 2); or the following (in mM): 92 N-methyl-D-glucamine (NMDG), 2.5 KCl, 1.25 NaH2PO4, 30 NaHCO_3_, 20 4-(2-hydroxyethyl)-1-piperazineethanesulfonic acid (HEPES), 25 glucose, 2 thiourea, 5 Na-ascorbate, 3 Na-pyruvate, 0.5 CaCl_2_·4H2 O and 10 MgSO_4_·7H_2_O (pH ∼7.3 , ∼300 mOsm/L) (Ting et al., 2018) for the remainder. Samples were transported over 15-20 min to the laboratory in a sealed transport bottle. Slices of ∼350 µm thickness were prepared as described in Field et al. (2021), except for samples A-C the cutting ACSF was gradually replaced by recording ACSF, in which they were stored at room temperature until recording. The recording ACSF contained the following (in mM): 130 NaCl, 3.5 KCl, 1.3 NaH_2_PO_4_, 24 NaHCO_3_, 3 CaCl_2_, 1.5 MgSO_4_, 12.5 glucose, (pH ∼7.3, ∼300 mOsm/L). The storing ACSF contained the same components as the NMDG-based cutting ACSF except the NMDG was replaced with 92 mM NaCl. All solutions were continuously bubbled with carbogen.

### Electrophysiological recordings

Electrophysiological recordings were performed over 10-16 hours after slicing between 12 PM and 4 AM the next day as described previously (Bocchio et al., 2019; Field et al., 2021). Slices in the recording chamber were perfused continuously with recording ACSF saturated with carbogen at a flow rate of ∼10 ml/min using a peristaltic pump (Gilson). The time required for a new solution to reach the chamber was 45-60 seconds. Glass capillaries (4-7 MΩ) were filled with an internal solution containing the following (in mM): 126 K-gluconate, 4 KCl, 4 ATP-Mg, 0.3 GTP-Na_2_, 10 Na_2_-phosphocreatine, 10 HEPES, 0.03 ethylene glycol-bis(2-aminoethylether)-N,N,N’,N’-tetraacetic acid (EGTA), and 0.05% (w/v) biocytin (pH ∼7.3, 280–290 mOsmol/L). Neurons were visualised by differential interference contrast (DIC) microscopy with an Olympus BX51WI microscope equipped with a LUMPlanFL 60x water immersion objective (Olympus) and a digital camera (Zyla, ANDOR), connected to a desktop computer. Whole-cell patch-clamp recordings were performed from neurons in layers II and III, at 33-37 °C, using either an EPC-10 triple patch clamp amplifier with Patchmaster software (HEKA), or a Multiclamp 700B amplifier with pClamp software (Molecular Devices). Data was digitized at 100 kHz for current-clamp recordings and at 50 kHz for voltage-clamp recordings on the EPC-10 amplifier; on the Multiclamp amplifier data was digitized at 10 kHz in both modes. The reported voltage values are not compensated for a 16.5 mV junction potential. For each cell, voltage responses were recorded to a series of 800 ms long current square pulses starting from holding current -100 pA until rheobase (RB) +100 pA with 20 pA increments between sweeps in current clamp mode (I-V traces). The initial holding current was between 0 and -100 pA, and was aimed to be a multiple of 20 pA. Bridge balance was not adjusted during current clamp recordings. In paired recordings, a pair or a train of five action potentials (APs) were evoked at 50 ms intervals in one neuron by brief current injections, while the other neuron was continuously held at just subthreshold membrane potentials (-40 --45 mV, E_Cl_-≈ -94 mV) in voltage-clamp mode in order to detect GABA_A_ receptor mediated unitary evoked inhibitory postsynaptic currents (eIPSCs). For pair No-2 a test presynaptic AP was evoked 1 s after the last AP in the previous train (see fig. 3 D). Spontaneous inhibitory postsynaptic currents (sIPSCs) were recorded by continuously clamping the neurons to just subthreshold membrane potentials (-40 --45 mV). After recording a stable baseline for 3-5 min, group III mGluRs were activated by bath application of the orthosteric agonist, L-2-amino-4-phosphonobutyric acid (L-AP4, 50 or 300 μM). At the lower concentration, L-AP4 is thought to activate only the high affinity mGluR subtypes, 4 and 8, whereas at the higher concentration it would also activate mGluR7 (Cartmell & Schoepp, 2000; Kogo et al., 2004). Finally, the drug was allowed to wash out for 5-15 minutes, by perfusion of recording ACSF. Uncompensated series resistance was monitored every minute by application of a 10 ms voltage step of -10 mV. Action potentials and ionotropic glutamate receptors were not blocked.

In order to test the nature of the spontaneous synaptic currents recorded under our conditions we recorded 6 interneurons and 2 pyramidal cells in whole cell voltage clamp mode held at -50 mV membrane potential with an E_Cl_ = -94 mV. Four of the interneurons were visualised and they all confirmed to be interneurons with cell bodies in layer II. Two had sparsely spiny dendrites, whereas the dendrites of the other two were smooth and beaded. Two of the interneurons showed immunoreactivity to cannabinoid type 1 receptor (CB1) along their axonal membranes, one of them also being immunoreactive to vesicular glutamate transporter 3 (VGlut3) in its boutons and cholecystokinin (CCK) in the cell body. Further characteristics of these cells are not reported further in this study. Bath application of the GABA_A_ receptor antagonist gabazine (16 μM) completely abolished sIPSCs recorded in these interneurons and pyramidal cells, while sEPSCs remained unaffected (supplementary fig. 1 A). In an additional pair of visualised and synaptically connected interneurons located in layers III gabazine blocked the eIPSCs (supplementary fig. 1 B). The presynaptic interneuron was a bitufted neuron with long, radially running, smooth dendrites and dense local axon, and the postsynaptic interneuron was a multipolar smooth-dendritic cell, whose axon was not recovered. The presynaptic neuron was tested for CB1 in the axon and was immunonegative, and both cells were tested for VIP in their soma and both were immunonegative. The effect of gabazine is consistent with the prediction that spontaneous and evoked unitary IPSCs recorded under our experimental conditions were mediated by GABA_A_ receptors.

**Figure 1.**
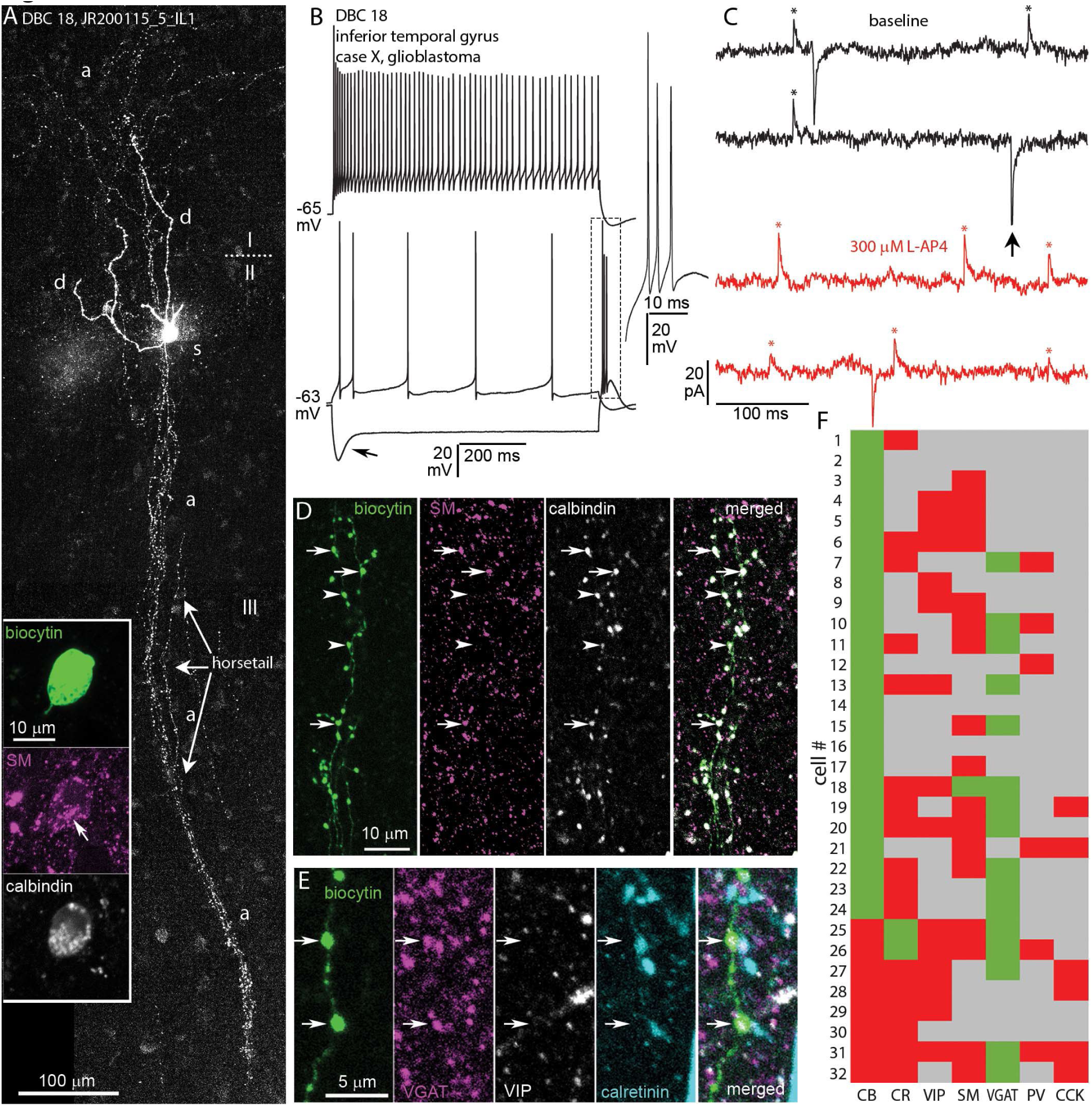
In vitro recorded and biocytin labelled DBC in the human inferotemporal neocortex. **A**. Tiled confocal fluorescent image of the cell with descending columnar “horsetail” axon. The cell body (s) is located in layer II with dendrites (d) penetrating layer I. The axon (a) diffusely innervates layer I (Z stack of 50 optical slices of 1.8 μm thickness, total depth, 45.9 μm); roman numbers indicate layers; inset: the cell body showing immunoreactivity for SM (Golgi apparatus, magenta, arrow) and CB (white) in the cytoplasm (Z stack of 5 optical slices of 0.9 μm thickness, total depth, 2.4 μm). **B**. Voltage responses of the cell in A to current injections of rheobase (RB), RB +100, and holding current -100 pA, showing a voltage sag (arrow) and rebound spikes (boxed area, inset) in response to hyperpolarising current injection and firing with spike frequency adaptation and strong spike amplitude accommodation when continuously depolarised; inset showing the expanded rebound burst from the boxed area. **C**. Example current traces recorded in voltage-clamp mode at -50 mV clamping potential, showing spontaneous inhibitory (asterisks, upward deflections, sIPSC) and excitatory (arrow, downward deflections, sEPSC) postsynaptic currents in baseline condition (black) and after bath application of 300 μM L-AP4 (red). **D**. Boutons of the cell in A are immunopositive for SM (magenta, arrows) and CB (white); note that some boutons are immunonegative for somatostatin (arrowheads) ( Z stack of 3 optical slices of 0.9 μm thickness, total depth, 1.8 μm). **E**. Boutons (arrows) of the cell in A are immunopositive for VGAT (magenta), and immunonegative for VIP (white) and CR (cyan) (Z stack of 3 optical slices of 0.9 μm thickness, total depth, 1.8 μm). **F**. Immunoreactivities in all recovered and tested neurons with descending axon bundles: green, immunopositive; red, immunonegative; grey, not tested/inconclusive.

**Figure 2.**
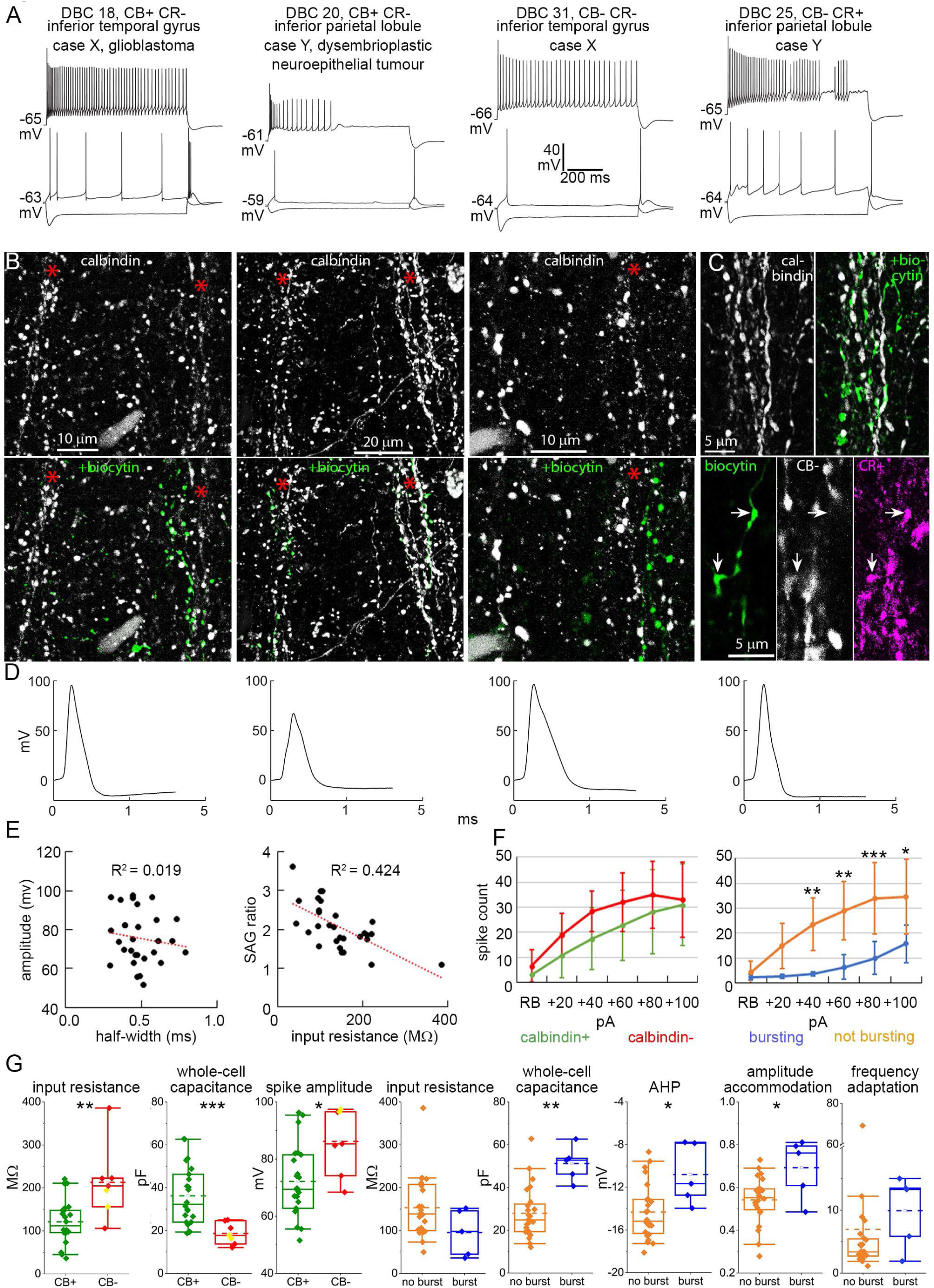
Diversity of DBCs with “horsetail” axons. **A.** Voltage responses of DBCs to current injections (RB, RB + 100 pA, and holding current – 100 pA steps). Immunoreactivity to CB and CR, cortical are and pathology are noted. **B.** Confocal fluorescent images of immunoreaction to CB (top, white) and overlayed visualisation of biocytin (bottom, green), revealing the contribution of individual DBC axons to CB+ axonal columns (asterisk; for DBC 18,20,31 respectively: Z stacks of 32,25,7 optical slices of 0.9,0.9,0.7 μm thickness, total depth, 13.3,11.2,3.0 μm). **C.** Top: Immunoreaction to CB (white) and overlayed axonal biocytin (green) for the CR+ DBC 25 above (Z stack of 14 optical slices of 0.9 μm thickness, total depth, 5.2 μm); Bottom: CR+ boutons of DBC 25 (arrows) are intermingled with but immunonegative for CB (Z stack of 3 optical slices of 0.9 μm thickness, total depth, 1.6 μm). **D.** Voltage traces of the first action potentials evoked at RB current injection for each cell in **A**. **E.** Scatter plots of action potential amplitude vs. half-width (left, Pearson corr. -0.136, p = 0.480); and sag ratio vs. input resistance (right, Pearson corr. -0.651, p = 1.3 * 10^-4^) of all DBCs (n = 29, adjusted R^2^ values are indicated). **F.** Firing index (number of spikes elicited by each 800 ms long current injection step) of CB+ (green) vs. CB-(red) DBCs (left, two-way repeated measures ANOVA, within subjects effect of current step: F = 50.712, p = 1.1 * 10^-7^; within subjects effect of interaction: F = 1.353, p = 0.246; between subjects effect of immunoreactivity: F = 2.164, p = 0.153) and of bursting (blue) vs. not bursting (orange) DBCs (right, two-way repeated measures ANOVA, within subjects effect of current step: F = 28.365, p = 1.2 * 10^-5^; within subjects effect of interaction: F = 7.201, p = 5.3*10^-6^; between subjects effect of bursting: F = 13.840, p = 9.2 * 10^-4^, Tukey pairwise comparisons: bursting vs not bursting within each current step: * p < 0.05, ** p < 0.01, *** p < 0.001; whiskers represent SD). **G.** Comparison of passive and active membrane properties between CB+, (green, n = 22) and CB-(red, n = 7) DBCs (CR+ cells, yellow), and between bursting (blue, n = 5) and not bursting (regular firing, orange, n = 24) DBCs. Boxes represent median and IQR, whiskers represent 5 and 95 percentiles, mean is indicated by dashed line. For all CB+ vs CB-, and the AHP and AP amplitude accommodation comparisons student’s t-test was used, for the remainder Mann-Whitney U-test, * p < 0.05, ** p < 0.01, *** p < 0.001

**Figure 3.**
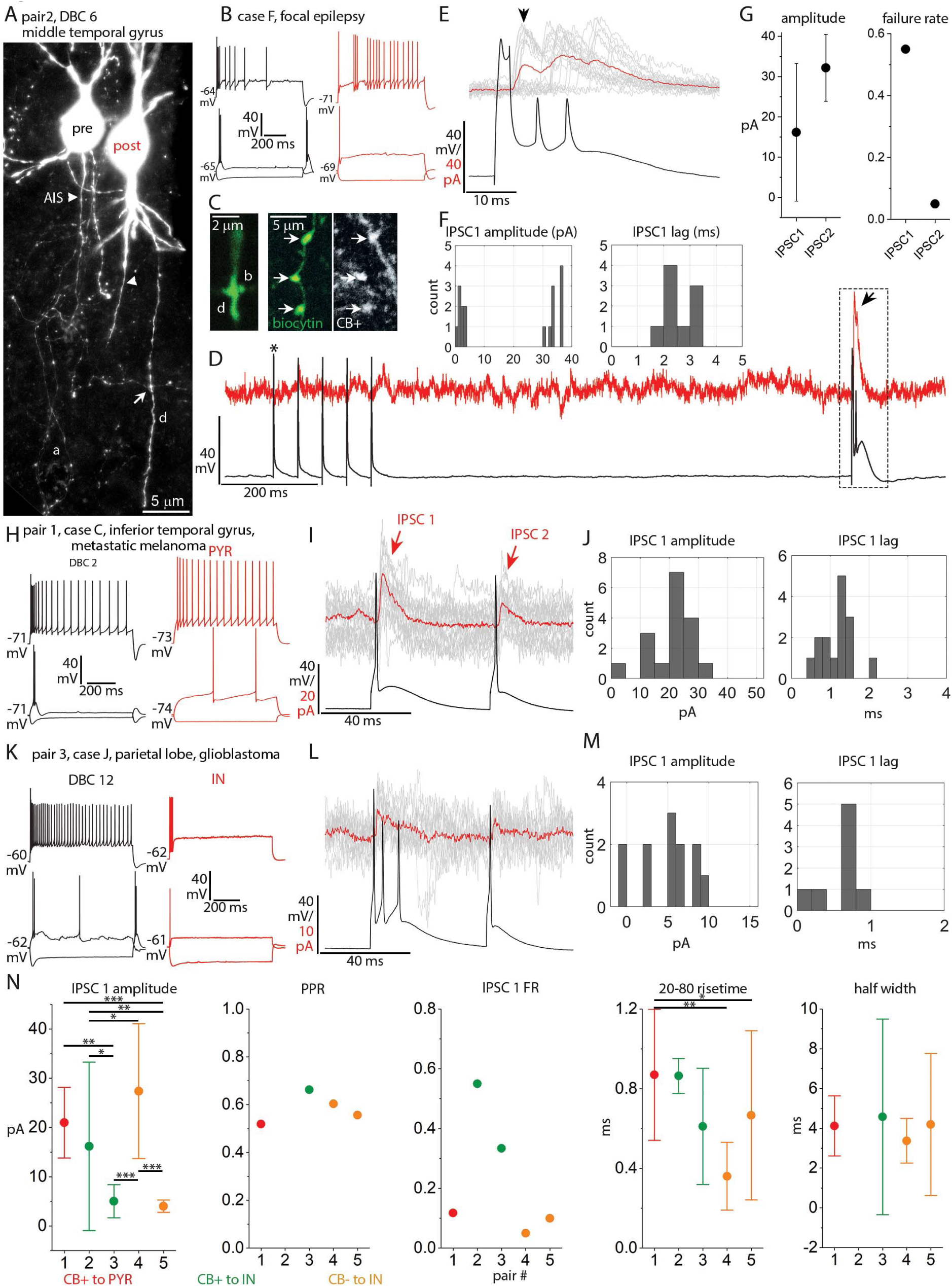
Synaptic currents evoked by DBCs. **A.** Fluorescent image of two simultaneously recorded synaptically connected interneurons (pair 2). The presynaptic DBC (pre) has a “horsetail” axon (a) and a collateral forms a bouton in close apposition to the dendrite (d, arrow) of the postsynaptic interneuron (post). Axon initial segments (AIS), arrowheads. (Z stack of 16 optical slices of 14.3 μm thickness, total depth, 42.8 μm). **B**. Voltage responses of the pre- (black) and postsynaptic (red) cells in **A** to current injections of RB, RB +100 and holding current-100 pA. **C. Left:** Fluorescent image of one contact site (arrow in A) showing the putative presynaptic bouton (b) in close juxtaposition with the postsynaptic dendrite (d, single optical slice of 0.7 μm thickness). **Right:** Boutons (arrows) of the presynaptic DBC (green) are CB+ (grey) (single optical slice of 0.9 μm thickness). **D.** Example of the current response in the postsynaptic cell (red, single sweep) voltage clamped at -47 mV to a train of single action potentials (APs, asterisk) 50 ms apart and an action potential burst (boxed area) evoked in the presynaptic DBC (black) in current clamp. There is an IPSC response to the burst (arrow) but not to the single APs (failures). **E.** IPSC responses in the postsynaptic cell (single responses in grey, arrow 1^st^ response, average of 20 responses in red) to the action potential burst in the presynaptic DBC (black, single sweep). **F.** Distributions of the peak amplitude (n = 20, including failures) and response lag (n = 9, excluding failures) of the IPSC after the first AP in the burst (bin width: 1 pA for amplitude and 0.5 ms for lag). **G.** Mean response amplitude including failures (left) and failure rate (right) of the IPSCs after the first and second APs in the burst (n = 20 for each of IPSC1 and 2, whiskers represent SD). **H.** Voltage responses of the presynaptic DBC 2 (black) and postsynaptic pyramidal cell (red) of Pair 1 to current injections of RB, RB +100 and holding current -100 pA. **I.** Current responses in the postsynaptic cell of pair 1 (single responses in grey, average of 17 responses in red) to a pair of APs evoked in the presynaptic DBC 2 (black) 50 ms apart. **J.** Distribution of peak amplitude (n = 17, including failures) and response lag (n = 15, excluding failures) of IPSC 1 (bin width: 5 pA for amplitude and 0.2 ms for lag). **K-M.** Same as **H-J** for pair 3 (bin width: 1 pA for amplitude and 0.2 ms for lag; n = 12 including, n = 8 excluding failures). The presynaptic DBC fires AP bursts to rheobase current injection and the postsynaptic cell is an interneuron (IN). **N.** Mean amplitude (including failures) of IPSC 1 (n = 17, 20, 12, 20 and 20 responses for neuron pairs 1-5 respectively; One-way ANOVA, F = 16.22, p = 4.7 * 10^-8^, Tukey post-hoc test; p < 0.05 for pairs 4 vs. 2 and 3 vs. 2; p < 0.01 for pairs 5 vs. 2 and 3 vs. 1; P < 0.001 for pairs 4 vs. 3, 4 vs. 5 and 5 vs. 1), paired-pulse ratio (PPR, average response2/response1 mean amplitude), failure rate, 20-80 % rise-time (One-way ANOVA, F = 7.37, p = 3 * 10^-4^, Tukey post-hoc test, p < 0.05 for pair 5 vs. 1; p < 0.001 for pair 4 vs. 1) and half-width (One-way ANOVA, F = 0.47, p = 0.707) of IPSC 1 for 5 DBC synapses onto different types of postsynaptic neuron. Whiskers represent SD, plots are colour coded according to immunoreactivity of the presynaptic DBC for CB and the postsynaptic neuron class (PYR: pyramidal cell; IN: interneuron).

### Data analysis and inclusion criteria

The following criteria were applied for inclusion of cells in the analysis: 1. The holding current was between 0 and -100 pA for holding the cell at -75 mV; 2. At least one overshooting AP could be elicited by depolarising current injections; 3. Fast pipette capacitance was successfully compensated and no oscillation artefacts were observed in current clamp mode. The following additional criteria were applied for inclusion of the voltage clamp recordings of eIPSCs and sIPSCs: 1. Series resistance (Rs), calculated as the ratio of the -10 mV voltage step and the peak capacitive current transient, was < 35 MΩ during the entire recording and did not change more than 25% from the initial value; 2. There were no sudden shifts in Rs.

PatchMaster (.dat) files were opened in Igor Pro software v7.0.8.1 (WaveMetrics) using Patchers’s Power Tools (Department of Membrane Biophysics, Max Planck Institute for Biophysical Chemistry, Göttingen, Germany^1^). After digitally adjusting the I-V traces to remove the artefact resulting from the bridge balance, the following parameters were measured: input resistance (IR) was calculated as the slope of the linear fit to subthreshold steady state membrane voltage values and the injected current (V = a + b*I, where V is the steady state membrane voltage, I is the injected current, and b is the input resistance). Resting membrane potential (Vm), was measured as the steady state voltage in response to 0 pA current injection, or was calculated as the constant (variable ‘a’) in the formula above. The membrane time constant (τ) and whole-cell capacitance (C_m_) were calculated from current-clamp recordings as described (Golowasch et al., 2009): τ was calculated as the time constant of the slowest component of a double-exponential fit to the voltage in response to holding current -100 pA current injection from the beginning of the step until the peak negative deflection 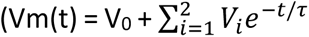 where V_0_ is the baseline membrane potential and V*_i_* and τ are the amplitude and the time constant of each exponential term); C_m_ was obtained by dividing τ with the amplitude of the voltage deflection associated to the slowest exponential term. The sag ratio was calculated as the ratio of the maximum and the steady state voltage deflection in response to holding current -100 pA current injection. The rheobase was measured as the current injected evoking the first AP. Single AP kinetic parameters were measured on the first AP elicited at RB current injection in Matlab R2020a (MathWorks), using a custom written script as described previously (Field et al., 2021).

Voltage-clamp recordings were filtered with a 2 kHz lowpass Bessel (8-pole) filter and boxcar smoothing (11 points) in Clampfit or IgorPro before analysis. Evoked IPSCs were detected with TaroTools toolbox for Igor Pro (https://sites.google.com/site/tarotoolsregister), using a threshold (4-7 pA) detection method, followed by visual inspection. Trials in which no eIPSC was detected, were considered as failures. The response lag was measured from the peak of the presynaptic AP to the onset of the eIPSC. The mean amplitude of eIPSCs were calculated from all trials, including failures. The paired-pulse ratio, an indicator of presynaptic release probability, was calculated as the mean amplitude of IPSC2 over IPSC1. The half-width and the risetime (20-80%) were calculated automatically by the detection software. Spontaneous IPSCs were analysed in Clampfit v10.7.0.3 (Molecular Devices) from all 3-5 minutes of baseline and the last two minutes of drug application to allow for the drug to reach maximum concentration in the recording chamber and achieve reasonable penetration into the slice. Washout periods were not analysed due to instability. For the detection of sIPSCs, first a typical IPSC was visually identified in the baseline period, and was used as a template for automatic detection. Events with an amplitude smaller than 5 pA, and a risetime (20-80%) longer than 5 ms were removed, and the remaining events were averaged to create a second template, which was then used in a second round of event detection. Control recordings for testing potential changes in sIPSCs over time were recorded alternately with test recordings during every session by perfusion of vehicle (recording ACSF) for 15-20 minutes.

All statistical tests were carried out in Origin Pro (v 94E 2017, OriginLab), and are reported in full in the results. Values are reported as mean ± standard deviation, unless stated otherwise. Normality of the data was tested with Kolmogorov-Smirnov test, and the appropriate parametric or non-parametric tests were applied. For data with < 10 observations, a non-parametric test was used. Differences were considered statistically different at alpha < 0.05.

### Streptavidin visualisation and immunohistochemistry

After completion of the electrophysiological recording, slices were immersed in fixative, containing 4% (w/v) paraformaldehyde (PFA) and 15% (v/v) saturated picric acid (PA) in 0.1 M PB at pH ∼7.2 at 4°C overnight. For samples from cases O-W (see table 2) the fixative also contained 0.05% (w/v) glutaraldehyde (GA). Next, slices were thoroughly washed and re-sectioned into 3-5, ∼60 µm thick sections on a vibratome as described previously (Field et al., 2021). From each slice two sections, including the one in which the soma was predicted to be found, were selected for fluorescent visualization. Sections were visualised using Alexa 488-conjugated streptavidin (1:1000) as described previously (Bocchio et al., 2019; Field et al., 2021).

The expression of multiple molecules in labelled cells and in control tissue were tested using immunohistochemical reactions on single 60-70 µm thick sections. Immunohistochemical reactions were performed as described previously (Unal et al., 2015; Joshi et al., 2017). Up to five different primary antibodies (ab), produced in different host species, against the epitopes of the molecules of interest were used in appropriate dilutions on any single section. For a list of all primary abs see table 1. Secondary abs against immunoglobulin G of the host species of the primary ab, conjugated to different fluorophores, were added at appropriate dilutions (Alexa405/DyLight405 (blue)- and cyanine5 (Cy5)/DyLight649 (infra-red)-conjugated abs at 1:250; Alexa488 (green)-conjugated abs at 1:1000; Cy3 (red)-conjugated abs at 1:400). All secondary abs were raised in donkey (Jackson Immuno Research).

### Wide field epifluorescence and confocal laser scanning microscopy

Cells visualised with Alexa 488-conjugated streptavidin and fluorescent immunoreactions were first evaluated in a wide field epifluorescence microscope (Leitz DMRB, Leica) using PL FLuotar 20x/0.5 and 40x/0.7 lenses and photographed with an ORCA ER digital camera (Hamamatsu Photonics) controlled by OpenLab software (Improvision). The light source was either a pE-300 LED lamp (CoolLED) or a mercury arc lamp (HBO, Osram), for which the light was spectrally separated by dichroic mirrors to obtain optimal excitation and emission bandwidths for each fluorophore. For higher resolution imaging, confocal laser scanning microscopy was performed using an LSM 710 axioImager.Z1 microscope (Zeiss) and DIC M27 Plan-Apochromat 40x/1.3, 63x/1.4 and alpha Plan-Apochromat 100x/1.46 oil immersion objective lenses, controlled by ZEN 2008 software (v 5, Zeiss), as described previously in detail (Lasztóczi et al., 2011). Signal from each fluorophore was recorded in separate scanning tracks and channels, using the following lasers: for Alexa405 and DyLight405, a 405 nm solid state laser; for Alexa488, a 488 nm argon laser; for Cy3, a 543 nm He-Ne laser; for Cy5 and DyLight647, a 633 He-Ne laser. Pinhole size was adjusted optimally for similar optical slice thickness between tracks. The step size along the Z imaging axis was set to half of the thickness of the optical slices, as per the Nyquist criterion for optimal sampling. Details of the optical slice thickness are given in the figure descriptions. Some neurons were imaged in the entire ∼60 μm thick section to demonstrate the distribution of their dendrites and axon (see fig. 1A). In order to remove the autofluorescence signal of the lipofuscin, these sections were imaged in a spectral scanning mode (λ-stack mode in ZEN software) with a fully opened pinhole, after which regions of interest including the labelled processes of neurons, lipofuscin and the background respectively were selected to determine the emission spectra of each. This was used to separate the signals into different pseudo channels by spectral linear unmixing (Dickinson et al., 2001), which was performed in the ZEN software.

### Peroxidase reactions

Sections of labelled cells used for immunohistochemistry were ultimately converted by avidin-biotin horseradish peroxidase (HRP) reaction with 3,3’-diaminobenzidine (DAB) as chromogen for the visualisation of biocytin and embedded in epoxy resin (Durcupan, Sigma) for conventional light microscopic examination. Sections were incubated in biotinylated peroxidase complex (B) 1:100 v/v (Vectastatin ABC elite kit, Vector Laboratories) in TBS-Tx for 4 hrs at room temperature, then incubated in avidin + B (A+B)1:100 v/v in TBS-Tx over 36 hrs at 4°C. Next, sections were pre-incubated in 0.5 mg/ml DAB in Tris Buffer (TB) for 10 minutes in dark. Subsequently, 0.002% w/v H_2_O_2_ substrate was added to initiate precipitation of DAB. After 12 to 20 minutes of conversion, depending on the intensity of the labelling, reactions were stopped and sections were washed 4x10 minutes in 0.1M PB. For contrast enhancement, sections were incubated in 0.5% w/v Osmium tetroxide solution in 0.1M PB for 1 hour at room temperature. Before mounting, sections were dehydrated using increasing concentrations of ethanol and final steps of propylene oxide (Sigma) for 2x10 min. From propylene oxide, sections were quickly transferred into Durcupan epoxy resin (Sigma) and left overnight and finally mounted on glass slides.

For electron microscopy, sections of slices fixed without GA were postfixed with same fixative as above with the addition of 0.05% (w/v) GA for 2 hrs at 4°C. Sections were then incubated in 20% sucrose (Sigma Aldrich) in 0.1M PB for 2 hrs at room temperature for cryoprotection, followed by quickly freezing the sections using liquid nitrogen and thawing them in PB with sucrose (1-3 times). All following steps were the same as for HRP reactions on Triton-treated sections with the exception that the buffer did not contain detergent. During dehydration, these sections were treated with 1% (w/v) uranyl acetate (UA) dissolved in 70% ethanol for 40 min in dark for further enhancement of contrast for electron-microscopy.

### Light microscopy and reconstruction of labelled cells

Transmitted light microscopy of HRP-converted sections was performed using a Leitz Dialux22 microscope (Leica), equipped with NPL Fluotar 10x/0.3, 25x/0.55, 40x/0.78 dry and PL Apo 100x/1.32 oil immersion objective lenses. Images were taken using a Canon EOS 40D digital camera controlled by EOS Canon Utility software. Some neurons were reconstructed from multiple sections in order to demonstrate the distribution of their dendrites and axon across different neocortical layers, as well as for the identification of their axonal boutons, which were later examined using electron-microscopy. For 2D reconstruction of labelled cells (see figs 4A and 7A), neurons were manually traced using a drawing tube attached to a transmitted light microscope (Leitz Dialux22, Leica), equipped with a Pl Apo 63x/1.4 oil immersion objective. The India ink drawings were digitised using a digital camera (EOS 40D, Canon) and processed in Adobe Photoshop. Digital 3D reconstruction of some neurons (see fig. 6A) was performed with Neurolucida software (v 2020.2.2 MBF Bioscience) and an Axioskop 2 mot plus (Zeiss) equipped with a Plan Apochromat 100x/1.4 Oil lens and Retiga 2000R (QImaging) digital camera, or with a v 2021.1.3 software and a Leica DMR microscope equipped with a HCX PL Fluotar 100x/1.30 Oil lens and Lumina HR (mbf bioscience) digital camera.

### Serial section transmission electron microscopy of labelled cells and reconstruction of target dendrites

After the labelled neurons were reconstructed, some were selected for electron-microscopic study of their synapses (see table 3 and fig. 6). Areas of the HRP-converted sections containing the “horsetail” axon of DBCs in layer III were removed from the glass slides and re-embedded in cylindrical blocks of epoxy resin. Electron-microscopic sections (∼50 nm) were cut using an Ultra 45 Diatome diamond knife (TAAB), and mounted on pioloform-coated copper slot grids (2x1 mm, 10-20 sections/grid). Grids were examined using either a Fei Tecnai 12 transmission electron microscope (TEM) at 120 keV accelerating voltage and photographed with a Gatan OneView CMOS camera controlled by Gatan Digital Micrograph software (GMS3), or using a Jeol-1011 microscope at 80 keV accelerating voltage and photographed with a TRS Sharp Eye camera controlled by iTEM software. The labelled processes and their synaptic targets were photographed in serial electron microscopic sections, were imported into TrackEM2 reconstruction plugin of ImageJ software and were aligned using an automatic non-linear alignment method, based on block correspondence. After alignment the axon and boutons of the labelled cell, the target structure and the synaptic specialisation was traced through the serial sections and was rendered into a 3D reconstruction (see fig. 6 K, L).

**Table 3.**
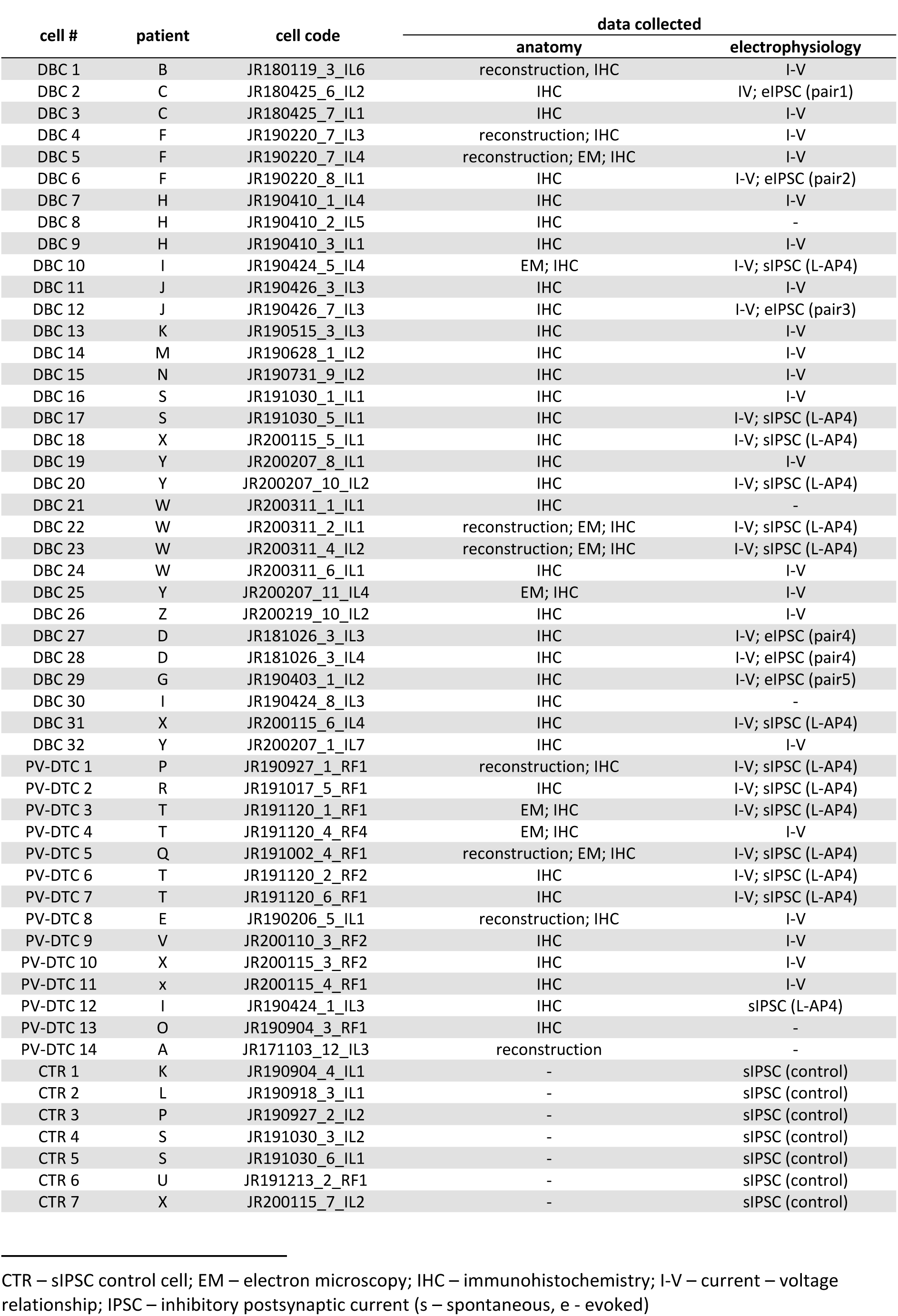
Information on recorded and labelled DBCs, PV-DTCs and cells used for pharmacological control measurements

### Post-embedding immunohistochemistry for GABA and counting of synapses

Samples AA-CC were fixed with 4% (w/v) PFA, 2.5% (w/v) GA and 15% (v/v) PA and were used for post-embedding immunohistochemistry for GABA. Sections of these samples were treated with 1% (w/v) Osmium tetroxide, treated with 1% (w/v) UA and mounted on glass slides in epoxy resin (Durcupan). From each sample one area of layer III was re-embedded for electron microscopy. Serial sections of ∼60 nm thickness were mounted on pioloform coated nickel slot grids (2x1 mm). Post-embedding immunohistochemistry was performed as described (Somogyi & Soltész, 1986) using rabbit antiserum to GABA at 1:1000 dilution (Hodgson et al., 1985) and 6 nm gold-coupled anti rabbit IgG (Aurion) as secondary antibody. Silver enhancement was used for easier detection of the particles in electron micrographs as recommended by the manufacturer (Aurion).

For counting synapses, electron micrographs of serial immunoreacted sections (5-10) were taken with a Philips CM100 transmission electron microscope equipped with a Gatan UltraScan 1000 CCD camera at 80 keV accelerating voltage. Four, five and two different areas were photographed from samples AA, BB and CC respectively, chosen not to contain somata or blood vessels. Electron micrographs were imported in trackEM2 software and serial sections were aligned as described above. Synapses were counted using a stereological method, by tracing the pre-and post-synaptic elements in 3D through a specified volume within a 3D counting frame (West, 1999; Cano-Astorga et al., 2021). The first section and the top and left sides formed the exclusion planes for any synapse, whereas the last section and the bottom and right sides formed the inclusion planes. Synapses were excluded if the synaptic specialisation in the pre- or post-synaptic membrane (see fig. 8) touched any of the exclusion planes. The volume of the counting frames were corrected by a factor of 1.23 compared to the scale of the micrographs to compensate for compression along the direction of cutting of the electron microscopic sections. This was calculated by measuring in the direction of cutting the block surface width and the section width after cutting. Thus the calculated density of synapses is given for fixed, osmicated and dehydrated specimens, not the living brain.

## Results

### Recorded cells and samples

We recorded a total of 622 neurons *in vitro* in samples from 56 (4 epilepsy, 52 tumour) patients, of which 569 were visualised and fully or partially recovered for anatomical analysis. Of these neurons, 356 (∼63 %) were smooth dendritic or sparsely spiny interneurons and 213 (∼37 %) were spiny pyramidal cells, with cell bodies in either layer II or III due to our selection. We further grouped interneurons based on the distribution of their axons and boutons, and selected two easily distinguishable groups, which had adequate number of cells for comparison. Interneurons with cell body in layer II or upper layer III and a radially oriented axonal plexus, restricted to a narrow (10 – 20 μm) cortical column descending through layers II to V were identified as DBCs (n = 32, fig. 1) (Somogyi & Cowey, 1981; DeFelipe et al., 2002; DeFelipe, 2011). Another group of interneurons had widely distributed axon projecting more than 200 μm in every direction from the soma and large, multipolar dendritic trees. Subsequent analysis showed that these cells express the calcium binding protein parvalbumin (PV), thus we refer to them as *parvalbumin-expressing dendrite-targeting cells* (PV-DTC, n = 14, fig. 4), based on their synaptic targets (see below). We studied the dendritic and axonal distributions, firing patterns, molecular expression and synaptic connections of these two groups of cells in detail to better understand their correspondence to well defined cell types in other species. Furthermore, we compared the effects of group III mGluR activation on spontaneous GABAergic synaptic inputs in these two cell types. A total of 53 neurons were analysed in samples from 29 patients, including 14 different cortical regions and 13 different diagnoses (table 2). An additional three samples were imported from the University of Szeged, Hungary, for post-embedding immunohistochemistry and electron microscopic study of GABAergic synapses (table 2).

**Figure 4.**
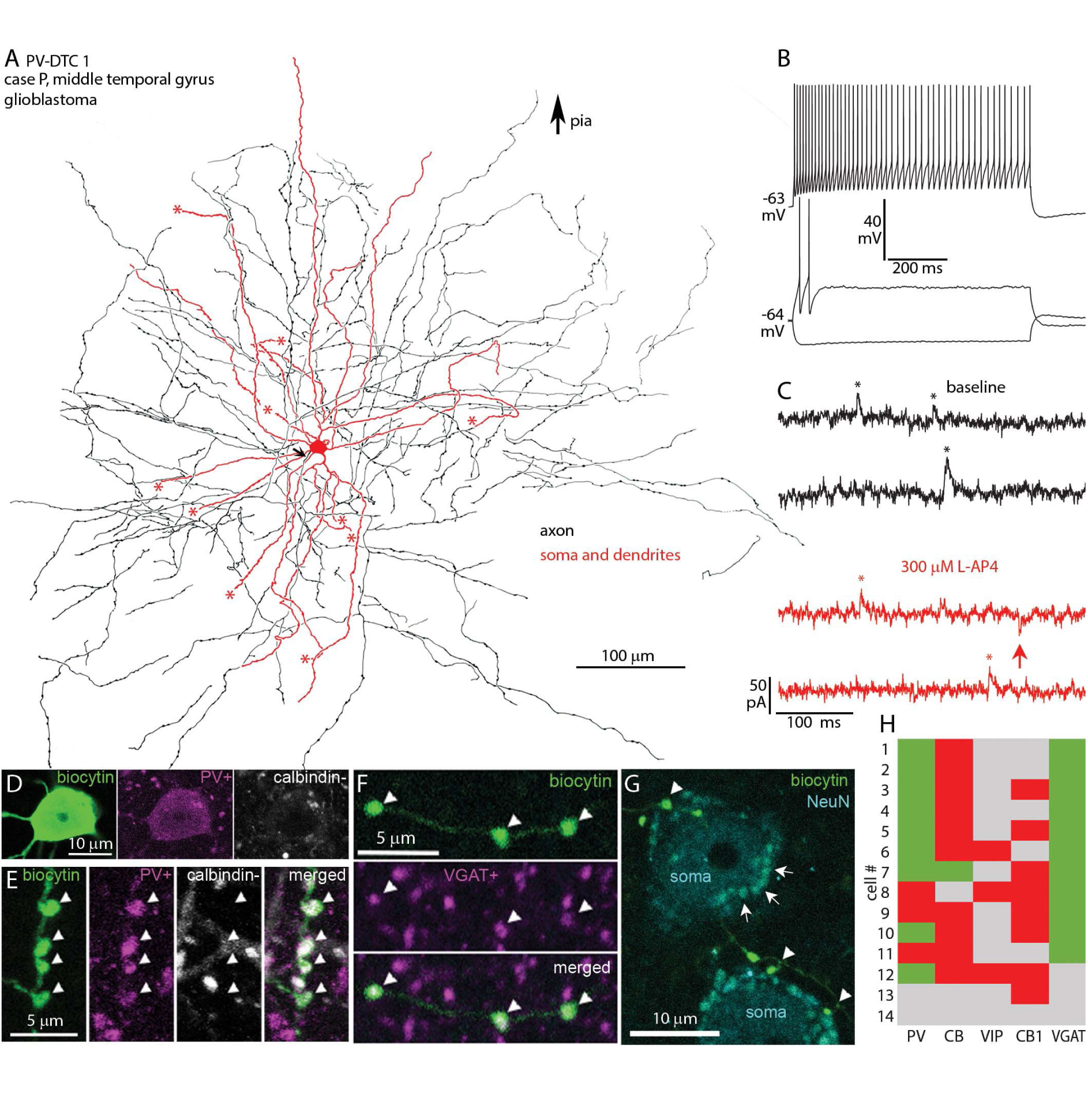
In vitro recorded and biocytin labelled PV-DTC in the human temporal neocortex. **A**. Partial 2D reconstruction (2 sections of ∼ 60 μm thickness) of the cell with cell body in layer III; the axon did not enter layer I. The axon initial segment (arrow), cut ends of dendrites (asterisks) and direction of the pial surface are indicated. **B**. Voltage responses of the cell in A to current injections of RB, RB +150, and holding current -150 pA. Firing is a regular train of action potentials with spike frequency adaptation and small spike amplitude accommodation. There is a minimal voltage sag and no rebound spikes in response to hyperpolarising current injection. **C**. Example current traces recorded in voltage-clamp mode at -50 mV clamping potential, showing sIPSCs (asterisks, upward deflections) and sEPSCs (arrow, downward deflections) in baseline condition (black) and after bath application of 300 μM L-AP4 (red). **D**. The cell body is immunopositive for PV (magenta) and immunonegative for CB (white) in the cytoplasm (Z stack of 9 optical slices of 0.9 μm thickness, total depth, 4.7 μm). **E.** Boutons of the cell in A are immunopositive for PV (magenta, arrowheads) and immunonegative for CB (white) (Z stack of 13 optical slices of 0.9 μm thickness, total depth, 7.9 μm). **F**. Boutons (arrowheads) of the cell in A are immunopositive for VGAT (magenta) (single optical slice of 0.7 μm thickness). **G.** Boutons (arrowheads) of the cell in **A** (biocytin, green) are in close proximity with NeuN (cyan) immunopositive cell bodies (soma), which also contain autofluorescent lipofuscin granules (e.g. arrows). (Z stack of 4 optical slices of 0.7 μm thickness, total depth, 2.4 μm). **H**. Immunoreactivities in all recovered and tested neurons with multipolar dendrites: green, immunopositive; red, immunonegative; grey, not tested/inconclusive.

### Axonal and dendritic features and immunohistochemical characterisation of DBCs

The distinguishing feature of DBCs is their “horsetail”-like axon (Jones, 1975; Somogyi & Cowey, 1981; Meyer, 1983; Lund & Wu, 1997; DeFelipe, 2011), formed of at least three, but often more, interwoven axon branches traveling in the radial direction from the soma in layer II and upper layer III, towards layers V and VI, forming a 10-20 μm wide columnar plexus (fig. 1 A; fig. 6 A). We visualised 32 DBCs for anatomical and immunohistochemical analysis. Of these, the current-voltage response of 29 DBCs were analysed (table 3), three being excluded because of bad capacitance compensation resulting from changing ACSF levels in the recording chamber.

The axon of DBCs could be traced as deep as layer V (see fig. 6 A). The number of branches in the “horsetails” varied between 3 and 7. Occasionally, short collaterals branched from the “horsetail” at different depths and travelled laterally but remained within 100 μm from the “horsetail” (see fig. 6 A). The majority of DBC boutons were *en-passant*, but all DBCs had a few *en-termineux* boutons on axonal side branches. In addition to the “horsetail” DBCs also had axon around the cell body and in layer I, where it did not form a columnar structure, but spread 100 - 500 μm laterally (fig. 1 A). The axon of 22 DBCs entered layer I, with one (DBC 19) exclusively innervating a narrow subpial band. The axon of 9 DBCs spread uniformly across the depth of layer I, whereas another 9 innervated only the deep half. Three DBCs had only a few collaterals entering layer I. The dendrites of 29 DBCs were recovered, of which 24 were smooth, whereas the remainder were sparsely spiny. The dendritic trees were usually small with a median of 4 (range: 2 - 7) main branches originating from the soma, which travelled mostly towards the pial surface, and less than 100 μm laterally. The distribution of the dendrites was in most cases unipolar towards the pia (n = 18), or bipolar in the radial direction (n = 9). Two DBCs had uniformly distributed dendritic trees around their soma. All, except DBC 1 had dendrites in layer I.

Previous studies have demonstrated calbindin-immunopositive (CB+) radial axonal columns in human (del Río & DeFelipe, 1995, 1997b; Ballesteros Yáñez et al., 2005; DeFelipe et al., 2006) and monkey cortex (DeFelipe et al., 1989; DeFelipe et al., 1990; Peters & Sethares, 1997), which were predicted to originate from DBCs. We have confirmed the presence of such CB+ axon bundles reminiscent of the “horsetail” axon of DBCs, in the temporal, frontal and parietal areas of the human neocortex. By labelling individual DBCs with biocytin we were able to test directly their immunoreactivity for CB and establish the relationship of DBC “horsetails” to axonal bundles revealed by immunoreaction against CB. Furthermore, because interneurons with descending axons were found to be immunoreactive to the calcium binding protein calretinin (CR) and the neuropeptide VIP in rat (Kawaguchi & Kubota, 1996), monkey (Lund & Lewis, 1993; Condé et al., 1994; Meskenaite, 1997; Zaitsev et al., 2005) and human (Varga et al., 2015), we also tested if human DBCs were immunoreactive for these molecules. Other neuropeptides, such as somatostatin (SM) and cholecystokinin (CCK) were also tested. Finally, the calcium binding protein PV was also tested. All DBCs were tested for CB and 24 (75 %) were immunopositive in all three compartments (axon, soma and dendrites, n = 11), only in their axons (n = 8), only in their somata (n = 1), or in their somata and axons (n = 4). Eleven of these, tested for CR on their axon (n = 8), soma (n = 1) or both (n = 2) were immunonegative. Two DBCs were immunopositive for CR in their axons, and these were immunonegative for CB. Six DBCs were immunonegative for both CR and CB. None of the 14 tested DBCs (8 CB+, 2 CR+, 4 double immunonegative) were immunopositive for VIP in their axons (n = 11), somata (n = 1), or both (n = 2). Similarly, none of the 6 DBCs tested were immunopositive for PV in their axons or CCK in their somata (fig. 1 F). Of the 18 cells tested, only one cell (DBC 18) was immunopositive for SM and this cell was also immunopositive for CB (fig. 1 A, D, F). Interestingly, although there was granular immunoreactivity for SM in the soma of DBC 18, characteristic of peptides found in the Golgi apparatus (fig. 1 A), not all of its boutons were SM-immunoreactive (fig. 1 D). All 16 cells tested were immunopositive for the vesicular GABA transporter in their boutons (VGAT, fig. 1 E, F).

We found CB+ DBCs in the temporal (n = 13), frontal (n = 4), parietal (n = 4) and parieto-occippital (n = 3) cortices. Calbindin-immunonegative DBCs were found in the temporal (n = 4) and parietal (n = 4) cortices. One CR+ DBC was found in the temporal, whereas the other in the parietal cortex. The only SM immunopositive DBC was in the temporal cortex. The “horsetail” axons of the recorded and biocytin-labelled CB+ DBCs always aligned with the CB+ axonal bundles, where they intermingled with CB+ axons from unknown neurons (fig. 2 B 1^st^ and 2^nd^ columns). Furthermore, the axons of CR+ and CB-/CR-double immunonegative DBCs also aligned with and travelled within CB+ bundles (fig. 2 B, 3^rd^ column and fig. 2 C). While most DBCs’ axon formed one “horsetail”, DBCs 1 and 4 formed two, and DBC 20 formed three separate “horsetails”, each aligned with a different CB+ bundle (fig. 2 B, 2^nd^ column). However, cells with one main “horsetail” would also give off collateral branches leaving the main CB+ bundle and entering a neighbouring one, forming boutons also between bundles. Finally, in one case, where two DBCs (DBCs 27 and 28) were labelled close to each other, their axons intermingled to form a single “horsetail”. These results show that the cortical columns determined by the CB+ bundles are complex structures including the “horsetail” axons of multiple CB+ as well as CR+ DBCs, among other cells.

### Intrinsic membrane properties of DBCs

We have found that DBCs express firing phenotypes that distinguish them from PV-DTCs and other cell interneuron types in the neocortex. All DBCs, except DBCs 28 and 29, responded with a prominent voltage sag (fig. 1 B; 2 A; 3 B, H, K; 5 A; sag ratio: 2.07 ± 0.58, n = 29) and 19 of 32 DBCs also fired rebound spikes after continuous hyperpolarisation (fig. 2 A; 3 B, K; 5 A). Five of 32 DBCs showed burst firing either at rheobase current injection and/or as rebound after continuous hyperpolarisation (fig. 1 B), and all five were CB+. The recorded CB-DBCs did not show burst firing. When continuously depolarised by rheobase + 100 pA current injection, 14 DBCs fired a regular train of action potentials (Fig. 2 A, DBCs 18 and 31), 5 showed depolarisation block (fig. 2 A, DBC 20), and 7 fired either irregular or stuttering action potential trains (fig. 2 A, DBC 25). All DBCs showed strong spike amplitude accommodation (average 0.57 ± 0.12), and spike frequency adaptation (median: 3.43, IQR: 3.52). At rheobase + 100 pA current injection, DBCs fired at an average firing rate of 38.1 ± 19.7 Hz, but their coefficient of variation of the inter-spike-intervals (ISI-CV), a measure of the regularity of the spike train, varied between 0.09 for the most regularly firing cell (DBC 24) to 2.06 for DBC 11, with a stuttering firing pattern (average: 0.47 ± 0.39).

The action potential (AP) kinetics of DBCs were variable (fig. 2 D). Overshooting AP amplitude varied between 52 and 97 mV, the average being 75 ± 2.5 mV (n = 29). The resting membrane potential was -56.8 ± 4.3 mV. The AP half-width varied between 0.29 and 0.79 ms, the average being 0.48 ± 0.02 ms, as measured on the first elicited AP at rheobase current injection. There was no correlation between AP amplitude and half-width (fig. 2 E, left, Pearson corr. -0.136, p = 0.480). Input resistance and sag ratio also varied across DBCs and were inversely correlated (fig. 2 E, right, Pearson corr. -0.651, p = 1.3 * 10^-4^).

Next, we compared the firing patterns, the passive and active membrane properties of CB+ (n = 22) vs CB-DBCs (n = 7), and bursting (n = 5) vs regular firing (not bursting) DBCs (n = 24). The firing index plots (number of action potentials per current step) indicated no difference in input gain between CB+ and CB-DBCs (fig. 2 F, left, two-way repeated measures ANOVA, within subjects effect of current step: F = 50.712, p = 1.1 * 10^-7^; within subjects effect of interaction: F = 1.353, p = 0.246; between subjects effect of immunoreactivity: F = 2.164, p = 0.153). Furthermore, those CB+ DBCs which showed a bursting phenotype fired significantly fewer action potentials than regular firing DBCs, which include both CB+ and CB-DBCs, at every current injection step above rheobase +20 pA (fig. 2 F, right, two-way repeated measures ANOVA, within subjects effect of current step: F = 28.365, p = 1.2 * 10^-5^; within subjects effect of interaction: F = 7.201, p = 5.3*10^-6^; between subjects effect of bursting: F = 13.840, p = 9.2 * 10^-4^, Tukey pairwise comparisons: bursting vs not bursting within each current step: * p < 0.05, ** p < 0.01, *** p < 0.001). Indeed, bursting DBCs often ceased firing after the initial burst even at large depolarising current injections. The input resistance of CB+ DBCs was lower than that of CB-DBCs (fig. 2 G, 121.1 ± 50.7 MΩ for CB+ DBCs vs 212.9 ± 86.8 MΩ for CB-DBCs, student’s t-test t(27) = -3.49, p = 0.002). Conversely, whole-cell capacitance was higher in CB+ DBCs than in CB-ones (36.2 ± 13.8 pF for CB+ DBCs vs 18.6 ± 5.1 pF for CB-DBCs, Welch’s t-test t(26.07) = 4.98, p = 3.5 * 10^-5^), indicating that CB+ DBCs are larger cells with bigger surface area. Finally, action potential amplitude was larger in CB-DBCs than in CB+ ones (72.2 ± 12.5 mV for CB+ DBCs vs 86.2 ± 11.6 mV for CB-DBCs, student’s t-test t(27) = -2.61, p = 0.014). The sag ratio, AP threshold, AP half-width, AHP, and the ISI CV were not different between CB+ and CB-DBCs.

Bursting DBCs had similar input resistance to regular firing (not bursting) DBCs (fig. 2 G, median: 138.5, IQR: 110.2 MΩ for not bursting vs median: 97.8, IQR: 108.6 MΩ for bursting, Mann-Whitney U test Z(88) = 1.59, p = 0.112), whereas the whole-cell capacitance was higher in bursting DBCs than in regular firing ones (median: 24.8, IQR: 13.4 pF for not bursting vs median: 52.8, IQR: 14.7 pF for bursting, Mann-Whitney U test Z(8) = -2.97, p = 0.003). The amplitude of the AHP was larger in non-bursting DBCs than in bursting ones (-14.3 ± 2.6 mV for not bursting vs -10.8 ± 2.8 mV for bursting, student’s t-test t(27) = -2.71, p = 0.012). Finally, not bursting DBCs showed stronger AP amplitude accommodation than bursting DBCs (0.54 ± 0.11 for not bursting vs 0.69 ± 0.14 for bursting, student’s t-test t(27) = -2.72, p = 0.011), whereas AP frequency adaptation was not different (fig. 2 G; median: 3.36, IQR: 2.78 for not bursting vs median: 13.3, IQR: 10.39 for bursting, Mann-Whitney U test Z(30) = -1.70, p = 0.089).

### Spontaneous and evoked IPSCs are mediated by GABAA receptors

We tested if GABAergic synaptic transmission to different types of interneuron in the human neocortex is differentially regulated by group III mGluRs. Specifically, we tested whether activation of group III mGluRs by the orthosteric agonist *L-2-amino-4-phosphonobutyric acid* (L-AP4), changes the frequency and/or amplitude of GABA_A_ receptor mediated spontaneous inhibitory postsynaptic currents (sIPSCs), as has been shown in rodents (Mitchell & Silver, 2000; Giustizieri et al., 2005; Cuomo et al., 2009). We refer to outward currents as sIPSCs, because under our experimental conditions in 4 visualised interneurons and 2 pyramidal cells the GABA_A_ receptor antagonist gabazine (16 μM) completely abolished outward currents, without changing inward currents (Supplementary fig. 1 and see Methods). It is important to note, that action potentials and fast glutamatergic synaptic transmission were not blocked in these experiments.

### Unitary synaptic currents evoked by DBCs

The target cell types and subcellular target domains of DBC efferent synapses and their postsynaptic effects are not known in humans. Following previous work in other species (Tamás et al., 1997a; Jiang et al., 2015), we tested unitary synaptic interactions between pairs of neurons consisting of one DBC and one other GABAergic interneuron or a pyramidal cell within a 50 µm distance from the DBC soma. We tentatively identified DBCs during the recordings based on their characteristic firing. We succeeded in testing 12 unitary synaptic connections from DBCs to interneurons and 4 connections to pyramidal cells, and detected postsynaptic responses in 3 interneurons (DBC-to-IN) and one pyramidal cell (DBC-to-PYR), a 25% connectivity ratio (fig. 3). We also found two unitary synaptic connections from interneurons to DBCs out of 9 tested.

The identity of presynaptic DBCs (n=5) was confirmed by light microscopy based on their “horsetail” axon (fig. 3A). Three of them (DBCs 2, 6 and 12) were immunopositive for CB (fig, 3C), the other two (DBCs 28 and 29) were CB-/CR-double immunonegative (see fig. 1F). The cell bodies of the recorded cell pairs were located in layers II or III. One of the postsynaptic interneurons (pair 4), was also identified as a CB-/CR-double immunonegative DBC (DBC 27, see fig. 1F); there was no detectable connection from DBC 27 to DBC 28. We were not able to determine the cell type of the other three postsynaptic interneurons. One of them (pair 2), had a bipolar dendritic tree originating from two main trunks at the superficial and deep poles of the soma (fig. 3A), and its axon had loose branches descending towards deeper layers. Its firing pattern was stuttering and showed spike amplitude accommodation; it had no sag potential (fig. 3B). In rodents, similar neurons are thought to express VIP. However, this postsynaptic human interneuron was double immunonegative for VIP as well as for CB. In pair 3, the postsynaptic cell had a firing pattern resembling that of multipolar basket cells, such as high frequency regular firing and depolarisation block (fig. 3K), had a small sag potential and a multipolar dendritic tree. In contrast, the postsynaptic interneuron in pair 5 had a slow irregular firing pattern, no sag potential and dendrites oriented radially. This cell was immunonegative for CR, VIP and CB. The axons of two postsynaptic interneurons (pairs 3 and were not recovered. In pair 1, the postsynaptic pyramidal cell was identified based on its densely spiny dendrites.

In order to test for unitary synaptic connections, we evoked pairs or trains (max. 5) of action potentials 50 ms apart in one cell held in current clamp between -70 and -75 mV by brief (1 – 5 ms) current injections of 100 – 500 pA, while recording the other neuron in voltage-clamp mode at just below firing threshold potentials (-45 to -50 mV). The peak amplitude of the evoked Inhibitory postsynaptic currents (eIPSCs), the paired-pulse ratio (PPR, amplitude of IPSC2 / amplitude IPSC1) and the failure rate of the first eIPSC were evaluated, as well as the 20 to 80 % risetime and the half-width of the first eIPSC. In pair 2, single APs evoked 50 ms apart in the presynaptic DBC did not elicit IPSCs in the postsynaptic cell; only evoking a presynaptic AP burst resulted in detectable bursts of postsynaptic IPSCs (fig. 3D, E). In this case the PPR was not available. In all other pairs, single APs elicited IPSCs in the postsynaptic cells. However, in pair 3, although single APs could elicit detectable IPSCs, the first presynaptic stimulus was always a burst of 2-3 APs, even at the lowest current injection (fig. 3L).

We found large variability in the response parameters of the 5 synaptic connections made by DBCs onto interneurons and pyramidal cells. The mean amplitude of the first IPSC in the DBC-to-PYR connection (pair 1) was 21 ± 1.7 pA (n = 17, including failures), with a failure rate of 0.12 and a PPR of 0.52. In the DBC-to-IN connections, the amplitude was 16.2 ± 3.8 pA (n = 20, including failures) in pair 2, it was significantly higher in pair 4 (27.4 ± 3.1 pA, n = 20, including failures) and was significantly lower in pairs 3 (5 ± 0.9 pA, n = 12, including failures) and 5 (4 ± 0.3 pA, n = 20, including failures; One-way ANOVA, F = 16.22, p = 4.7 * 10^-8^, Tukey post-hoc test; p < 0.05 for pairs 4 vs. 2 and 3 vs. 2; p < 0.01 for pairs 5 vs. 2 and 3 vs. 1; P < 0.001 for pairs 4 vs. 3, 4 vs. 5 and 5 vs. 1; fig. 3N, IPSC 1 amplitude). Conversely, IPSC1 failure rates were low in pairs 4 and 5 (0.05 and 0.1 respectively), and higher in pairs 2 and 3 (0.55 and 0.33 respectively; fig. 3N, IPSC1 FR). All pairs, except pair 2 showed paired-pulse depression of eIPSC amplitude, the PPR being 0.66, 0.6 and 0.56 for pairs 3, 4 and 5 respectively (fig. 3N, PPR), indicating a decrease of release probability from AP1 to AP2. In contrast, in pair two, where the presynaptic input was a burst of APs, the delay between the first and second APs within the burst varied around 10 ms. Here we measured paired-pulse facilitation, from 16.2 ± 3.8 pA (n = 20, including failures) of IPSC1 to 32.2 ±1.9 pA (n = 20, including failures) of IPSC2 (fig. 3E, G). This paired-pulse facilitation was accompanied by a decrease in failure rate of IPSC2 compared to IPSC1 from 0.55 to 0.5 (fig. 3G). The increased release probability at AP2 is likely explained by the short (∼ 10 ms) delay between the first two APs in the burst and accumulation of intracellular calcium.

We also analysed the response kinetics of these connections. The largest response delay (from AP peak to IPSC onset) was in pair 2 (2.5 ± 0.2 ms, n = 9 IPSCs, omitting failures; fig. 3F), and the shortest in pair 3 (0.6 ± 0.1 ms, n = 8 IPSCs, omitting failures; fig. 3M); IPSC1 delay in pairs 1, 4 and 5 was 1.2 ± 0.1 (n = 15), 1.6 ± 0.05 (n = 19) and 1.8 ± 0.1 (n = 18) ms respectively (One-way ANOVA, F = 34.75, p = 2.1 * 10^-15^, Tukey post-hoc test, p < 0.05 for all comparisons). The mean risetime (20-80 %) was the shortest in pair 4 with 0.36 ± 0.039 ms (n = 19). Pairs 3 and 5 had a risetime of 0.61 ± 0.1 (n = 8) and 0.67 ± 0.1 (n = 18 IPSCs, no failures) ms respectively. In pair 1, the risetime was 0.87 ± 0.058 ms (n = 15 IPSCS, no failures) (One-way ANOVA, F = 7.37, p = 3 * 10^-4^, Tukey post-hoc test, p < 0.05 for pair 5 vs. 1; p < 0.001 for pair 4 vs. 1; fig. 3N, 20-80 risetime). Finally, we found no difference in half width between the four connections (pair 2 excluded due to burst) (One-way ANOVA, F = 0.47, p = 0.707; fig. 3N, half-width).

In summary, the most reliable connection was made between DBC 28 and DBC 27 (pair 4), and this synapse also showed the fastest kinetics. Connections made by CB+ DBCs to interneurons were less reliable than other connections. Finally, the connection by the CB+ DBC onto the pyramidal cell was also reliable, but showed slower kinetics than the connection in pair 4.

### Axonal and dendritic features and molecular characterisation of PV-DTCs

We have fully recovered 14 (12 cells in temporal and 2 cells in the frontal cortex) cells with large, multipolar, smooth dendritic trees and extensive axon spreading uniformly in all directions from the soma (fig. 4A). The cell bodies of these cells were located in layer III at an average distance of ∼ 1000 ± 430 μm (n = 3) from the pia. The 6 – 8 main dendrites travelled radially on average to ∼ 375 ± 37 μm (n = 4) from the soma and had a relatively straight course (fig. 4A). The axons of these cells were even more extensive and travelled to ∼ 500 ± 97 μm on average uniformly in every direction (fig. 4A). Importantly, the axons did not project to layer I and formed mostly *en-passant* and only few *en-termineux* boutons, which occasionally contacted neuronal somata (fig. 4G).

Neurons with similar dendritic and axonal distributions have been described previously in humans and other species. One such interneuron type is the *large basket cell*. Basket cells innervate the somata and the proximal dendrites of pyramidal neurons, as well as dendritic spines (Somogyi et al., 1983; Kisvárday et al., 1987; Kisvárday et al., 1990; Kisvárday et al., 1993). However, cells with similar dendritic trees and axonal distribution have also been found innervating mostly dendrites, including spines and no cell bodies (Tamás et al., 1997a). Furthermore, basket cells have been demonstrated previously to be of at least two types in rodents, which can be differentiated based on immunoreactivity to PV or CCK (Freund et al., 1983; Freund et al., 1986; Kawaguchi & Kubota, 1998; Somogyi et al., 2004; Armstrong & Soltész, 2012). Those that are immunopositive for CCK have been found to be also immunopositive for the cannabinoid type 1 receptor (CB1R) along the axonal membrane in the hippocampus (Katona et al., 1999). Thus, we tested our multipolar cells for immunoreactivity to PV and CB1. We also tested them for CB, as this was previously found to be co-expressed with PV or CCK in interneurons (Gulyás et al., 1991; Kubota et al., 1994). Nine out of 12 tested multipolar cells were immunopositive for PV as shown either in their cell body (n = 2) or boutons (n = 10; fig. 4D, E, H), whereas none of the 5 PV+, 3 PV-, and one PV-untested cells were immunopositive for CB1 along their axons (fig. 4H). Eight of the nine PV+ multipolar cells were immunonegative for CB in their boutons (n = 5) or soma (n = 3, fig. 4E); one was immunopositive in the soma. Two PV-multipolar cells tested were also immunonegative for CB in their boutons. The boutons of two PV+ and one PV-multipolar cell were tested for VIP and were immunonegative. The boutons of all 11 multipolar cells tested were immunopositive for VGAT (fig. 4F, H).

The expression of PV by the majority of the multipolar cells and their axonal and dendritic structures are characteristic of neocortical *large basket cells*, and *dendrite-targeting cells* described in other species (Hartwich et al., 2009). However, because we found the majority of our multipolar cell synapses innervating dendrites and only a few synapses targeting somata (see later), instead of naming them basket cells, which might be misleading, we refer to them as *parvalbumin-expressing dendrite-targeting cells* (PV-DTC).

### Intrinsic membrane properties of PV-DTCs in comparison with DBCs

Based on recording quality, the current-voltage responses of 11 of the PV-DTCs were analysed (table 3). Several active and passive membrane properties differed between PV-DTCs and DBCs; PV-DTCs showed a smaller amplitude voltage sag in response to hyperpolarising current injection than DBCs, and lacked rebound spikes after the hyperpolarising steps (fig. 4 B, fig. 5 A, B, C, sag ratio: PV-DTCs 1.11 ± 0.06 vs. DBCs 2.07 ± 0.58, Welch’s t-test, p = 1.6 x 10^-9^). Only the PV+/CB+ double immunopositive PV-DTC 7 fired rebound spikes. During continuous depolarisation, PV-DTCs fired a regular train of APs with less spike amplitude accommodation than DBCs (fig. 5 D; PV-DTCs 0.85 ± 0.07 vs. DBCs 0.57 ± 0.12, Student’s t-test, p = 2.6 x 10^-8^); spike frequency adaptation was similar in the two cell types (fig. 5 E; PV-DTCs median: 2.45, IQR: 1.83, DBCs median: 3.43, IQR: 3.52, Kolmogorov-Smirnov test, p = 0.074). Two PV-DTCs developed depolarisation block at RB + 100 pA current injection. In contrast to DBCs, only one cell, PV-DTC 2, responded with a burst of 4 APs, all other PV-DTCs fired 1 or 2 action potentials to RB current (e.g. fig. 4B, PV-DTC 1 and fig. 5A, PV-DTC 9). The mean firing rate of PV-DTCs at RB +100 pA current injection was similar to that of DBCs (PV-DTCs 36.9 ± 18.3 Hz vs. DBCs 38.1 ± 19.7 Hz, Student’s t-test, p = 0.87), but the regularity of the AP trains were different, as the coefficient of variation of the inter-spike intervals (ISI CV) was smaller in PV-DTCs than DBCs (PV-DTCs median: 0.2, IQR: 0.08 vs. DBCs median: 0.33, IQR: 0.33, Kolmogorov-Smirnov test, p = 0.021).

**Figure 5:**
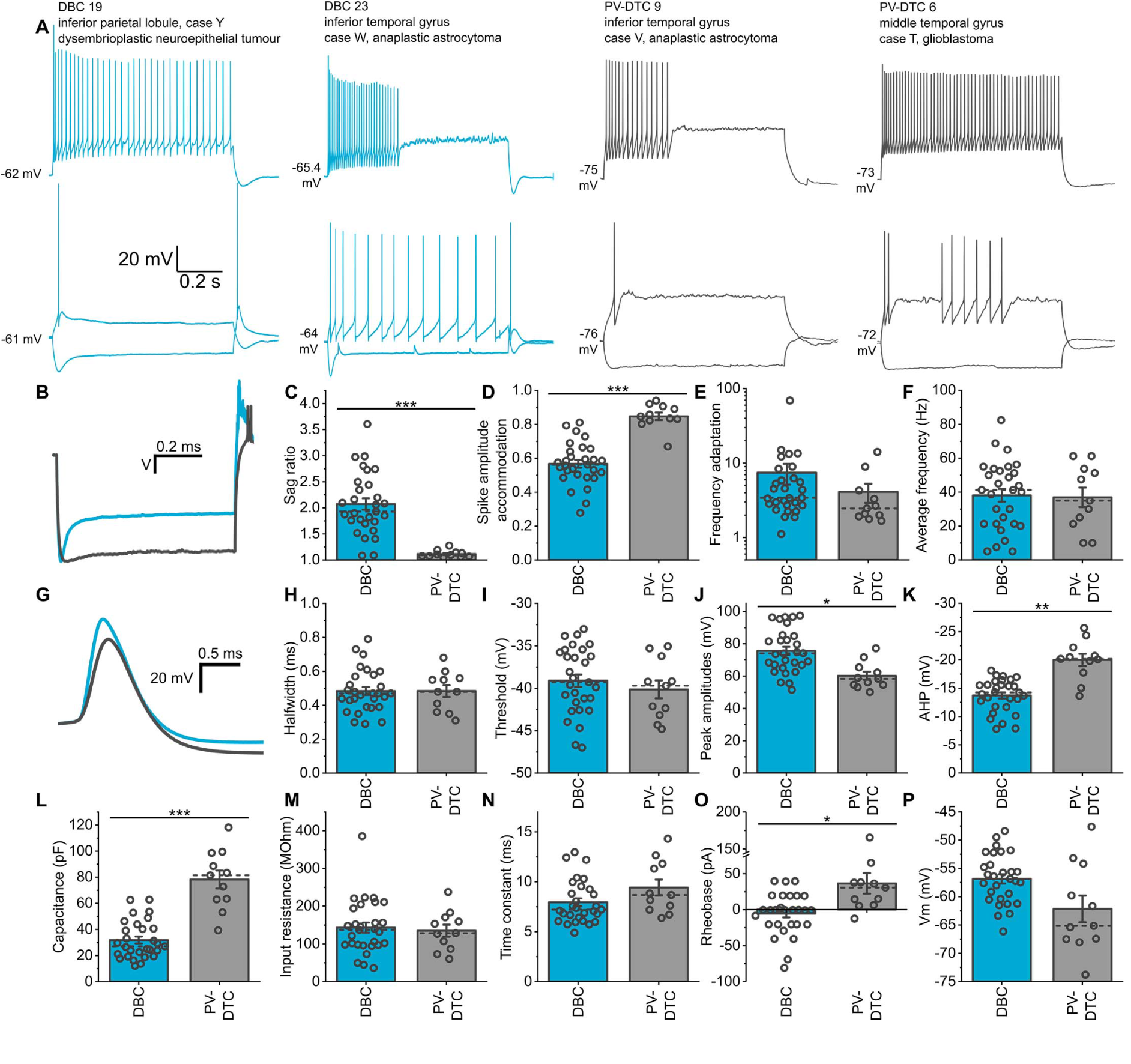
Comparison of membrane properties of DBCs and PV-DTCs. **A.** Example traces from two DBCs (blue) and two PV-DTCs (grey) showing voltage responses to current injections of rheobase, holding current -100 pA and rheobase +100 pA. **B.** Average waveforms of the voltage responses of the DBCs (blue, n = 29) and PV-DTCs (grey, n = 11) to holding current - 100 pA current injection. The average traces were normalised such that the initial baseline values and the lowest points on the voltage responses are equal. **C.** Ratios of the sags in the voltage responses to stimulation with holding current -100 pA, calculated as the peak voltage deflection divided by the amplitude of the steady state voltage response (Welch’s t-test, t_(29.9)_ = 8.55, p = 1.6 x 10^-9^). **D.** Spike amplitude accommodation ratios, calculated by dividing the amplitude of the last action potential by the amplitude of the first generated in response to rheobase +100 pA current injection (Student’s t-test, t_(38)_ = -6.98, p = 2.6 x 10^-8^). **E.** Firing frequency adaptation ratios calculated by dividing the last inter-spike interval with the first one of the response to rheobase +100 pA current injection (Kolmogorov-Smirnov test, D = 0.429, Z = 1.21, p = 0.074). **F.** Average frequencies of action potentials generated in an 800 ms response to rheobase +100 pA current injection (Student’s t-test, t_(38)_ = 0.16, p = 0.87). **G.** Average waveforms of the first action potentials generated at rheobase current injection for each cell. **H - J.** Halfwidths (**H**, Student’s t-test, t_(38)_ = -0.008, p = 0.99), thresholds (**I**, Student’s t-test, t_(38)_ = 0.76, p = 0.45), and amplitudes (**J**, Student’s t-test, t_(38)_ = 3.5, p = 0.001), of the first action potential generated in response to rheobase current injection for each cell. **K.** Afterhyperpolarisation amplitudes of the first action potential for each cell at rheobase current injection, calculated relative to the threshold of that action potential (Student’s t-test, t_(38)_ = 5.71, p = 1.4 x 10^-6^). **L.** Whole cell capacitances, calculated from the voltage amplitude and the time constant associated with the slowest component of a double exponential fit to the initial voltage deflections of the responses to holding current -60 pA current injection (Student’s t-test, t_(38)_ = -7.69, p = 2.9 x 10^-9^). **M.** Comparison of input resistances (Student’s t-test, t_(38)_ = 0.34, p = 0.733). **N.** Membrane time constants, measured as the slowest component of the double exponential fit described in **l** (Student’s t-test, t_(38)_ = -1.82, p = 0.077). **O.** The amount of current required to elicit at least one action potential within an 800 ms period of stimulation for each cell (Welch’s t-test, t_(12.89)_ = -2.74, p = 0.017). Note that 20 pA current injection steps were used, and the rheobase was taken as the holding current of the first step to elicit an action potential. **P.** The membrane voltage of each cell recorded in response to a 0 pA current injection, or estimated based on the cell’s current-voltage relationship if no 0 pA step was recorded (Welch’s t-test, t_(12.49)_ = 2.13, p = 0.053).

**Figure 6.**
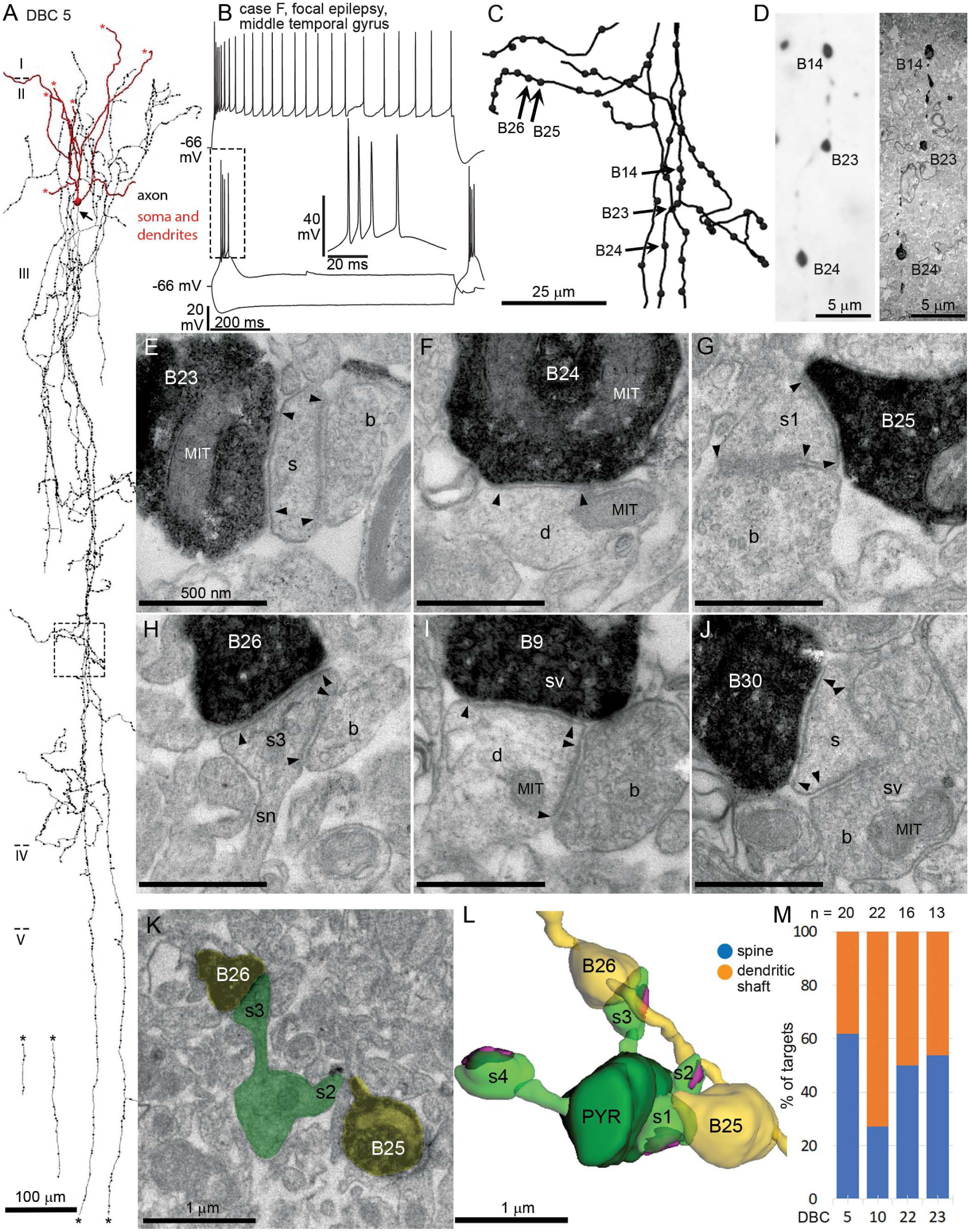
Synaptic targets of DBCs in layer III of the human neocortex. **A.** Partial 3D reconstruction (3 sections of ∼ 60 μm thickness) of DBC5 with dendrites entering layer I, and a “horsetail” axon (arrow, initial segment) descending to layer V (asterisks indicate cut ends of axon and dendrites); note short lateral collaterals in layer III. **B.** Voltage responses of the cell to current injections (+60, +160 and -100 pA steps) in whole-cell patch-clamp; inset showing the burst in boxed area. **C.** Enlarged view of the boxed area in A, included in serial electron microscopic analysis; numbered boutons are shown in E-H, K-L. **D. Left:** Light micrograph of a descending axon collateral with three boutons visualised by HRP reaction product (single optical section within a ∼60 μm thick osmium treated section); **Right:** low power electron micrograph of the same boutons in a 50 nm thick resin section. **E-J.** High power electron micrographs of the boutons of the cell in A forming type-2 synapses with their targets. The edges of synapses are indicated by black arrowheads. Note the smooth apposition of the pre- and postsynaptic membranes and synaptic vesicles (sv) at the pre- synaptic active zones. Boutons 23, 25, 26 and 30 innervate dendritic spines (s), whereas Boutons 9 and 24 innervate dendritic shafts (d). All targets, except the dendrite in F are innervated by additional boutons (b) forming type-1 synapses (arrowheads). MIT: mitochondrion; sn: spine neck. **K - L.** electron micrograph (K) and 3D reconstruction (L) of the axon with two boutons ( B25, B26, yellow) and their target spiny pyramidal cell dendrite (PYR, green). The target of B25 (s1) is a stubby spine (see G), that of B26 (s3) is a mushroom spine (see H). Two other spines were also identified (2, 4). All four spines receive a type 1 synapses (magenta). **M.** Proportion of synaptic targets (n, shown on top) of four CB+ DBCs (cell numbers as in Fig. 1).

Parvalbumin-expressing basket cells in the cortex of other species are also referred to as fast-spiking basket cells, because of their short duration action potentials (AP) and high frequency firing *in vivo* and *in vitro* (McCormick et al., 1985; Baranyi et al., 1993; Kawaguchi, 1995; Erisir et al., 1999). In our sample PV-DTCs had a mean AP half-width of 0.48 ± 0.12 ms, similar to that of DBCs (fig. 5 G, H; 0.48 ± 0.11 ms, Student’s t-test, p = 0.99). Under our conditions, the AP threshold of PV-DTCs was not different from that of DBCs (fig. 5 I; PV-DTCs -40.1 ± 3.4 mV vs. DBCs -39.1 ± 3.8 mV, Student’s t-test, p = 0.45), but on average PV-DTCs had smaller AP amplitude than DBCs (fig. 5 J; PV-DTCs 60.2 ± 7.9 vs. DBCs 75.6 ± 13.3 mV, Student’s t-test, p = 0.001) and larger AHP amplitude (fig. 5 K; PV-DTCs 20 ± 3.4 mV vs. DBCs 13.7 ± 2.9 mV, Student’s t-test, p = 1.4 x 10^-6^).

PV-DTCs had larger whole-cell capacitance than DBCs (fig. 5 L; PV-DTCs 78.3 ± 21.8 pF vs. DBCs 31.9 ± 14.1 pF, Student’s t-test, p = 2.9 x 10^-9^), reflecting the larger cell bodies and longer dendrites of PV-DTCs compared to DBCs. The input resistance (IR) and membrane time constant of PV-DTCs was not different from those of DBCs (fig. 5 M IR; PV-DTCs 135 ± 50 MΩ vs. DBCs 143 ± 70 MΩ, Student’s t-test, p = 0.733; fig. 5 N tau; PV-DTCs 9.4 ± 2.5 ms vs. DBCs 7.9 ± 2.1 ms, Student’s t-test, p = 0.077). Furthermore, PV-DTCs had larger rheobase than DBCs (fig. 5 O; PV-DTCs 36.5 ± 45.5 pA vs. DBCs -5.5 ± 28.4 pA, Welch’s t-test, p = 0.017), whereas their resting membrane potentials were similar (fig. 5 P; PV-DTCs -62.2 ± 7.4 mV vs DBCs 56.9 ± 4.3 mV, Welch’s t-test, p = 0.053).

### Synaptic targets of DBCs and PV-DTCs

In non-human species DBCs innervate both pyramidal cells and other interneurons (Somogyi & Cowey, 1981; DeFelipe et al., 1989; DeFelipe & Jones, 1992; Tamás et al., 1997a), in agreement with our paired IPSC recordings in humans. The connectivity rate of 25 % for interneurons in the vicinity of DBCs may be an underestimate due to dendritic filtering of small IPSCs. We could not record a representative sample of DBC-to-PYR connections to estimate the connectivity rate. Furthermore, because our paired recordings were restricted to the vicinity of the DBC soma in layers II-III, we don’t know if these interactions were mediated by the “horsetail” of DBCs or the more loosely distributed axon collaterals in the vicinity of the DBC somata.

In order to explore the synaptic targets of the “horsetail” axons, we used serial section electron microscopy of individual DBCs’ axonal boutons and compared their distributions to the synapses made by PV-DTCs, which have a very different axonal shape. We studied the “horsetail” axons of 4 CB+ DBCs (5, 10, 22 and 23), and one CR+ DBC (25). All samples were taken from layer III and included small collaterals (fig. 6A). All CB+ DBCs examined were recorded in the temporal association cortex, whereas the CR+ DBC was recorded in the inferior parietal lobule. One DBC (5) was recorded in a sample from an epilepsy patient, and the rest from tumour patients (see table 2).

Synaptic junctions were identified based on well-established electron microscopic criteria. These included the electron opaque synaptic cleft, a regular and uniform apposition of the presynaptic and postsynaptic membranes, the slight widening of the extracellular space, accumulation of synaptic vesicles at the pre-synaptic active zone and the presence of a varying width of postsynaptic density (fig. 6E – J). We examined a total of 132 boutons (20 - 34 from each DBC), of which 100 were found to form synaptic junctions. For 32 boutons the synaptic junctions could not be identified because of an incomplete series of sections or because of the unfavourable cutting angle relative to the synaptic membranes. Three boutons formed two separate synaptic junctions each, all other boutons (97 %) formed just one synapse, thus the total number of identified synaptic junctions was 103. In 9 instances the synaptic junction was identified, but the small diameter target structure could not be defined, so a total of 94 synaptic targets were identified. All synapses made by DBCs were of Gray’s type II, or symmetric (Gray, 1959; Colonnier, 1968) with a thin if any postsynaptic density (fig. 6E - J), consistent with the GABAergic nature of DBCs shown here. The shape and size of vesicles could not be studied in the labelled boutons, being covered by the electron opaque DAB/OsO4 precipitate (fig. 6E - J). All examined synaptic junctions were made with dendritic spines or dendritic shafts; no synapse was found on soma. Small diameter processes of triangular or elongated shapes and without mitochondria were identified as spines if the process could be followed to its end in one direction through serial sections (e.g. fig. 6E, G, H, J; total no. of spines: 39). Some spines (33 %) contained a spine apparatus, a membranous structure with multiple cisternae (fig. 6H). All dendritic spine targets were also innervated by an unlabelled type I (asymmetric) synapse (fig. 6E, G, H, J). We followed spine targets through serial sections in an attempt to identify the parent dendritic shaft to predict its origin from an interneuron or a pyramidal cell. Due to loss of sections and the very thin and long shape of some of the spine necks, only 9 spines could be traced back to their parent shafts. Three of these dendritic shafts emitted spines other than that innervated by the DBC and none of the dendrites received type I synapses on their shaft, thus most likely belonged to pyramidal cells. Nevertheless, given the abundance of dendritic spines on pyramidal cells relative to interneurons, it is likely that the majority of spine targets belong to pyramidal cells. In the case of the CB+ DBC 5, one postsynaptic dendrite was innervated by two labelled boutons, both of them targeting spines. These spines also received a type I synapse each. Two other spines on the same target dendrite received a single type I synapse each (fig. 6K, L).

Dendritic shafts postsynaptic to DBCs could not be followed to an end in either direction and contained mitochondria and/or microtubules amongst other organelles (fig. 6F, I). We identified a total of 55 dendritic shaft synaptic targets and followed them through serial sections. Of these 10 had at least 4 (average: 6.4 ± 2.6) type I and type II additional synapses covering their surface densely. As no spines originated from these dendrites, we identified them as belonging to GABAergic interneurons. We grouped 23 dendritic shaft targets with at least 3 innervating synapses (including the labelled synapse) and lacking spines as originating from GABAergic interneurons. On 32 dendritic shaft targets, either no, or only one additional synapse could be found due to short and incomplete series of sections; 3 of these also emitted spines. These three dendrites were grouped as pyramidal cell dendrites, whereas 29 postsynaptic dendritic shafts remained unidentified. The average diameter of the target shafts was of median 623, IQR: 454 nm diameter, and some were of very small diameter (e.g. fig. 6 F), corresponding to distal dendrites.

Among the CB+ DBCs the average proportion of dendritic spine targets was 48 ± 15 % (Fig. 6 M; DBCs 5, 10, 22 and 23, n = 23, 22, 18 and 13 respectively), the remainder (52 ± 14 %) being dendritic shafts. The CB-/CR+ DBC 25 had the lowest proportion (17 %) of dendritic spine targets (3/18), the majority of postsynaptic elements (83 %) being dendritic shafts. The frequency distribution of synaptic targets was different between all DBCs, including the CB-/CR+ DBC 25 (*X*_2 (4, N = 94)_ = 11.31, p = 0.023), meaning that the proportion of shaft and spine targets depends on the identity of the DBCs. When looking at only the CB+DBCs no such dependence was observed (*X*_2 (3, N = 76)_ = 5.51, p = 0.138). Among the targets of CB+ DBCs identified as originating from GABAergic interneurons or pyramidal cells, the average proportion of pyramidal cell targets was 76.5 ± 18 %, whereas that of identified interneuron targets was 23.5 ± 18 % (DBCs 5, 10, 22 and 23, N = 17, 11, 11 and 6 respectively). On the other hand, the proportion of pyramidal cell targets for the CB-/CR+ DBC No 25 was only 21.4 %, the majority of the targets (78.6 %) being interneurons. These synaptic target distributions are similar to what was reported in other species (Somogyi & Cowey, 1981; DeFelipe et al., 1989; DeFelipe & Jones, 1992; Tamás et al., 1997a), but the homology of cell types is difficult to establish (see discussion).

We also studied a total 104 boutons from 3 PV-DTCs (3, 4 and 5), and identified 68 synaptic junctions with 63 identified postsynaptic structures (fig. 7). In the case of 36 boutons the synapse could not be found due to incomplete series of sections or unfavourable cutting angle; in the case of 5 synapses the postsynaptic structure could not be defined. The PV-DTCs 3 and 4 were recorded in the temporal cortex, whereas PV-DTC 5 in the frontal cortex. All samples came from tumour patients. All synapses made by PV-DTCs were of Gray’s type II (fig. 7 F - K), but in contrast to DBCs, PV-DTCs also targeted somata in addition to dendrites. Based on the presence of the nucleus (fig. 7 E) or on the size of the profile, a total of 8 targets were identified as soma; 4 of the somata were identified as pyramidal cells, based on the presence of only additional type I synapses. We identified 41 targets as dendritic shafts (fig. 7 G, H, J), of which 5 were identified as interneuron dendrites (fig. 7 H), and 36 as pyramidal cell dendrites (fig. 7 G, J), based on the same criteria as above. Finally, 13 targets were identified as dendritic spines (fig. 7 K), of which 6 contained a spine apparatus. One soma was found to be innervated by two boutons of a single PV-DTC. Interestingly, PV-DTC 4 had a higher proportion of spine targets (40 %) compared to the other two (13.3 and 12.5 % for PV-DTC 3 and 5 respectively; fig. 7 E). The average frequency of soma targets was 12.8 ± 3.4 % for PV-DTCs, whereas that of dendritic shafts and spines was 65.2 ± 13.9 % and 21.9 ± 16.7 % respectively. When comparing this with the target frequency distributions of CB+ DBCs, we found that there was significant correlation between the cell type and the frequency of different synaptic targets (*X*_2 (12, N = 126)_ = 28.066, p = 0.005), thus CB+ DBCs and PV-DTCs have different synaptic target preferences in the human neocortex. Although PV-DTCs innervate somata to some extent, the majority of their synapses are onto dendrites, similarly to DBCs. To test whether different dendritic domains are innervated by these two cell types, we measured the diameters of target dendrites. We found no difference in the diameter of target dendrites innervated by DBCs and PV-DTCs (DBCs median: 623, IQR: 454 nm vs. PV-DTCs median: 726, IQR: 550 nm, Mann-Whitney U-test U = 378, Z = -1.43, p = 0.153), suggesting that DBCs and PV-DTCs innervate overlapping dendritic domains.

**Figure 7.**
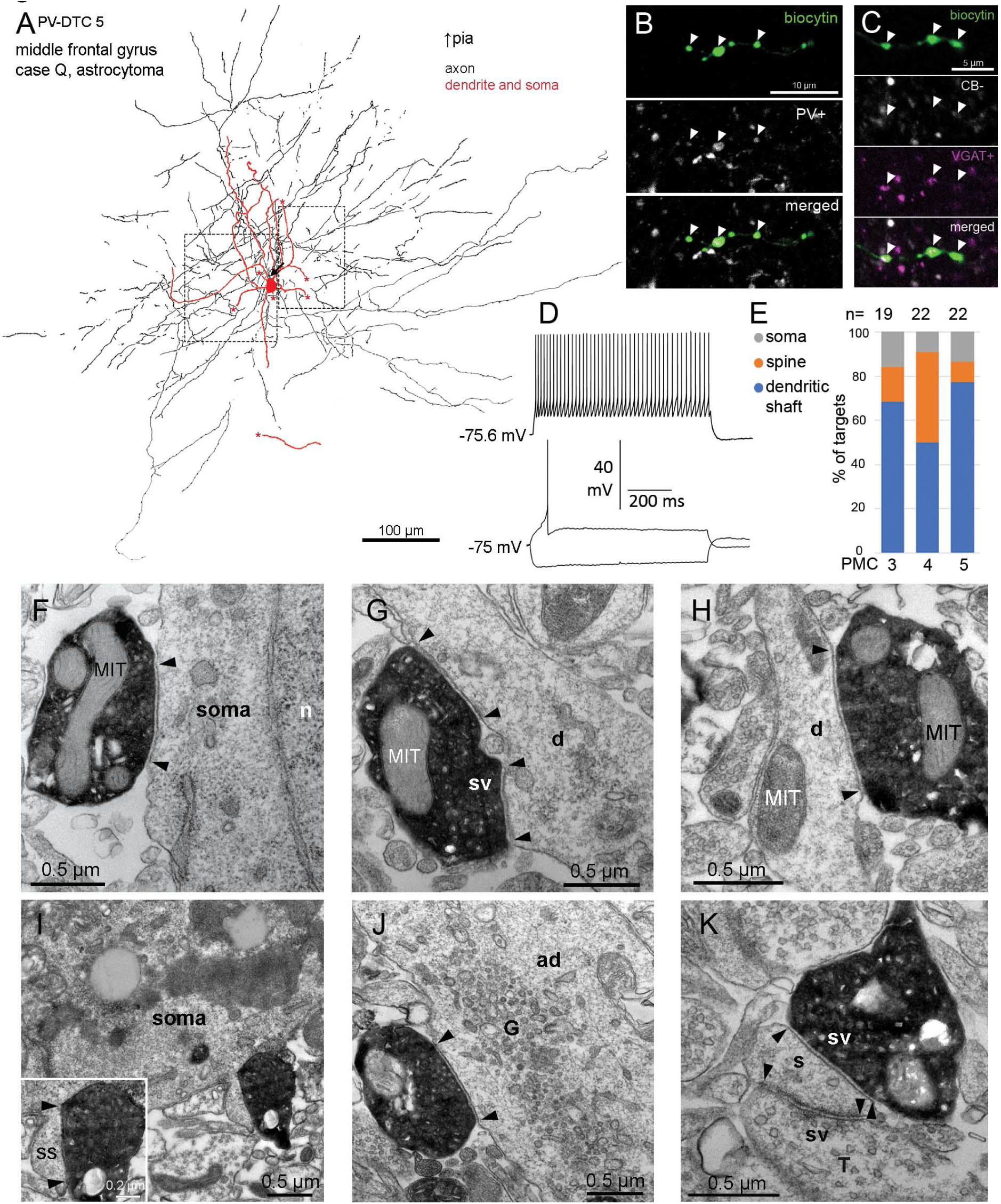
Synaptic targets of PV-DTCs in layer III of the human neocortex. **A.** Partial 2D reconstruction from 2 sections of ∼60 μm thickness of PV-DTC 5 with dendrites (red) and axons (black) extending in every direction uniformly around the cell body in layer III. The axon initial segment (arrow), cut ends of dendrites (asterisks) and the direction of the pial surface are indicated. Boxes indicate the areas re-embedded for electron-microscopy. **B.** Boutons (arrowheads) of the cell in **A** are immunopositive for PV (white) (single optical slice of 0.4 μm thickness). **C.** Boutons (arrowheads) of the cell are immunopositive for vesicular GABA transporter (VGAT, magenta) and immunonegative for CB (white) (single optical slice of 0.4 μm thickness). **D.** Voltage responses of the cell held in current-clamp mode to current injections of rheobase, holding current -100 pA and rheobase +100 pA. **E.** Proportion of synaptic targets for PV-DTCs 3, 4 and 5 (n = 19, 22 and 22 respectively), as identified by serial section electron microscopy. **F-K.** Electron micrographs of labelled boutons (black, electron opaque) of PV-DTC 5 (**F-H**) and PV-DTC 4 (**I-K**), forming type II synaptic junctions with somata (**E, I),** inset in **I** shows the bouton innervating a somatic spine (ss), dendritic shafts (d, ad; **G, H, J**), and a dendritic spine (s, **K**). Edges of the synaptic junctions are indicated by arrowheads. The spine in **K** also receives a type I synapse (arrowheads) from an unlabelled terminal (T). n: nucleus, MIT: mitochondrion, d: dendritic shaft, ad: apical dendrite; ss: somatic spine, sv: synaptic vesicles, G: Golgi apparatus.

**Figure 8.**
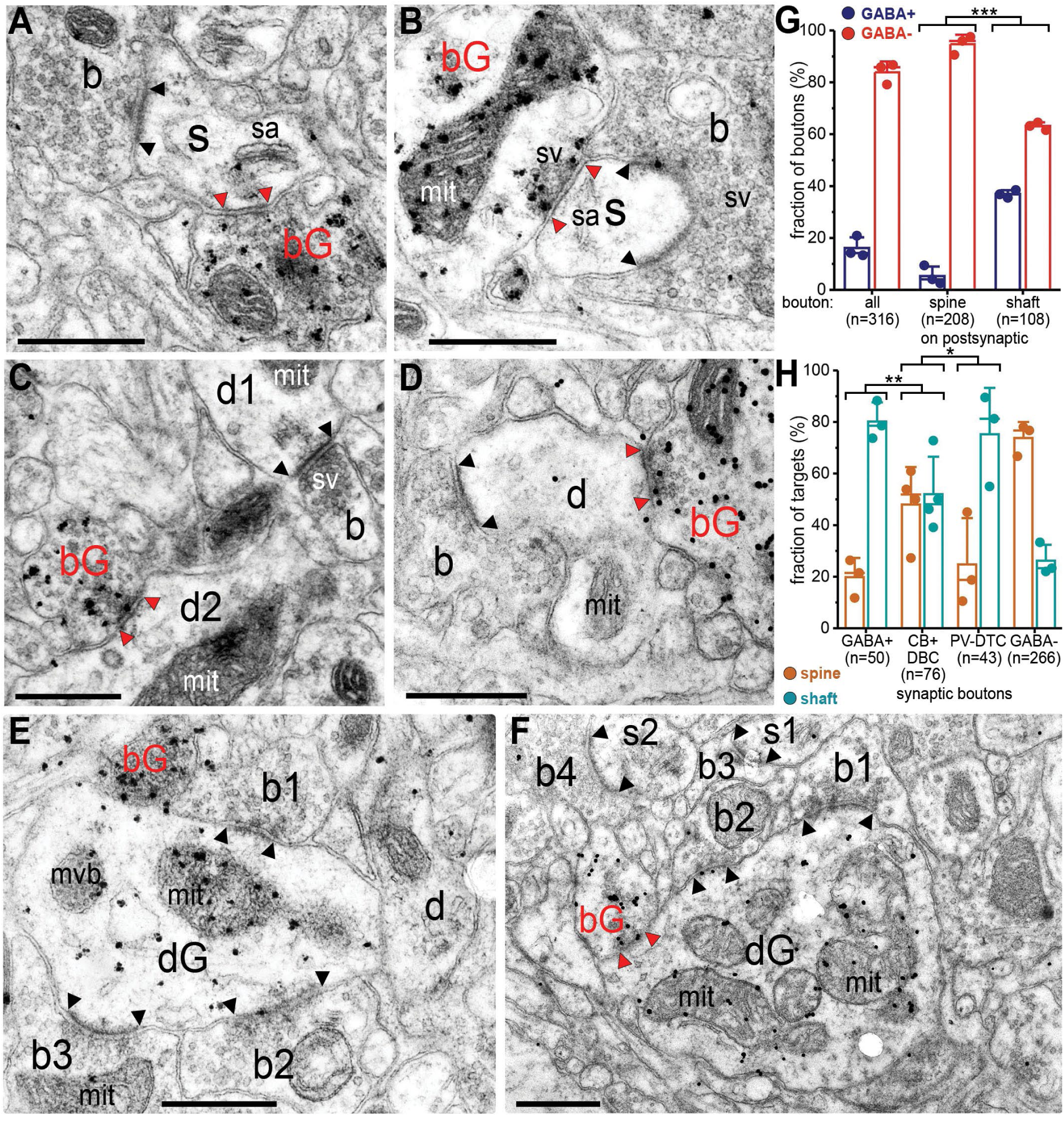
Synaptic targets of GABAergic and non-GABAergic synapses in the neuropil. **A-F** Electron micrographs of sections immunolabelled for GABA with silver-enhanced colloidal gold particles (electron opaque dots). Gold particles are enriched over the surface of GABA-positive boutons (bG) and some dendrites (dG). **A-B.** Double innervation of dendritic spines (s) with spine apparatus (sa) by a GABA+ bouton (bG) and another bouton not labelled for GABA (b). Some synaptic vesicles (sv) are indicated in the presynaptic boutons. **C.** A GABA+ bouton (bG) innervating a GABA-negative dendritic shaft (d2), identified by the presence of a mitochondrion (mit). A GABA-negative bouton (b) innervates a different GABA-negative dendritic shaft (d1). **D.** A GABA+ (bG) and a GABA-negative bouton (b) innervate the same GABA-negative dendritic shaft (d). **E.** Three GABA-negative boutons (b1-3) form type I synaptic junctions with a GABA+ dendritic shaft containing a multivesicular body (mvb); another GABA+ bouton (bG) is also in contact with the dendrite but a synaptic junction is not visible. A different dendrite not labelled for GABA (d) is present. **F.** Two GABA-negative (b1-2) and a GABA+ bouton (bG) innervate a GABA+ dendritic shaft (dG). Two other GABA-negative boutons (b3-4) innervate a dendritic spine each (s1-2). Note the large number of mitochondria present in the target GABA+ dendrite. The synaptic specialisations are indicated between arrowheads (red for GABAergic, black for non-GABAergic synapses). Scale bars in A-F, 0.5 μm. **G.** Fraction of GABA+ and GABA-negative (GABA-) synaptic boutons innervating dendritic spines or shafts (*X^2^*_(1, 316)_ = 55.44, p = 9.6*10^-14^ between spine and shaft targeting synapses). **H.** Fraction of dendritic spine and shaft synaptic targets innervated by different populations of presynaptic boutons, including those from physiologically recorded and labelled DBCs and PV-DTCs (*X^2^*_(1, 126)_ = 9.75, p = 0.0018 between GABA+ and DBC boutons; *X^2^*_(1, 119)_ = 5.46, p = 0.0195 between DBC and PV-DTC boutons). Bars: mean, whiskers: one standard deviation, line: median, number of boutons are indicated, individual data points are shown.

We conclude that CB+ DBCs have different synaptic target preference from the CB-DBC and PV-DTCs in the human neocortex, and dedicate a high proportion of their synapses to dendritic spines of pyramidal cells. While PV-DTCs also innervate dendritic spines, the majority of their targets are dendritic shafts and they also innervate somata, which DBCs omit. The dendritic domains targeted by DBCs and PV-DTCs overlap.

### Targets of GABAergic synapses in the neuropil of layer III

*Double bouquet cells* and PV-DTCs significantly differ in their output postsynaptic target distributions. In order to compare their selection of output synapse locations to the distribution of potential GABAergic synaptic target sites in the neuropil, we have studied the distribution of GABAergic synapses on dendritic spines and shafts in layer III. Using post-embedding immunogold reactions with antibodies to GABA in electron-microscopic sections (Somogyi & Soltész, 1986) and a stereological method based on 3D tracing of synapses and their targets in a defined volume through serial sections (West, 1999; Cano-Astorga et al., 2021) we determined the numerical density and the proportion of GABAergic synapses and their neuropil targets in samples from temporal and frontal cortices.

Boutons were identified as GABAergic based on the enrichment of silver-intensified gold particles over their cross-sectional surface consistently from section to section (fig. 8 A-F). Enrichment of gold particles was present over mitochondria in GABAergic postsynaptic dendrites (fig. 8 E, F) as well, but this signal was usually weaker than in boutons. Boutons making type I synapses, dendritic spines, spiny dendrites and most dendritic shafts only had a low level if any particles which we considered technical background (fig. 8 A-D). Boutons immunopositive for GABA (GABA+) established typical Gray’s type II synapses with thin post-synaptic densities and targeted mostly dendritic shafts (fig. 8 C-F) and less frequently dendritic spines (fig. 8 A, B). All but two of the GABA+ bouton innervated dendritic spines received an additional type I synapse from a GABA-negative bouton. Most boutons not labelled for GABA (GABA-) established typical type I synapses with thick post-synaptic densities (fig. 8 A-F) and targeted mostly dendritic spines.

We counted a total of 316 synapses in a volume of 252.5 μm^3^ of temporal cortex (n=4 3D probes, case AA) and 176.4 μm^3^ and 238.5 μm^3^ of frontal cortices (n=5 3D probes, case BB; n=2 3D probes, case CC; volumes are corrected for tissue compression along the cutting direction, but not for tissue shrinkage during dehydration). The mean numerical density of synapses across all three samples was 0.48 ± 0.04 per μm^3^ of neuropil, which excluded cell bodies, and appeared lower in the temporal (0.42) than in the frontal (0.52 and 0.50) cortex. The fraction of synapses made by GABA+ boutons was 16.1 ± 3.4 % (fig. 8 G; mean density: 0.08 ± 0.02 per μm^3^; n=3 patients). Dendritic spines were the targets of the majority of synapses (65.4 ± 5.2 %), and only 5.3 ± 3 % of these were made by GABA+ boutons (fig. 8 G). The proportion of spines receiving double innervation by one GABA+ and one GABA-bouton to those receiving at least one GABA-synapse was 4.6 ± 2.7 %. In contrast, the proportion of GABA+ boutons among those targeting dendritic shafts was significantly higher (36.9 ± 1.2 %; fig. 8 G; *X^2^*_(1, 316)_ = 55.44, p = 9.6*10^-14^ for the proportion of GABA+ boutons among those targeting dendritic spines vs. those targeting dendritic shafts). Amongst the dendritic shafts that received synapses, 9.3 ± 5.4 % were GABA-immunopositive. Finally, the majority of GABA+ boutons (80.2 ± 6 %) targeted dendritic shafts (fig. 8 H), whereas the majority of boutons not labelled for GABA (73.8 ± 5.1 %) targeted dendritic spines (fig. 8 H).

There was a significant association between the proportion of dendritic spine to dendritic shaft targets and the different populations of synaptic bouton (*X^2^*_(3, 435)_ = 81.49, p = 0, fig. 8 H). Specifically, a significantly higher proportion of dendritic spine targets was found among DBC boutons than among all GABA+ boutons (*X^2^*_(1, 126)_ = 9.75, p = 0.0018) or among PV-DTC boutons that targeted spines or shafts (excluding those innervating somata) (*X^2^*_(1, 119)_ = 5.46, p = 0.0195). The proportion of dendritic spine targets was not different between PV-DTC boutons and all GABAergic boutons (*X^2^*_(1, 93)_ = 0.41, p = 0.521). Taken together, we have observed that DBCs but not PV-DTCs selectively target dendritic spines as postsynaptic elements.

### Regulation of GABAergic inputs to DBCs and PV-DTCs by group III mGluRs

We have shown that in the human neocortex two distinct types of GABAergic interneuron, double-bouquet cells and basket cells, have distinct axonal and dendritic distributions and innervate different subcellular compartments of their target cells. Furthermore, differences in the dendritic distribution and passive and active membrane properties between DBCs and PV-DTCs suggest that they receive and process synaptic inputs differently. The synaptic inputs of DBCs are not known. Group III metabotropic glutamate receptors (mGluRs) are pre-synaptic regulators of both GABAergic and glutamatergic synaptic transmission in many distinct forebrain neuronal circuits in rodents. Synaptic pathway- and cell type-specific expression of group III mGluRs in the rodent cortex (Shigemoto et al., 1996; Dalezios et al., 2002; Somogyi et al., 2003; Ferraguti et al., 2005) is thought to result in the differential inhibition of GABAergic synaptic transmission to different types of interneuron and pyramidal cells (Semyanov & Kullmann, 2000; Kogo et al., 2004). However, neither the subcellular localisation nor the functions of group III mGluRs are known in the human neocortex.

We tested if GABAergic synaptic transmission to different types of interneuron in the human neocortex is differentially regulated by group III mGluRs. Specifically, we tested if activation of group III mGluRs by the orthosteric agonist *L-2-amino-4-phosphonobutyric acid* (L-AP4), depresses the frequency and/or amplitude of GABA_A_ receptor mediated spontaneous inhibitory postsynaptic currents (sIPSCs), as has been shown in rodents (Mitchell & Silver, 2000; Giustizieri et al., 2005; Cuomo et al., 2009). It is important to note, that action potentials and fast glutamatergic synaptic transmission were not blocked in these experiments.

We recorded sIPSCs in whole-cell voltage clamp, at just below firing threshold potential (- 50 to -45 mV), using a low Cl^-^ (4 mM) concentration in our internal pipette solution. In this setting sIPSCs presented as upward deflections in the holding current with a fast rising phase and an exponential decay (fig. 1C and 4C). In this experimental configuration we also recorded AMPA receptor mediated spontaneous excitatory postsynaptic currents (sEPSCs, fig. 1C), but these were not analysed. We recorded sIPSCs in a total of 185 neurons and tested the effects of L-AP4 on 133, of which 65 were analysed, including 49 putative interneurons and 16 putative pyramidal cells. The remainder 68 cells did not pass technical criteria to be included in the study (see methods). After microscopic examination of the visualised and pharmacologically tested cells, we identified 7 DBCs (see fig. 1), 7 PV-DTCs (see fig. 5), 2 neurogliaform cells, 1 rosehip cell (Boldog et al., 2018), and 1 axo-axonic cell, based on the specific distribution of their axon. The identity of 29 cells could not be determined, and 2 cells were not recovered at all. Four pyramidal cells were confirmed visually, based on their densely spiny dendrites. We compared the effects of L-AP4 on sIPSCs in DBCs and PV-DTCs as these two groups had enough identified cells to study the effects at the population level (fig. 9). To control for spontaneous rundown of synaptic activity *in vitro*, and variability in slice conditions and technical parameters, the effects were compared to vehicle-treated control cells (n = 7), including 4 interneurons and 3 pyramidal cells. Median sIPSC frequency was 2.0 Hz (IQR: 3.6 Hz, n = 18) in interneurons and 1.6 Hz (IQR: 1.3 Hz, n = 16) in pyramidal cells, whereas median amplitude was 9.0 pA (IQR: 5.9 pA) in interneurons and 12.3 pA (IQR: 5.7 pA) in pyramidal cells.

**Figure 9.**
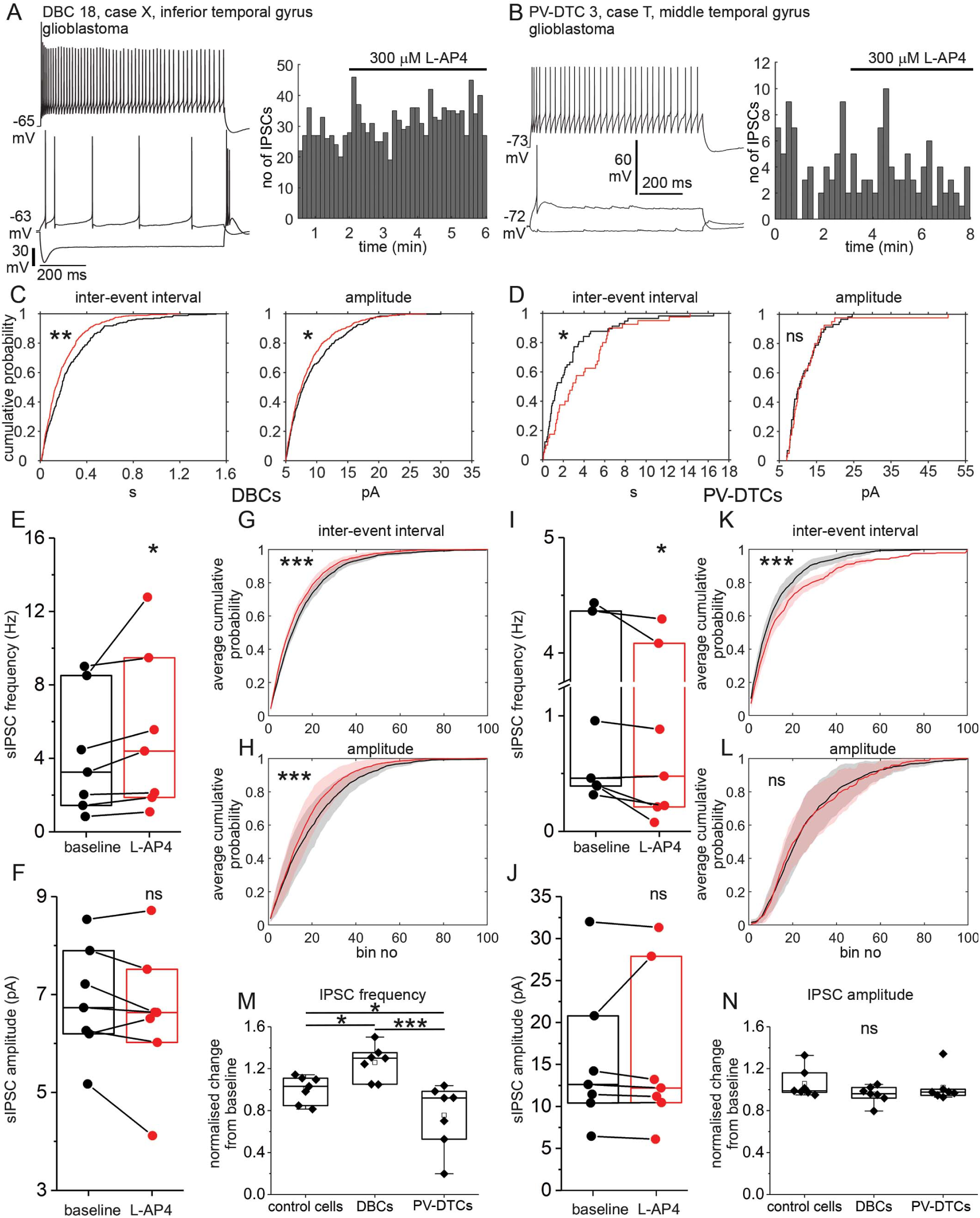
Effects of group III mGluR activation on sIPSCs in DBCs and PV-DTCs. **A.** Effect of group III mGluR activation on a DBC. Left: Voltage responses of DBC 18 to current injections of RB, RB +100 pA, and holding current -100 pA; right: event time histogram of sIPSCs recorded from DBC 18 during baseline and bath application of 300 μM L-AP4 (n = 1326 events, bin width: 8 s). **B.** Effect of group III mGluR activation on a PV-DTC. Left: Voltage responses of PV-DTC 3 to current injections of RB, RB +200 pA and holding current -100 pA; right: event time histogram of sIPSCs recorded from PV-DTC 3 during baseline and bath application of 300 μM L-AP4 (n = 140 events, bin width: 10 s). **C.** Cumulative probability distribution of sIPSC inter-event intervals (left) and amplitudes (right) in baseline (black, n = 326) vs. L-AP4 (red, n = 528) conditions in DBC 18 (Kolmogorov-Smirnov test, p = 0.001 for intervals and p = 0.0466 for amplitudes). **D.** same as C for PV-DTC 3 (baseline, black, n = 59 vs. L-AP4, red, n = 41; Kolmogorov-Smirnov test, p = 0.048 for intervals and p = 0.876 for amplitudes). **E.** Mean sIPSC frequency of DBCs in baseline vs. L-AP4 conditions (paired sample Wilcoxon signed rank test, p = 0.016, boxes represent median and IQR). **F.** Median sIPSC amplitude of all DBCs in baseline vs. L-AP4 conditions (paired sample Wilcoxon signed rank test, p = 0.219). **G.** Mean cumulative probability distribution (line) and 95 % CI (shaded areas) of sIPSC inter-event intervals across all DBCs in baseline (black) vs. L-AP4 (red) conditions (100 bins/cell, Kolmogorov-Smironov test, p = 4.8 * 10^-5^). **H.** Average cumulative probability distribution of sIPSC amplitudes across all DBCs (100 bins/cell, Kolmogorov-Smironov test, p = 3.2 * 10^-4^). **I-L.** Same as E-H for PV-DTCs (I, J: paired sample Wilcoxon signed rank test, p = 0.031 for intervals, p = 0.375 for amplitudes; K,L: Kolmogorov-Smirnov test, p = 5.7 * 10^-10^ for intervals and p = 0.099 for amplitudes). **M-N.** Normalised change from baseline to L-AP4 in sIPSC frequency (M) and amplitude (N) in control cells (n = 7), DBCs (n = 7), and PV-DTCs (n = 7) (Kruskal-Wallis ANOVA test, for frequency, p = 0.002, Conover post hoc test, adjusted by the Holm FWER method, p = 0.011 for controls vs DBCs, p = 0.043 for controls vs PV-DTCs, p = 0.0001 for DBCs vs PV-DTCs; for amplitude, p = 0.382, boxes represent median and IQR, square symbol, mean, whiskers, SD).

In three out of 7 DBCs, bath application of L-AP4 (50 μM, n = 3; or 300 μM, n = 4) unexpectedly shifted the cumulative distribution of sIPSC inter-event intervals to the left (fig. 9C left, two sample Kolmogorov-Smirnov test, p = 0.001, n = 854 IPSCs), meaning that the proportion of shorter inter-event intervals increased compared to longer ones, corresponding to an increase in the frequency of sIPSCs. An increase in the number of sIPSCs after application of L-AP4 can also be seen on the event time histogram for DBC 18 (fig. 9A). In the other 4 DBCs, the inter-event interval did not change. However, the leftward shift in the distribution of sIPSC intervals was still present when averaged across all DBCs (fig. 9G, two sample Kolmogorov-Smirnov test, p = 4.8 * 10^-5^, n = 100 bins). At the population level, the frequency of sIPSCs increased from median 3.25 Hz (IQR: 7.08 Hz) at baseline to median 4.4 Hz (IQR: 7.6 Hz; Wilcoxon signed rank test, p = 0.022, n = 7; fig. 9E). The increase in frequency was not accompanied by a change in amplitude (fig. 9F, baseline median: 6.7, IQR: 1.7 pA vs. L-AP4 median: 6.6, IQR: 1.5 pA, Wilcoxon signed rank test, p = 0.219), although in individual DBCs L-AP4 shifted the cumulative distribution of sIPSC amplitudes to the left (smaller amplitudes, e.g. fig. 9C right, two sample Kolmogorov-Smirnov test, p = 0.047, n = 854 IPSCs) in 4 cells; to the right (larger amplitudes) in one, and did not change in the remaining two. The average cumulative distribution of sIPSC amplitudes also shifted towards smaller amplitudes (fig. 9H, two sample Kolmogorov-Smirnov test, p = 3.2 * 10^-4^, n = 100 bins). Washout of L-AP4 from the bath did not lead to the recovery of the baseline frequencies or amplitudes. These results are in contrast with previous studies in rodents, where L-AP4 suppressed spontaneous and evoked IPSCs (Gerber et al., 2000; Matsui & Kita, 2003; Drew & Vaughan, 2004; Lu, 2007; Cuomo et al., 2009).

In contrast to DBCs, in PV-DTCs bath application of 300 μM L-AP4 shifted the cumulative distribution of sIPSC inter-event intervals to the right, corresponding to lower sIPSC frequency (e.g. fig. 9D left, two sample Kolmogorv-Smirnov test, p = 0.048, n = 100 IPSCs) in 2 out of 7 cells, also seen as a decrease in the number of sIPSCs after application of L-AP4 in the event time histogram of PV-DTC 5 (fig. 9B). In 5 cells the distribution did not change. However, averaged across all PV-DTCs the distribution of sIPSC intervals shifted to the right, upon application of L-AP4 (fig. 9K, two sample Kolmogorov-Smirnov test, p = 5.7 * 10^-10^, n = 100 bins). At the population level, although the median frequency increased from 0.46 Hz (IQR: 4 Hz) in baseline to 0.48 Hz (IQR: 3.9 Hz), the pairwise decrease was significant (fig. 9I, Wilcoxon signed rank test, p = 0.031). The average cumulative distribution of sIPSC amplitudes did not change upon application of L-AP4 (fig. 9L, two sample Kolmogorov-Smirnov test, p = 0.099, n = 100 bins), nor did the median amplitude at the population level (fig. 9J, baseline median: 12.6, IQR: 10.4 pA vs. L-AP4 median: 12.2, IQR: 17.4 pA, Wilcoxon signed rank test, p = 0.375, n = 7). These effects are in agreement with results in rodents (Gerber et al., 2000; Matsui & Kita, 2003; Drew & Vaughan, 2004; Lu, 2007; Cuomo et al., 2009).

We further assessed whether the changes measured upon L-AP4 application are different from spontaneous changes in frequency and amplitude due to changing slice conditions, technical parameters, and synaptic rundown. We compared the baseline normalised changes measured in DBCs and PV-DTCs to the changes measured in vehicle treated control cells (fig. 9M, N). The median change in sIPSC frequency was to 103 % (IQR: 26 %) of baseline in vehicle treated controls (n = 7). This was significantly smaller than the change in DBCs (median: 130 %, IQR: 30 %, Conover post hoc test, adjusted by the Holm FWER method, p = 0.011 for controls vs DBCs), and significantly larger than the change observed in PV-DTCs (median: 92 %, IQR: 46 %, Conover post hoc test, adjusted by the Holm FWER method, p = 0.043) (fig. 9M). The changes were also different between DBCs and PV-DTCs (Conover post hoc test, adjusted by the Holm FWER method, p = 0.0001). Conversely, normalised change in sIPSC amplitude in DBCs and PV-DTCs did not differ from the change in vehicle treated controls (fig. 9N, Kruskal-Wallis test, p = 0.382). These comparisons show that L-AP4 alters the frequency rather than on the amplitude of sIPSCs, which indicates that L-AP4 primarily acts through modulation of neurotransmitter release at the level of axon terminals, in accordance with previous studies in rodents (Gerber et al., 2000; Semyanov & Kullmann, 2000; Matsui & Kita, 2003; Drew & Vaughan, 2004; Kogo et al., 2004; Cuomo et al., 2009). However, in our experiments action potentials and fast glutamatergic synaptic transmission were not blocked, thus such conclusions could not be drawn, and the unexpected increase of sIPSC frequency in DBCs is likely the result of a complex network effect, by group III mGluRs acting at both GABAergic and glutamatergic synapses and multiple levels of synaptic connections, resulting in the disinhibition of a GABAergic synaptic input specific to DBCs. Nevertheless, these results show a cell type-specific modulation of GABAergic synaptic input to two distinct types of GABAergic neuron in the human neocortex.

### Analysis of the effects of group III mGluR activation on sIPSC frequency with respect to patient related variables

We showed differential effects of group III mGluR activation on sIPSCs in two different GABAergic neuron types on a small number of neurons which were recorded in samples of different cortical regions from patients with various different diagnosis and medical history. Samples from tumour patients were taken as far from the MRI-imaged tumour mass as possible and in deep tumours cortical tissue was not affected by infiltration, as assessed by MRI. However, some samples showed some degree of infiltration and/or oedema (table 2). Furthermore, chemical signals may not be limited to structures directly infiltrated by the tumour mass. Particularly, seizures associated with brain tumours and epilepsy can spread through the entire neocortex. Because group III mGluRs have been associated previously with epilepsy and seizures in rodents, we tested if the effects of group III mGluR activation on sIPSC frequency measured here, correlated with the clinical symptoms of patients, such as the presence or absence of seizures, antiepileptic medication, or other pathological processes related to sample quality, such as the presence of infiltration or oedema. We also tested if the measured effects depended on the cortical region where the sample was taken from and the age and sex of the patients (see table 2).

We generated several multiple linear regression models, which included different sets of independent variables, in order to find the ones which best explain the variability in the data. However, due to the low number of pharmacologically tested neurons (n = 14 cells, 7 DBCs, 7 PV-DTCs) and the uneven frequency of the different cortical areas and pathologies, the predictive value of these models is limited. Almost all pharmacological response data came from patients between 50 – 66 years of age with the exception of one outlier (age: 31 years; mean: 57.5 ± 2.4 yrs.). Furthermore, 9 (64 %) cells were recorded in samples from glioblastoma patients, and none in samples from TLE patients. Similarly, 11 cells (79 %) were recorded in the temporal, 2 (14 %) in the frontal and only one (7 %) in the parietal cortex. Of the 7 models generated (table 4), models 1 (Adj. R^2^: 0.47, F = 4.79, p = 0.026), 4 (Adj. R^2^: 0.75, F = 6.44, p = 0.019), 5 (Adj. R^2^: 0.83, F = 8.51, p = 0.027), 6 (Adj. R^2^: 0.88, F = 14.74, p = 0.002) and 7 (Adj. R^2^: 0.47, F = 3.93, p = 0.04) were significantly different from a constant. All models included the cell identity as a factor. Model 1 included the age and sex in addition to the cell identity, model 4 included the age, sex, cortical region, pathology, the presence of oedema in the sample, and seizures, and model 5 included all parameters, except hypertension as factors. Model 6 included age, sex, the presence of oedema, seizures and treatments with anticonvulsive and steroid medication, but not the cortical region and the pathology, whereas model 7 only included age, sex and hypertension in addition to the cell identity (table 4). In models 1 and 7 the cell identity was the only factor significantly different from zero, contributing to the pharmacological response, however these models explained less than half of the variance in the data (table 4). In model 5, seizures were a significant factor in addition to the cell identity. Interestingly, in model 4 all factors, except age and pathology were significant. In model 6, in addition to the cell identity and age, seizures and anticonvulsive medication were significant factors with seizures adding to the increase in sIPSC frequency induced by L-AP4 (0.67 ± 0.14, p = 0.003), and anticonvulsive treatment having the opposite effect (-0.6 ± 0.17, p = 0.013; table 4). Model 6 also explained the most of the variability in the change in sIPSC frequency (Adj. R^2^: 0.88). In all the different models, the most consistently significant factor was the cell identity, however seizures were also significant in three models. Although these models support that the effect of group III mGluR activation on sIPSC frequency is cell type-specific, considering the small number of data points, it cannot be excluded that other factors, particularly seizures and antiepileptic medication might also have an effect on the measured responses.

**Table 4.**
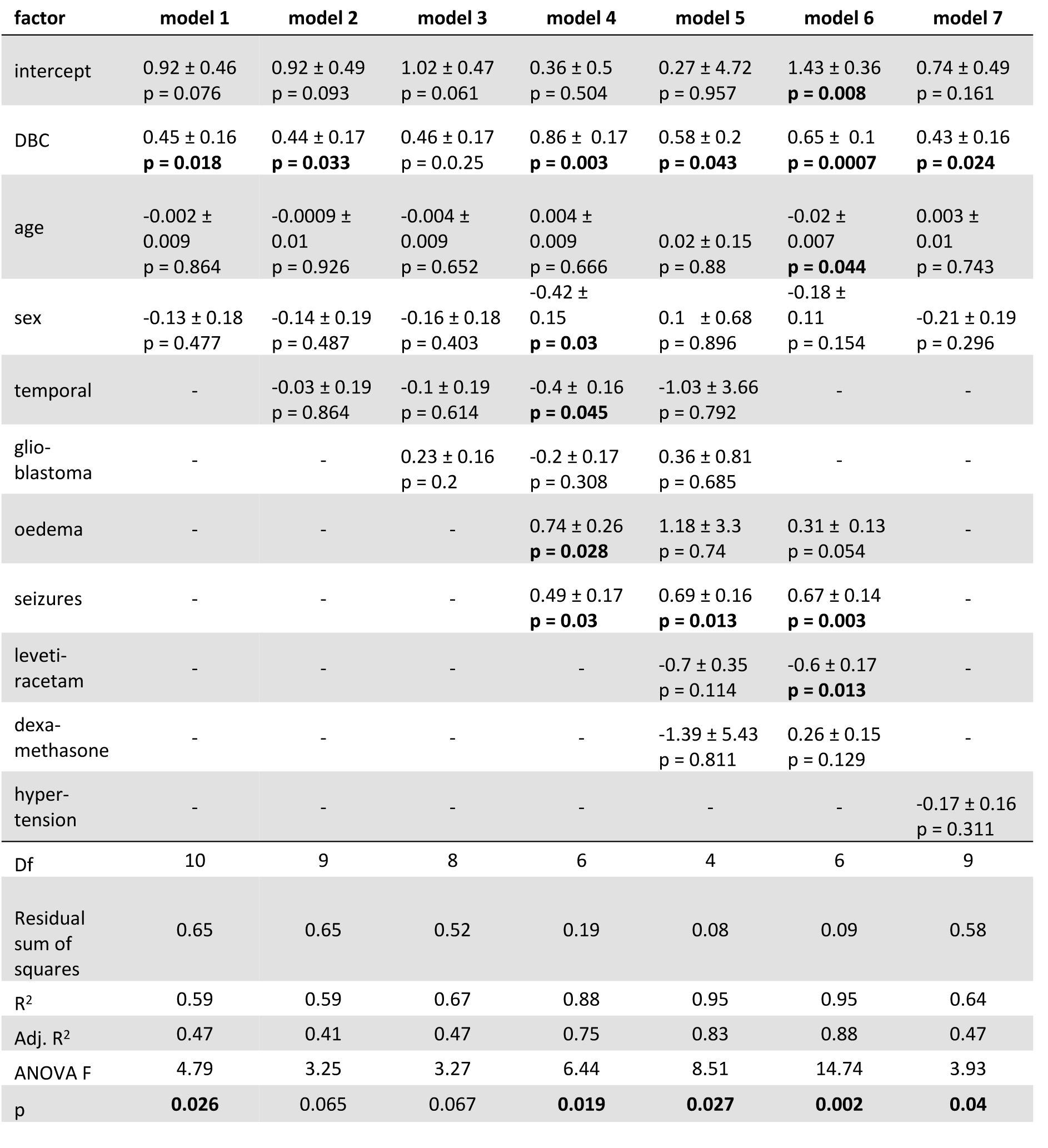
Multiple linear regression models for the effect of group III mGluR activation on sIPSC frequency.

| factor | model 1 | model 2 | model 3 | model 4 | model 5 | model 6 | model 7 |
| --- | --- | --- | --- | --- | --- | --- | --- |
| intercept | 0.92 ± 0.46<br>p = 0.076 | 0.92 ± 0.49<br>p = 0.093 | 1.02 ± 0.47<br>p = 0.061 | 0.36 ± 0.5<br>p = 0.504 | 0.27 ± 4.72<br>p = 0.957 | 1.43 ± 0.36<br>**p = 0.008** | 0.74 ± 0.49<br>p = 0.161 |
| DBC | 0.45 ± 0.16<br>**p = 0.018** | 0.44 ± 0.17<br>**p = 0.033** | 0.46 ± 0.17<br>p = 0.025 | 0.86 ± 0.17<br>**p = 0.003** | 0.58 ± 0.2<br>**p = 0.043** | 0.65 ± 0.1<br>**p = 0.0007** | 0.43 ± 0.16<br>**p = 0.024** |
| age | -0.002 ± 0.009<br>p = 0.864 | -0.0009 ± 0.01<br>p = 0.926 | -0.004 ± 0.009<br>p = 0.652 | 0.004 ± 0.009<br>p = 0.666 | 0.02 ± 0.15<br>p = 0.88 | -0.02 ± 0.007<br>**p = 0.044** | 0.003 ± 0.01<br>p = 0.743 |
| sex | -0.13 ± 0.18<br>p = 0.477 | -0.14 ± 0.19<br>p = 0.487 | -0.16 ± 0.18<br>p = 0.403 | -0.42 ± 0.15<br>**p = 0.03** | 0.1 ± 0.68<br>p = 0.896 | -0.18 ± 0.11<br>p = 0.154 | -0.21 ± 0.19<br>p = 0.296 |
| temporal | - | -0.03 ± 0.19<br>p = 0.864 | -0.1 ± 0.19<br>p = 0.614 | -0.4 ± 0.16<br>**p = 0.045** | -1.03 ± 3.66<br>p = 0.792 | - | - |
| glio-blastoma | - | - | 0.23 ± 0.16<br>p = 0.2 | -0.2 ± 0.17<br>p = 0.308 | 0.36 ± 0.81<br>p = 0.685 | - | - |
| oedema | - | - | - | 0.74 ± 0.26<br>**p = 0.028** | 1.18 ± 3.3<br>p = 0.74 | 0.31 ± 0.13<br>p = 0.054 | - |
| seizures | - | - | - | 0.49 ± 0.17<br>**p = 0.03** | 0.69 ± 0.16<br>**p = 0.013** | 0.67 ± 0.14<br>**p = 0.003** | - |
| leveti-racetam | - | - | - | - | -0.7 ± 0.35<br>p = 0.114 | -0.6 ± 0.17<br>**p = 0.013** | - |
| dexa-methasone | - | - | - | - | -1.39 ± 5.43<br>p = 0.811 | 0.26 ± 0.15<br>p = 0.129 | - |
| hyper-tension | - | - | - | - | - | - | -0.17 ± 0.16<br>p = 0.311 |
| Df | 10 | 9 | 8 | 6 | 4 | 6 | 9 |
| Residual sum of squares | 0.65 | 0.65 | 0.52 | 0.19 | 0.08 | 0.09 | 0.58 |
| R<sup>2</sup> | 0.59 | 0.59 | 0.67 | 0.88 | 0.95 | 0.95 | 0.64 |
| Adj. R<sup>2</sup> | 0.47 | 0.41 | 0.47 | 0.75 | 0.83 | 0.88 | 0.47 |
| ANOVA F | 4.79 | 3.25 | 3.27 | 6.44 | 8.51 | 14.74 | 3.93 |
| p | **0.026** | 0.065 | 0.067 | **0.019** | **0.027** | **0.002** | **0.04** |

For each independent variable (factor) its coefficient is given as mean ± SE and the p-values are listed. ANOVA statistics for each model are given at the bottom.

## Discussion

Diverse types of local interneurons with specialised roles, as reflected in their distinct molecular expression, intrinsic biophysical properties and synaptic relationships provide most of the GABAergic innervation of the cerebral cortex (Kawaguchi & Kubota, 1997; Somogyi et al., 1998; Gupta et al., 2000; Defelipe et al., 2013; Hodge et al., 2019). We have characterised two distinct GABAergic interneuron types in human association neocortex, which differ in calcium-binding protein expression, intrinsic membrane properties, synaptic output and the group III mGluR-mediated regulation of their GABAergic inputs. The DBCs are a cell type in the upper layers of the neocortex of primates with a striking columnar “horsetail” axon (Ramón y Cajal, 1899; Jones, 1975; Valverde, 1978; Somogyi & Cowey, 1981; DeFelipe et al., 1989; DeFelipe et al., 1990; Lund & Wu, 1997) with as yet no clear homologue in rodents (Ballesteros Yáñez et al., 2005). In contrast, PV-DTCs with widely distributed axons resembling neurons described as ‘basket cells’ have been illustrated in rodents (Kawaguchi & Kubota, 1998; Kawaguchi & Kondo, 2002; Packer & Yuste, 2011), carnivores (Meyer, 1983; Somogyi et al., 1983; Meyer & Ferres-Torres, 1984; Kisvárday et al., 1987) and primates (Hendry et al., 1989; Lund & Lewis, 1993; Gabbott & Bacon, 1996a; Lund & Wu, 1997) including humans (Marin-Padilla & Stibitz, 1974; Kisvárday et al., 1990; Szegedi et al., 2017; Bakken et al., 2021). In terms of synaptic outputs, human CB-expressing DBCs innervate dendrites but not somata and preferentially target dendritic spines. In contrast, PV-DTCs target mainly dendritic shafts and to a lesser extent somata and dendritic spines. These two cell types also differ in the response of their GABAergic inputs to the activation of group III mGluRs, presynaptic receptors that inhibit GABA and glutamate release in rodents (Mitchell & Silver, 2000; Semyanov & Kullmann, 2000; Kogo et al., 2004).

### Molecular diversity of cortical GABAergic neurons

Amongst the many molecular differences of GABAergic interneurons, the selective expression of calcium binding proteins characterises families of cell types in rodents (Cauli et al., 1997; Kubota et al., 2011; Rudy et al., 2011; Tasic et al., 2018; Yao et al., 2021) and primates (Lund & Lewis, 1993; Condé et al., 1994; Gabbott & Bacon, 1996b; Zaitsev et al., 2005), including humans (DeFelipe, 1997; González-Albo et al., 2001; Varga et al., 2015; Hodge et al., 2019). Each calcium binding protein is expressed by several distinct cell types and cell types may express more than one calcium binding protein, hence on their own they are insufficient to delineate functionally defined cell types. In our recorded interneuron population, CB and CR were found in separate individual DBCs, which were all immunonegative for PV. Conversely, PV-DTCs were positive for PV and negative for CB and CR, except for one cell, which was double immunopositive for PV and CB. The differential expression of PV, CB and CR in the interneurons we studied is in agreement with the mutually exclusive expression of CB and PV in the neocortex of monkeys (DeFelipe et al., 1989; Gabbott & Bacon, 1996a), but different from rodent neocortex, where a large population of GABAergic neurons including basket cells co-expresses CB and PV (Kubota & Jones, 1993; Cauli et al., 1997; Kawaguchi & Kubota, 1997; Hartwich et al., 2009). In humans CB and PV are co-expressed in axo-axonic cells (del Río & DeFelipe, 1997a).

Transcriptomic analysis indicates 4 main GABAergic neuronal types in the middle temporal gyrus (MTG)(Hodge et al., 2019) and 4 in the primary motor cortex (M1)(Bakken et al., 2021) expressing mRNA for CB (CALB1), based on median RNA counts. In the MTG, among those groups with cell bodies in layers II and III where DBCs are located the two largest groups either co-express somatostatin (*Inh L1-3 SST CALB1*) or CCK (*Inh L2-6 LAMP5 CA1*), although the latter group is more frequently found in layers V and VI. In M1, GABAergic neuron clusters with the highest expression of CALB1 and cell bodies in layers II and III are *Inh L1-2 SST CCNJL*, *Inh L1-2 PRRT4* and *Inh L1-2 SST CLIC6*, all of which co-express high levels of SST (Bakken et al., 2021). However, we only found one CB+ DBC to be immunoreactive for somatostatin, and none immunoreactive for CCK. Although in the cat visual cortex, CCK was suggested to be expressed in DBC-like GABAergic neurons (Freund et al., 1986), it is unclear if those were homologous to the DBCs in humans. Furthermore, patch-seq data indicates that transcriptomic cell types in mice corresponding to these human clusters mainly include upper layer Martinotti cells (Hodge et al., 2019; Callaway et al., 2021). In M1, the cluster *Inh L5-6 LAMP5 CRABP1* co-expresses CB and CCK, however, the homologous mouse cluster *LAMP5 LHX6* corresponds to neurogliaform cells (Callaway et al., 2021). Finally, in the MTG the CALB1 expressing cluster *Inh L1-6 PVALB SCUBE3* homologous to the mouse clusters of axo-axonic cells (Hodge et al., 2019) expresses CB together with PV. The corresponding cluster in M1 (*Inh L1-6 PVALB COL15A1*) does not express CALB1 (Callaway et al., 2021). In accordance with the co-expression of CB with PV in one PV-DTC described here, scattered expression of CALB1 is also found in the human MTG cluster *Inh L2-4 PVALB WFDC2*, which is homologous to clusters including fast-spiking basket cells in mice (Hodge et al., 2019; Callaway et al., 2021). In summary, we are unable to allocate a clear transcriptomic cell type to DBCs at present. Considering their high abundance in humans, it is possible that they are vulnerable to cell sorting and are not represented in their real proportion in transcriptomic datasets. Alternatively, they may be included in the numerous *Inh L1-3 SST CALB1* group mixed with Martinotti cells, which have a very different axon. Our inability to detect somatostatin immunoreactivity in DBCs could be due to a genuine lack or very low levels of peptide expression. The same may apply to *Inh L2-6 LAMP5 CA1* having CCK peptide levels undetectable with our methods. Future patch-seq data in human cortex would clarify this issue.

Regarding PV-DTCs, in human MTG and M1 there are 2 and 4 PV-expressing transcriptomic cell types, respectively, with cell bodies in layers II and III (Hodge et al., 2019; Bakken et al., 2021). Of these the *Inh L2-4 PVALB WFDC2* in MTG, and the *Inh L2-5 PVALB RPH3AL* in M1, are homologous to mouse transcriptomic cell types thought to include superficial-layer basket cells (Hodge et al., 2019; Gouwens et al., 2020; Callaway et al., 2021). However, because the synaptic target preference of the individual neurons mapped onto these transcriptomic clusters was not tested (Gouwens et al., 2020; Bakken et al., 2021), it is not known whether dendrite-targeting cells comparable to the PV-DTCs described here are represented in the same or different transcriptomic clusters.

The visualisation of individual CB-positive DBCs proves that they contribute a major component of the CB immunoreactive radial axon bundles in humans and probably also in monkeys (DeFelipe et al., 1989; DeFelipe et al., 1990; Ferrer et al., 1992; del Río & Defelipe, 1997b; Ballesteros Yáñez et al., 2005; DeFelipe et al., 2006). Importantly, however, a radial axon bundle likely contains axons from multiple DBCs, including CB-immunonegative and CR-immunoreactive ones, and other CB-expressing interneuron types may also contribute. In many cortical areas supragranular pyramidal cells express CB (Gabbott & Bacon, 1996a; Kawaguchi & Kubota, 1997), thus their axons may also be included. In carnivores immunoreaction to CB also reveals axon bundles (Ballesteros Yáñez et al., 2005), but it is not clear if DBCs with horse tail axons contribute. Bundles of CB-positive axons have not been documented in rodents, lagomorphs and ungulates (Ballesteros Yáñez et al., 2005). Calretinin-immunoreactive (CR+) bundles of axon were reported in humans (del Río & Defelipe, 1997b), but individual DBCs have not been previously visualised. Two of our recorded DBCs were immunopositive for CR and immunonegative for CB, but because of the small number of CR+ DBCs, we could not compare their features to the CB+ population. The available evidence from synaptic output of CR+ interneurons suggests that multiple cell types exist, some of which may specialise in innervating other interneurons (del Río & DeFelipe, 1997c; Gabbott et al., 1997; Meskenaite, 1997), in agreement with the synaptic target preference of the one CR+ DBC reported here and in contrast to the spine-preferring CB+ DBCs.

It is not yet clear whether DBCs are present in rodents, or if they are, whether they express CB. Neurons with descending axons, visualised and named DBCs in rats, do not have comparable “horsetail” axons to those in primates and express VIP, corticotropin releasing factor (CRF) and CR (Kawaguchi & Kubota, 1996, 1997; Karube et al., 2004; Kubota et al., 2007). In transcriptomic studies VIP is co-expressed with CB in only one group of cells (*L1-2 VIP WNT4*) in human M1 (Hodge et al., 2019; Bakken et al., 2021)(Allen Institute for Brain Science (2021). Allen Cell Types Database -- Human Multiple Cortical Areas [dataset]. Available from celltypes.brain-map.org/rnaseq). Interneurons with descending “horsetail” axons and unknown molecular profiles were described in mice (Jiang et al., 2013; Jiang et al., 2015), but another similar large scale study did not report such neurons (Gouwens et al., 2020). Furthermore, several other types of GABAergic interneuron in rodents, cats and primates, particularly those expressing VIP and/or CCK, and/or CR (Freund et al., 1986; Condé et al., 1994; Kawaguchi & Kubota, 1996, 1997; Meskenaite, 1997; Caputi et al., 2009), and some basket cells (Somogyi et al., 1983; DeFelipe et al., 1986; Somogyi & Soltész, 1986; Kisvárday, 1992; Jiang et al., 2015) also have descending axon collaterals crossing several layers without forming a narrow bundle. In conclusion, the available evidence suggests that CB-positive DBCs are a primate specialisation, and their preference for dendritic spine innervation (see below) is shared by homologous cell types in carnivores.

Parvalbumin-expressing interneurons include axo-axonic and basket cells in rats (Kawaguchi & Kubota, 1998; Hartwich et al., 2009) and primates (Zaitsev et al., 2005) including humans (Szegedi et al., 2020). The term ‘basket cell’ is often used loosely for PV-expressing multipolar cells without information if the cells so named form perisomatic baskets or terminate on somata at all. In all species, there is a high density of PV-positive synaptic boutons on cell bodies (Blümcke et al., 1991; Czeiger & White, 1997; DeFelipe, 1997), but they may be provided only by certain types of PV-expressing interneuron, the genuine basket cells. In the human cerebral cortex, such soma-preferring cells have not been visualised individually with their axons. The PV-DTCs recorded and visualised here targeted dendritic shafts and dendritic spines on proximal, as well as on distal dendrites with the majority of their synapses, whereas somata constituted only a small fraction of their targets. Similar dendrite-targeting GABAergic cells were also described in the frontal cortex of rats, some immunoreactive for PV (Kawaguchi & Kubota, 1998), others for CB (Hartwich et al., 2009). Large, molecularly uncharacterised dendrite-targeting cells with similar synaptic target distribution were also reported in the visual cortex of cats (Somogyi & Soltész, 1986; Tamás et al., 1997a), and shown to evoke different synaptic responses from those of synaptically defined basket cells (Tamás et al., 1997a).

The differential expression of CB and PV in DBCs and PV-DTCs, respectively, correlated with differences in their intrinsic firing properties, as described for neocortical GABAergic neurons in other species (Kawaguchi & Kubota, 1993; Kawaguchi & Kubota, 1996; Cauli et al., 1997; Cauli et al., 2000; Zaitsev et al., 2005). The most common features of DBC voltage responses were the large amplitude voltage sag in response to hyperpolarising current injection and spike amplitude accommodation in response to continuous depolarising current injection. The amplitude of the voltage sag in DBCs is comparable to that of rosehip cells in humans (Boldog et al., 2018; Field et al., 2021), however other yet undefined interneuron types, such as some expressing CB1 receptor along their axonal membrane, may also show similarly large voltage sag (Field et al., 2021). Surprisingly, neither neurons with radial axons expressing CB in the prefrontal cortex of monkeys (Zaitsev et al., 2005), nor neurons with radial axons recorded in the prefrontal cortex of rats named DBCs (Jiang et al., 2015) show these voltage patterns. This raises doubts about the homology between neurons called DBCs in different species. In addition to these features, some human DBCs also showed burst firing. In the rat prefrontal cortex, burst firing is most common in interneurons expressing neuropeptides such as VIP, CCK and SM, including neurons in layers II and III, named DBCs based on their descending axonal arbours expressing VIP and\or CRF and\or CR (Kawaguchi & Kubota, 1996; Karube et al., 2004) and Martinotti cells in layer V with ascending axonal arbours co-expressing SM and CB (Kawaguchi & Kubota, 1996). Conversely, PV-DTCs had similar voltage sag and spike amplitude accommodation to PV-expressing fast-spiking neurons described previously in humans (Szegedi et al., 2017) and in other species (Kawaguchi, 1995; Zaitsev et al., 2005; Povysheva et al., 2013). Importantly however, there was no difference in the action potential duration of DBCs and PV-DTCs, and both were shorter than that of regular-firing interneurons and longer than typical fast-spiking interneurons in other species (Kawaguchi, 1995; Cauli et al., 1997; Krimer et al., 2005).

### Dendritic spines as GABAergic synaptic targets – contributions of DBCs and other interneurons

We have confirmed that dendritic spines are the targets of the majority of synapses in the neocortex (Rakic et al., 1986; Adams, 1987; Beaulieu et al., 1992; Peters et al., 2008; Kasthuri et al., 2015; Domínguez-Álvaro et al., 2019; Cano-Astorga et al., 2021), and most of these are glutamatergic as predicted by the type I postsynaptic specialisation. More than a quarter (26%) of such presumed glutamatergic synapses are on dendritic shafts of both pyramidal neurons and interneurons in humans. Only a small proportion of dendritic spines (∼ 1.5 - 6%) are double innervated also by proven or presumed GABAergic synapses (Beaulieu et al., 1992; Kubota et al., 2007; Domínguez-Álvaro et al., 2018; Kwon et al., 2019; Cano-Astorga et al., 2021). Imaging studies of GABAergic synapses along a single pyramidal cell dendrite suggest a higher density and proportion of double innervated spines on distal than on proximal dendritic segments in mice (Chen et al., 2012). However, this might depend on the identity of the postsynaptic pyramidal cell.

The preferential innervation of dendritic spines by human DBCs confirms similar synaptic target selection in the cat (Tamás et al., 1997a; Tamás et al., 1998) and monkey (Somogyi & Cowey, 1981). There is evidence that an individual DBC innervates both the dendritic spines (n=6) and shafts (n=4) of a given pyramidal cell in the same proportion as its overall postsynaptic target distribution in layer III (Tamás et al., 1997a), and both distal and proximal dendrites were innervated. The same is true for *dendrite-targeting cells* (Tamás et al., 1997a), which in addition to dendritic spines and shafts also innervate somata. We have also found evidence that multiple spines on the same pyramidal dendrite may be targeted by the same DBC, providing a focal input. The electron microscopic examination of CB+ axonal bundles in humans and monkeys (DeFelipe et al., 1989; DeFelipe et al., 1990; DeFelipe & Jones, 1992; del Río & DeFelipe, 1995; Peters & Sethares, 1997) also suggested that spines are extensively innervated, but the contribution of individual DBCs to the bundles, or any bias in their target selection could not be established. There is some variability in the proportion of postsynaptic dendritic spine targets in the cat (38 and 69%), monkey (40%) and amongst the CB+ human DBCs (48%) studied here. Furthermore, three other Golgi impregnated interneurons with descending “horsetail” axons in cat visual cortex made much lower proportion of synapses on dendritic spines, the majority (∼86%) of synaptic targets being dendritic shafts (Somogyi and Cowey 1981), similar to the CR+ cell reported here (83% dendritic shafts). In the absence of molecular data on these cells, it is not clear if they were homologous to the DBCs in the monkey and humans. Although they were named DBCs based on their axonal patterns, these neurons may have been interneuron target preferring cell types, as a significant proportion of their synapses were found innervating non-pyramidal neuron somata and dendrites (Somogyi & Cowey, 1981). Calretinin-expressing cells in other species frequently target interneurons (del Río & DeFelipe, 1997c; Gabbott et al., 1997; Meskenaite, 1997; Gonchar & Burkhalter, 1999; Caputi et al., 2009).

The PV-DTCs studied here had similar synaptic target distribution to that of *dendrite-targeting cells* (Somogyi & Soltész, 1986; Tamás et al., 1997a) in the visual cortex of the cat, and to some fast-spiking PV-expressing interneurons in the frontal cortex of rats (Kawaguchi & Kubota, 1998; Kubota et al., 2007). Others have also reported CB-expressing dendrite-targeting cells in the rat frontal cortex (Hartwich et al., 2009). Although the proportion of dendritic spine synaptic targets of PV-DTCs was significantly lower (∼20%) than that of DBCs, and not different from the average proportion of GABAergic synapse targets in the human cortical neuropil, they may still make up significant proportion of spine innervation depending on the abundance of this cell type, which is not known. Other GABAergic neuronal types, including Martinotti cells, neurogliaform cells and basket cells, are also expected to contribute to dendritic spine innervation (Somogyi et al., 1983; Somogyi & Soltész, 1986; Meskenaite, 1997; Tamás et al., 1997a; Tamás et al., 1997b; Kawaguchi & Kubota, 1998; Tamás et al., 2003; Kubota et al., 2007; Chiu et al., 2013). However, how dendritic spines receiving glutamatergic input from different sources are selected by GABAergic cell types remains unknown.

### Multiple, location-dependent effects of GABA in cortical circuits

The differential placement of GABAergic synapses along the postsynaptic pyramidal cell surface by DBCs and PV-DTCs in the human neocortex is most likely driven by a temporal division of labour between distinct types of GABAergic neuron, as most apparent in the hippocampus (Somogyi et al., 2014), but also evident in the neocortex (Somogyi et al., 1998; Hartwich et al., 2009; Massi et al., 2012; Averkin et al., 2016). For example, the GABAergic innervation of principal cells, which include e.g. pyramidal cells, spiny stellate cells and dentate gyrus granule cells, along their axon initial segments depolarises spike initiation voltage threshold, which is best achieved by concentrating all the synapses by axo-axonic cells to the action potential initiation site. In contrast, the independent somatic GABAergic innervation decreases input resistance, shunting excitatory inputs (Rojas et al., 2011). This latter inhibitory function is best achieved by GABA distributed to multiple sites not only on the soma but also to the dendrites and even dendritic spines, as we have found for PV-DTCs and has been documented also for various basket cells (Kisvárday et al., 1985; Somogyi & Soltész, 1986; Kawaguchi & Kubota, 1998). Furthermore, perisomatic GABAergic innervation is well suited for controlling the timing of postsynaptic firing in rodents (Cobb et al., 1995; Miles et al., 1996; Tamás et al., 2000; Tamás et al., 2004) and contributes to high frequency network oscillations *in vivo* (Cardin et al., 2009). Conversely, experimental data and modelling indicate that dendrite-targeting GABAergic synapses modulate integration in distinct dendritic branches (Miles et al., 1996; Bloss et al., 2016), as has been shown for the rosehip cells in humans, which innervate the apical tufts of pyramidal cells in layer I (Boldog et al., 2018). On dendritic spines, GABAergic synapses should have an even more local effect on the glutamatergic terminal to the same spine. Indeed, these can selectively shunt penetration of somatically evoked Ca^2+^ signals into single dendritic spines and modulate local synaptic plasticity (Chiu et al., 2013). Soma- or dendrite-targeting interneuron types are differentially recruited by local pyramidal cells in both mouse and human neocortex (Obermayer et al., 2018). Nevertheless, GABAergic inputs to dendrites can also have a strong influence on postsynaptic firing through gating of dendritic spikes (Larkum et al., 1999; Lovett-Barron et al., 2012), and entraining firing rhythms at various frequencies (Szabadics et al., 2001; Tamás et al., 2004). The centripetally spreading shunting effect of multiple GABAergic synapses located on dendrites even distally to excitatory inputs may lead to inhibition of action potential generation (Gidon & Segev, 2012).

The specific spatial distribution and target neuron selectivity, together with the intricate compartmentalised placement of GABAergic synapses on the surface of single neurons can also lead to complex interactions with other inputs in the cortex (Varga et al., 2012; Somogyi et al., 2014). In this context, modelling suggests that GABAergic inputs on distal dendrites, where some of the DBC and PV-DTC synapses are located, may have an excitatory effect when arriving ahead in time to more proximally located glutamatergic input, even if E_GABA-A_ is negative to the action potential threshold (Lombardi et al., 2021).

Spatial compartmentalisation of molecular components involved in GABAergic signalling may further differentiate synaptic effects of individual presynaptic terminals. For example, a major determinant of intracellular Cl^-^ is the potassium chloride co-transporter, KCC2 that reduces intracellular Cl^-^. Interestingly, it is expressed at relatively high density in hippocampal pyramidal cell dendritic spines, particularly at the periphery of glutamatergic synapses (Gulyás et al., 2001), although its concentration along dendritic shaft membranes is higher (Báldi et al., 2010). Due to its diffusional isolation by the thin dendritic spine neck, even low expression of KCC2 would likely results in a negative E_GABA_ in dendritic spines and a hyperpolarising effect of fast GABA_A_ receptor signalling facilitated by the relatively small volume-to-surface ratio of spines. However, whether KCC2 expression is similar in human neocortical pyramidal neurons and in dendritic spines double innervated by GABAergic and glutamatergic synapses, is not known. In any case, the GABAergic synapse made by the DBC is likely to have a major role on the plasticity of the glutamatergic synapse on the same spine and is well suited for an effective postsynaptic shunting or blocking of NMDA receptor-mediated currents and Ca^2+^-dependent synaptic plasticity (Chiu et al., 2013), as well as for a presynaptic inhibition of glutamate release through GABA-B receptors.

We have shown that DBCs and PV-DTCs are GABAergic, but have not directly tested whether they were inhibitory. The postsynaptic effects of DBCs in unitary interactions seen here were consistent with fast phasic synaptic signalling mediated by GABA_A_ receptors, as shown previously in the cat (Tamás et al., 1997a). Phasic signalling involves fast and transient conductances through activated GABA_A_ receptors concentrated in the postsynaptic specialisation in response to synaptic release of GABA (Mody et al., 1994; Nusser et al., 1997; Mozrzymas, 2004; Farrant & Kaila, 2007). The effect of synaptic GABA_A_ receptor activation on the postsynaptic membrane potential would depend on the reversal potential of GABA_A_ (E_GABA-A_) determined primarily by the local transmembrane gradient of Cl^-^ (Farrant & Kaila, 2007), as well as on the timing of other inputs under physiological conditions. The DBC-evoked IPSCs recorded at the soma of postsynaptic neurons were outward currents, corresponding to a hyperpolarising effect locally. The CB+ DBCs mainly innervated pyramidal cells, but one CR+ DBC mainly targeted interneuron dendrites. Selective inhibition of GABAergic neurons could lead to disinhibition of their target neurons (Pi et al., 2013). Phasic signalling through GABA_A_ receptors may inhibit postsynaptic excitability and firing both by hyperpolarization and/or effectively shunting the inward currents generated by glutamate receptor activation particularly the slower NMDA receptor-mediated component (Dingledine et al., 1986; Collingridge & Bliss, 1987; Staley & Mody, 1992). Interacting with other time and voltage-dependent conductances, GABAergic phasic signalling may also change the spike timing of postsynaptic neurons without actually ‘inhibiting’ the firing rate (Cobb et al., 1995).

Further selective effects could emerge from the differential allocation of synaptic and extrasynaptic GABA_A_ receptor species (Farrant & Nusser, 2005; Field et al., 2021), as well as from the location of pre- and postsynaptic GABA_B_ receptors. Some GABAergic interneuron types, such as neurogliaform cells, have been shown to evoke both slow phasic and tonic GABA_A_ receptor signalling through extrasynaptic GABA_A_ and GABA_B_ receptors (Tamás et al., 2003; Oláh et al., 2007; Oláh et al., 2009). The tight axonal bundles of DBCs form an exceptionally high volume density of GABA-releasing boutons to which several individual cells contribute. If these neurons fired together, which may be facilitated by their mutual synaptic interactions as we show here, they would increase GABA concentration locally in a column that could lead to the activation of tonic currents and GABA-B receptors in dendrites and axons passing through such a column. Such a mechanism is unlikely to be involved in the action of PV-DTCs whose boutons are widely spread across several hundred microns of cortical tissue.

### Regulation of GABAergic inputs to DBCs and PV-DTCs by group III mGluRs

The GABAergic inputs of the two cell types were differentially affected by group III mGluR activation by LAP4. The spontaneous IPSCs in some PV-DTCs were suppressed as predicted from recordings in rodents, as group III mGluRs inhibit both evoked and spontaneous GABAergic synaptic transmission *in vitro* (Bonci, 1997; Schrader & Tasker, 1997; Mitchell & Silver, 2000; Wittmann et al., 2001; Matsui & Kita, 2003; Turner & Salt, 2003; Valenti et al., 2003; Kogo et al., 2004; Panatier et al., 2004; Rusakov et al., 2004; Chu & Moenter, 2005; Giustizieri et al., 2005; Cuomo et al., 2009; Summa et al., 2013) and *in vivo* (Salt & Eaton, 1995; Marabese et al., 2005; Naie & Manahan-Vaughan, 2005; MacInnes & Duty, 2008). Surprisingly, spontaneous GABAergic input to DBCs were enhanced. This may not be a direct effect on their monosynaptic input synapses, as voltage-gated Na channels were not blocked in our experiments, thus the recorded sIPSCs represented both action potential-dependent, and action potential-independent GABA release from presynaptic terminals in the network. Furthermore, glutamatergic synaptic transmission was intact, thus, group III mGluR action at glutamatergic synapses may have also contributed to the observed effects. Thus, the effects were not restricted to monosynaptic inputs. Facilitation of sEPSCs by group III mGluR activation was reported in layer V neurons in the entorhinal cortex (Evans et al., 2000; Woodhall et al., 2007) and the amygdala (Ren et al., 2011) of rats *in vitro*, under conditions similar to ours with intact GABAergic neurotransmission. In our experiments the changes in frequency of sIPSCs were not accompanied by changes in amplitude in either cell type, indicating a presynaptic site of action, consistent with the presynaptic location of group III mGluRs (Shigemoto et al., 1996; Shigemoto et al., 1997; Dalezios et al., 2002; Somogyi et al., 2003). The most likely explanation for the observed facilitation of sIPSCs is a selective disinhibition of a specific GABAergic input to DBCs through network effect. Such a mechanism may also explain observations of apparent facilitation of neurotransmitter release by group III mGluRs, as measured by microdialysis in other species (Marabese et al., 2005; Li et al., 2008).

The sources of GABAergic innervation of CB+ DBCs are not known. Immunohistochemical studies have shown that CB+ neurons in the monkey visual and temporal cortex receive synaptic input from CR+ neurons and one third of them receive synaptic inputs from PV-positive neurons (DeFelipe et al., 1999). Because GABAergic inputs to PV-DTCs in humans are suppressed by group III mGluR activation, which may lead to their disinhibition, they could contribute to the increased GABAergic synaptic input to DBCs. The GABAergic inputs to PV-DTCs may include VIP-expressing neurons, a subset of which are known to selectively innervate other GABAergic neurons (Acsády et al., 1996), including those that express PV in the neocortex of rodents (Dávid et al., 2007; Hioki et al., 2013). Consistent with this hypothesis, VIP-expressing neurons in non-human species have been shown to express high levels of group III mGluRs, including both the high affinity subtype, mGluR8, as well as the low affinity mGluR7 (Dalezios et al., 2002; Katona et al., 2020). Multiple datasets of single cell RNA sequencing in human neocortex also shows selective expression of mGluR8 in some VIP-, SM- and PV-expressing GABAergic transcriptomic cell types (https://portal.brain-map.org/atlases-and-data/rnaseq).

### Potential roles of DBCs in the neocortex

The physiological activity of DBCs *in vivo* is not known, but they form a significant radial GABAergic axon system (Somogyi et al., 1981). Any hypothesis for their roles must take into account two unique features of CB+ DBCs, namely, the striking radial “horsetail” axon and the high proportion of dendritic spine output synapses. The regularly distributed CB+ or “horsetail” axon bundles found ∼ 15-30 *μ*m apart horizontally (DeFelipe et al., 1990; Peters & Sethares, 1997; Ballesteros Yáñez et al., 2005) and spanning layers III-V in primates, including humans, highlights the modular (Szentágothai, 1975) and columnar organisation in the neocortex (Hubel & Wiesel, 1977; Mountcastle, 1997). The radial GABAergic CB+ axon bundles appear to run together with bundles of myelinated axons (DeFelipe et al., 1989; Peters & Sethares, 1997) adjacent to radially aligned pyramidal cells forming ‘minicolumns’ (Mountcastle, 1997) as well as the radial apical dendrite bundles (Peters & Sethares, 1996; Peters & Sethares, 1997) through multiple layers. The myelinated axon bundles include, but are not restricted to, the axons of some of the pyramidal cells (Peters & Sethares, 1996), and some other pyramidal cells in layers II-III, which have unmyelinated axons making numerous glutamatergic synapses as they descend through the layers. The fine-grained spatial relationships of these components however are not yet clear. In addition, we have shown that multiple individual DBCs, including CB+, CB- and CR+ cells, with diverse firing patterns and dendritic structure contribute to radial axonal bundles, and further components remain to be explored. All electron microscopic studies agree (Somogyi & Cowey, 1981; DeFelipe et al., 1989; DeFelipe et al., 1990; Peters & Sethares, 1997) that apical dendritic shafts are not targeted by DBCs, but it cannot be excluded that spines originating from them are innervated. Most dendritic spines and dendritic shafts postsynaptic to DBC GABAergic terminals are probably basal dendrites and/or the oblique dendrites originating from apical dendrites. We hypothesise that the radial beam of high-density GABA release sites from DBCs follows and converges on the same postsynaptic targets as the radial beam of descending pyramidal cell axon terminals. Thus, DBCs are in a position of modulating the effect of their glutamatergic pyramidal cell synapses. This modulation may be inhibitory through both pre- and postsynaptic effects as well as tonic inhibition through extrasynaptic GABA receptors. In turn the glutamate released by the pyramidal cell terminals may inhibit GABA release through presynaptic mGluRs and assist synaptic plasticity of the glutamatergic synapse (Klar et al., 2015). Several DBC axonal bundles (7 - 45) would cross the dendritic field of any given radial pyramidal cell column, segmenting the dendritic field and forming a ‘frame’ in which the different bundles may carry information about different horizontal locations in the representational space. If the postsynaptic basal and oblique dendrites of different pyramidal cells in the radial column were aligned horizontally at similar angles as viewed from the pia, the DBC axon bundles running through them would shape the responsiveness of the subset of dendrites at similar angles within the column. The preferential innervation of dendritic spines by CB+ DBCs suggests that they regulate glutamatergic synaptic plasticity of some inputs, one of which, as we suggest above, is from the column of pyramidal cells with cell bodies next to the axon bundle. Modelling hypothetical DBC effect on synaptic plasticity suggests that the result would be highly localised to the innervated dendritic column (Bar-Ilan et al., 2013). The CR+ DBCs may modulate signal processing in interneuron dendrites passing through the same radial column. In sensory cortical areas dendritic spines receiving co-tuned synaptic input are dispersed across the dendritic tree of pyramidal cells, (Jia et al., 2010; Chen et al., 2011), although the clustering of functionally related inputs on neighbouring spines at a fine spatial scale can be implemented (Scholl et al., 2017). The “horsetail” axons of DBCs are particularly well suited to modulate synaptic inputs on neighbouring spines along restricted dendritic segments of multiple radially aligned pyramidal cells. The enhanced inhibition of DBCs by the action of presynaptic group III mGluRs in the network that we observed could lead to select disinhibition of innervated dendritic spines on different dendritic segments of a pyramidal cell and provide a mechanism for the functional clustering of neighbouring dendritic spines, contributing to their stimulus selectivity (Wilson et al., 2016).

## Acknowledgements

We are grateful to Professor Colin Akerman at the Department of Pharmacology, Professor Zameel Cader and Dr Tatjana Lalic at the Weatherall Institute of Molecular Medicine, Professors John Morris Helen Christian, Andrew King and Victoria Bajo Lorenzana at the Department of Anatomy, Physiology and Genetics, all at the University of Oxford, and Professor Zoltán Nusser and Dr Andrea Lőrincz at the Institute of Experimental Medicine Hungarian Academy of Sciences, Budapest, Hungary, as well as to members of their laboratories, for allowing us to use equipment for sample preparation, electrophysiological recording, electron microscopy and Neurolucida digital reconstruction. We also thank Ms Hannah Brooks and Ms Carolyn Sloan at the Oxford Brain Bank for their support with sample administration and the many members of the surgical teams who assisted during the brain surgeries. We also thank Dr József Somogyi for advice on confocal microscopy.

## Author’s contributions

I.P.L ., R.F., M.F., E.H., M.H., S. H. and P. S. designed experiments, collected and analysed data; P. P., R. S., L. L., O. A. and G. T. provided samples, clinical and management services; all authors edited and approved the manuscript.

## Footnotes

1 http://www3.mpibpc.mpg.de/groups/neher/index.php?page=aboutppt

