## supplementary figure 1. for "Differential effects of group III metabotropic glutamate receptors on spontaneous inhibitory synaptic currents in spine-innervating double bouquet and parvalbumin-expressing dendrite-targeting GABAergic interneurons in human neocortex"

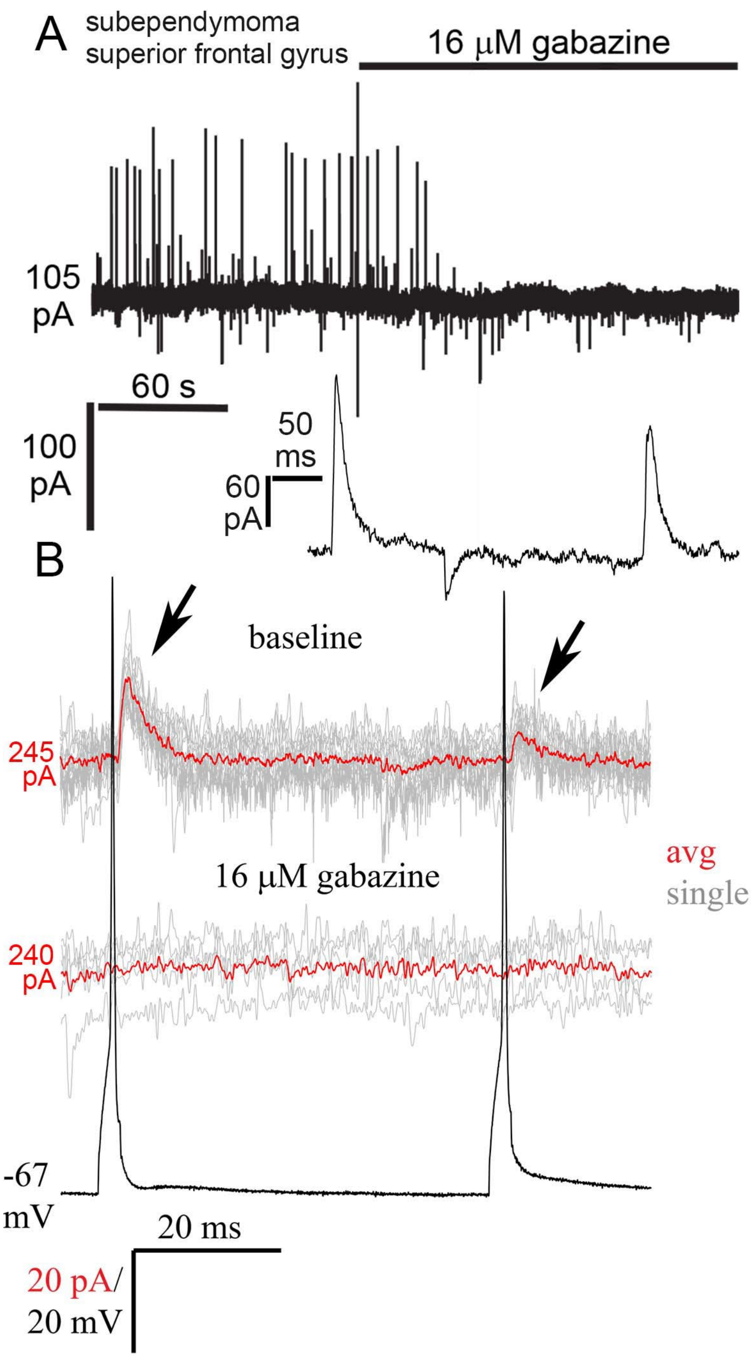


**Supplementary Figure 1. Spontaneous and evoked outward currents are IPSCs and mediated by $GABA_A$ receptors. A.** Continuous voltage-clamp recording of a visualised interneuron at -50 mV membrane potential and $E_{Cl}$ = -94 mV. The holding current is indicated. Outward currents, represented by upward spikes, correspond to sIPSCs, whereas inward currents (downward spikes) represent sEPSCs. Bath application of 16 mM gabazine, a $GABA_A$

receptor antagonist, blocks sIPSCs, but not sEPSCs. Inset shows a period before application of gabazine with expanded time axis to illustrate the kinetics of the synaptic currents. The interneuron was immunoreactive to CB1 along its axonal membrane and to VGlut3 in its boutons and CCK in its soma. **B.** Two action potentials evoked in a visualised bitufted interneuron (black trace) recorded in current-clamp mode, elicited monosynaptic eIPSCs (arrows, top trace: red – avg. of 15 sweeps) in another visualised multipolar interneuron, recorded in voltage-clamp mode, at -50 mV. Bath application of 16 μM gabazine completely abolished eIPSCs in the postsynaptic neuron (middle trace: red – avg. of 5 sweeps). The holding current in the postsynaptic neuron and the baseline membrane potential of the presynaptic neuron are indicated. Recordings are from superior frontal gyrus, pathology: subependyoma.
